# Small Extracellular Vesicles from Failing Heart Accelerate Tumor Growth

**DOI:** 10.1101/2023.09.03.555686

**Authors:** Tal Caller, Itai Rotem, Olga Shaihov–Teper, Daria Lendengolts, Yeshai Schary, Ruty Shai, Efrat Glick-Saar, Dan Dominissini, Menachem Motie, Idan Katzir, Rachela Popovtzer, Merav Nahmoud, Alex Boomgarden, Crislyn D’Souza-Schorey, Nili Naftali-Shani, Jonathan Leor

## Abstract

**Background:** Myocardial infarction (MI) and heart failure (HF) are associated with an increased incidence of cancer. The mechanism is complex and unclear. Here, we aimed to test our hypothesis that cardiac small extracellular vesicles (sEVs), particularly cardiac mesenchymal stromal cells-derived sEVs (cMSC-sEVs), contribute to the link between post-MI HF and cancer.

**Methods:** We purified and characterized sEVs from the whole heart and cultured cMSCs. Then, we analyzed cMSC-EV cargo and pro-neoplastic effects on several types of cancer cell lines, macrophages, and endothelial cells. Next, we modeled post-MI HF along with heterotopic and orthotopic lung and breast cancer tumors in mice. We used cMSC-sEV transfer to assess sEV biodistribution and its effect on tumor growth. Finally, we tested the effects of sEV depletion and spironolactone treatment on cMSC-EV release and tumor growth.

**Results:** Post-MI hearts, particularly cMSCs, produced more sEVs with pro-neoplastic cargo than non-failing hearts did. Proteomic analysis revealed unique protein profiles and higher quantities of tumor-promoting cytokines, proteins, and microRNAs in cMSC-sEVs from failing hearts. The pro-neoplastic effects of cMSC-sEVs varied with different types of cancer cells, substantially affecting lung cancer cells relative to other more aggressive cancer cell lines. We also found that post-MI cMSC-sEVs activated resting macrophages into pro-angiogenic and pro-tumorigenic states in vitro. At 28-day follow-up analysis, mice with post-MI HF developed larger lung tumors than did sham-MI mice. Adoptive transfer of cMSC-sEVs from failing hearts accelerated lung tumor growth, and biodistribution analysis revealed an accumulating cMSC- sEVs in tumor cells along with accelerated tumor cell proliferation. Significantly, sEV depletion reduced the tumor-promoting effects of HF, and adoptive transfer of cMSC-sEVs from failing hearts partially restored it. Finally, post-MI spironolactone treatment reduced the number of cMSC-sEVs and suppressed tumor growth during post-MI HF.

**Conclusions:** For the first time, we show that cardiac sEVs, specifically cMSC-sEVs from post-MI failing hearts, carry multiple pro-tumorigenic factors. Uptake of cMSC-sEVs by cancer cells accelerates tumor growth. Post-MI spironolactone treatment reduces the associated tumor growth. Thus, we provide new insight into the link between post-MI HF and cancer and propose a translational option to mitigate this deadly association.

---

Cancer and heart failure (HF) are common diseases with shared risk factors, mechanistic similarities, and lethal interactions.^1-4^ While anti-cancer therapies accelerate HF, the presence of HF may increase cancer incidence.^1,5-7^ The coexistence of these diseases worsens patients’ prognoses and negatively affects therapeutic options.^5,7,8^

The association between HF and cancer is complex and not entirely clear. It is partially attributed to shared risk factors.^1,2^ However, recent studies have suggested a more causal explanation: circulating factors released from injured hearts accelerate tumor growth, independent of risk factors.^9-13^ However, the link between HF and cancer is not fully understood.^1,2^

A potential unexplored mechanism that may link heart disease to cancer is small extracellular vesicles (sEVs). sEVs (<200 nm) have emerged as a new and powerful mechanism of communication between cells and their environment through sEVs’ ability to transmit multi-molecular biological messages with much greater complexity than a single factor.^14-17^ sEVs exert their function by altering recipient cells via delivery of RNA, cytokines, chemokines, growth factors, or surface protein signaling ^14-17^ sEVs play a role in disease processes like heart disease and cancer.^15,16^ It has been suggested that sEVs mediate the communication between the heart and other organs.^18^ However, whether and how sEVs from the failing heart promote tumor growth remains unknown.

Given the central role of sEVs in cell-to-cell communication, we hypothesized that cardiac sEVs contribute to the association between post-MI HF and tumor growth. Here, we tested our hypothesis in a mouse model of post-MI HF and cancer. Understanding how post-MI HF may promote cancer could help develop treatments simultaneously targeting both diseases. Additionally, understanding the mechanism of HF and cancer association could help identify patients with cardiovascular disease at higher risk of cancer and may allow for early risk stratification, prevention, and treatment.

## METHODS

Detailed methods are provided in the Supplemental Material. All data that support our findings are available within the article, the Data Supplement, and are available upon reasonable request. Animal experiments were conducted following the National Institutes of Health (NIH) Guide for the Care and Use of Laboratory Animals and complied with the standards and approval of the Sheba Medical Center Institutional Animal Care and Use Committee.

We purified and characterized sEVs from the whole heart and isolated cMSCs. We analyzed cMSC-sEV size distribution, markers, cargo, and effects on cancer cells, endothelial cells (ECs), macrophage activation, and tumor growth. We modeled post-MI HF in mice, along with heterotopic and orthotopic lung and breast cancer tumors. To monitor left ventricular (LV) remodeling and dysfunction, we used a small animal ultrasound (US) system. To assess tumor growth, we used US, bioluminescent IVIS Spectrum imaging, and micro-CT. We also used cMSC-EV transfer to assess sEV biodistribution, uptake, and effects on tumor growth. Finally, we tested the effects of EV depletion and spironolactone treatment on cMSC-EV release and tumor growth.

### Statistical Analysis

Data are expressed as mean± standard deviation (SD). Specific statistical tests are detailed in the figure legends. Statistical analyses were performed with GraphPad Prism v9.5.0 unless otherwise stated. Statistical analyses of the identification and quantification of the proteomic analysis (proteomics) were performed using Perseus 1.6.7.0, and statistical analysis for nanoparticle tracking analysis (NTA) data was performed using STATA BE v17.

Two-tailed Student’s t-test was performed to compare normally - distributed continuous variables and a two-tailed Mann-Whitney U test was used for non-normal distribution (tested by D’Agostino-Pearson normality test). When the experimental design included more than one comparison, we used multiple t-tests or multiple Mann-Whitney tests to account for multiple comparisons. For experiments with more than 2 groups, we used one-way or two-way ANOVA with Holm-Sidak’s post-test. Repeated measure tests were used if the same subject was sampled at different time points. To compare two frequency distributions (NTA data), we used zero-inflated negative binomial regression.

## RESULTS

### The Infarcted and Failing Heart Generated sEVs with Pro-Neoplastic Properties

To determine the role of cardiac sEVs in tumor growth, we first assessed the production of sEVs in the entire myocardium in response to post-MI HF (Fig-S1A). We subjected female C57BL/6 mice to MI and sham-MI. HF in MI mice was confirmed by left ventricular (LV) remodeling and dysfunction by echocardiography, plasma N-terminal pro-brain natriuretic peptide (NT-proBNP) (Fig-1A), and lung congestion (Fig-S2A). Our method of isolating and purifying whole-heart EVs was inspired by the work of Loyer et al.,^19^, and we purified sEVs by size exclusion chromatography (SEC) (Fig-S1B).^20,21^ To characterize sEVs, we used electron microscopy (TEM) (Fig-S1C) and western blot for the sEV markers: CD81 and tumor susceptibility gene 101 (TSG101) (Fig-S1D, Fig-S4A). Next, we used nanoparticle tracking analysis (NTA) to determine the morphology and size distribution of sEVs (Fig-S1E-F). We found that failing hearts produced more than twice the number of sEVs in sham-MI hearts (Fig-S1E-F). Overall, we found that in the early phase of post-MI HF, the LV myocardium produced more sEVs.

**Figure 1:**
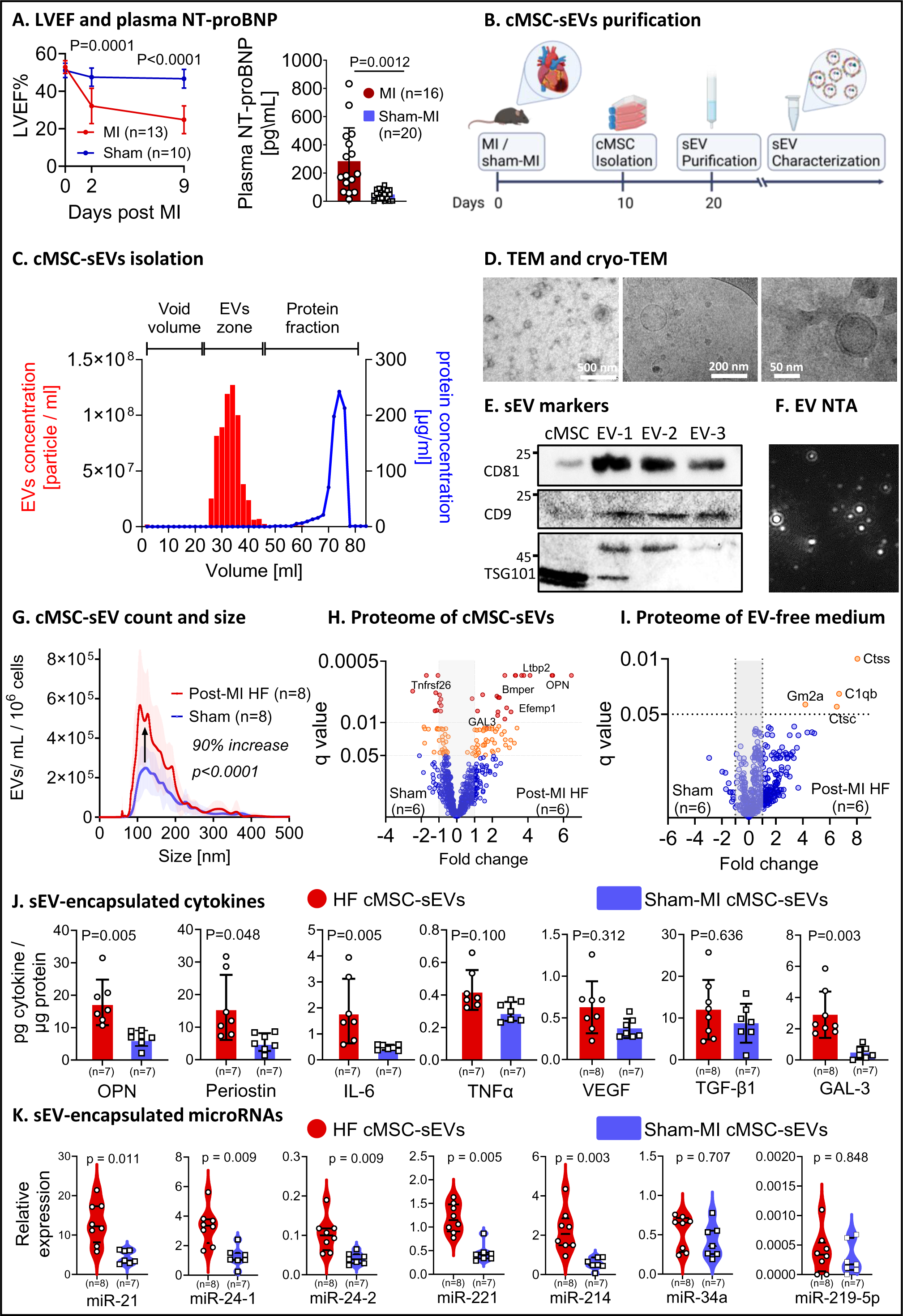
cMSC-sEVs from Post-MI Failing Hearts Harbored Tumor-Promoting Mediators. A) MI-induced LV dysfunction and plasma NT-proBNP elevation. To monitor cardiac function after MI or sham-MI, we performed serial echocardiography measurements and calculated LV ejection fraction at baseline, day 2, and day 9 post-MI. P values were calculated using two-way repeated measures ANOVA with Holm-Šídák’s post-test. Comparisons for MI vs. sham-MI are indicated on the graph. P for MI, p for time, and p for interaction all <0.0001. Next, we used ELISA to measure plasma NT-proBNP of mice 10 days post-MI or sham-MI. The data passed the normality test by D’Agostino-Pearson, and the p-value was calculated using a two-tailed unpaired t-test with Welsch correction for unequal variance. B) To isolate cMSC-sEVs, female C57BL/6 mice were randomized to either MI or sham-MI operation. At day 10, hearts were harvested, and cMSCs were isolated by enzymatic digestion and cultured in vitro. At 85% confluence, the medium was replaced by a serum-free medium for 72 hours. Then, EVs were purified from the conditioned medium using size exclusion chromatography. C) SEC successfully separated cMSC-sEVs from soluble proteins. D) Typical sEV morphology confirmed by negative staining transmission electron microscopy (TEM) (Left panel) and cryo-TEM (middle and right panels). Scale bars from left to right: 500 nm, 100 nm, and 50 nm. E) The presence of typical EV markers: CD81, CD9, and TSG101 were determined by a western blot. F-G) To determine cMSC-sEV secretion, we used nanoparticle tracking analysis (NTA). The failing hearts secreted twice more sEVs compared to sham-operated hearts. P values were calculated using zero-inflated negative binomial regression, and percent change (MI/sham-MI) was calculated using the ratios of the area under the curves. H-I) To determine the differences in protein expression between cMSC-sEVs from failing and sham-MI hearts, we carried out a comparative proteomic analysis. The volcano plot showed a statistically significant change in 114 proteins and a unique profile of cMSC-sEVs from failing hearts (H). Analysis of EV-free conditioned medium from cMSCs (I) revealed modest changes in protein profile compared with the changes observed in cMSC-sEVs. EV-free conditioned medium from cMSCs from the failing heart differentially (q<0.05) expressed 4 proteins compared with sham-MI. P values were determined by multiple, two-tailed, unpaired t-tests with the FDR method to account for multiple comparisons. J) To investigate cMSC-sEV encapsulated cytokines, we used ELISA assays. P values determined by multiple, two-tailed, unpaired Mann-Whitney U test with Holm-Šídák’s post-test to account for multiple comparisons. K) To identify miRs within cMSC-sEVs that could influence tumor progression, we used RT- PCR. Compared with sham-MI, we found that cMSC-sEVs from the failing heart encapsulated more tumor-promoting miRs, such as miR-221, that stimulate tumor development, cancer cell proliferation, and migration. Other miRs, more abundant in cMSC-sEVs from the failing hearts, include miR-21, which promotes digestive system cancers by supporting cell survival, proliferation, migration, and immunomodulation. cMSC-sEVs also harbored more miR-24-1, which is associated with ovarian cancer cell proliferation, and more; and miR24-2, which acts as a tumor suppressor or oncogenic miR, depending on the cancer type, and which promotes breast cancer by inducing cell proliferation and lung cancer by supporting angiogenesis. cMSC- sEVs from failing hearts also harbored more miR-214, which is upregulated in the lung, breast, prostate, and several other cancer types and may drive tumor development by promoting angiogenesis, cell division, invasion, and metastasis. Other tumor-promoting miRs were detected in similar amounts in cMSCs-EVs from HF and sham MI: miR-34a, associated with increased cancer cells proliferation and survival, and miR-219-5p, associated with metastasis and growth of gastric and colon cancer, but suppression of ovarian cancer. Additionally, miR- 208a, associated with tumor cell proliferation and invasion, was undetectable in cMSC-sEVs. P values by multiple, two-tailed, unpaired Mann-Whitney U test with Holm-Šídák’s post-test to account for multiple comparisons. Illustration created with BioRender.com.

Next, we assessed the pro-neoplastic properties of sEVs from the failing heart.

Compared with sham MI, cEVs from the failing heart accelerated proliferation of Lewis lung cancer (LLC) cells (Fig-S1G). The proliferative effects of cEVs followed a dose-response pattern (Fig-S1H). Moreover, sEVs from the failing heart accelerated migration of LLC cells compared with sham-MI sEVs (Fig-S1I), again, with a dose-response pattern (Fig-S1J). Collectively, post-MI failing hearts generated a high number of sEVs with pro-neoplastic properties. However, our experiment neither differentiated between cellular and extracellular vesicles nor identified the cell source of sEVs.

#### Mesenchymal Stromal Cells from Post-MI Failing Heart are a Rich Source of sEVs

The heart contains many types and sub-types of cells,^22,23^ all are potential sources of sEVs. We focused on cMSCs,^24^ because MSCs are activated during tissue injury, repair, and cancer. MSCs have pleiotropic effects, including pro-inflammatory,^25,26^ fibrotic,^27^ and tumor-promoting functions.^26,28^ Notably, activated cMSCs, particularly fibroblasts, maintain their in vivo signature through at least 4 passages and are therefore suitable for in vitro studies.^29^

First, we isolated and cultured cMSCs 10 days after MI or sham-MI, after resolution of the acute inflammatory phase (Fig-1A).^29^ Flow cytometry showed that cMSCs expressed the fibroblast markers collagen 1α (COL1α) (94%), CD90 (76%), and mEF-SK4 (98%) (Fig-S3A- C).^30^ The resident fibroblast marker, platelet-derived growth factor receptor α (PDGFRα),^30^ was downregulated after MI compared with sham-MI (36% vs. 48%, p=0.02, Fig-S3D). cMSCs did not express markers of macrophages or endothelial cells (Fig-S3E-F). Next, after 72 hours of incubation, we collected the serum-free conditioned medium. The viability of cMSCs at the time of medium collection was 90.5% and 86.7% for post-MI and sham-MI cMSCs (Fig-S3G). Then, we used SEC to isolate and purify sEVs from the cMSC-conditioned medium (Fig-1B-C).^20^

Purified cMSC-sEVs showed typical EV morphology with a double-layer membrane and normal appearance in cryogenic or negative staining TEM (Fig-1D). Western blot confirmed sEV membranous markers CD81 and CD9; and cytosolic marker TSG101 (Fig-1E and S4B). Like the entire heart, cMSCs from the failing heart secreted twice as many sEVs as did cMSCs from sham-MI (Fig-1F-G). The size distribution pattern in SEC fractions revealed that most of the sEVs were between 100 and 200 nm (Fig-1G). Overall, we showed that isolating cMSC-sEVs by SEC produces a pure and enriched population of sEVs without contaminated proteins. cMSCs from the failing heart produced more sEVs, mirroring the situation in the whole heart after MI.

#### Distinct Proteomic Profile of cMSC-sEVs from the Failing Heart

To explore the cargo of cMSC-sEVs, we conducted comparative proteomic profiling and identified 931 cMSC-EV encapsulated proteins. Comparing cMSC-sEVs from the failing heart to those from sham-MI, we found 64 upregulated and 50 downregulated proteins. (Fig-1H). For example, osteopontin, a reparative, tumor-promoting protein, was 5.4 times higher in cMSC- sEVs from the failing heart than in non-failing hearts. In addition, 174 proteins were detected exclusively in cMSC-sEVs from failing hearts, and 31 were absent from cMSC-sEVs from failing hearts (Fig-S5A). Thus, cMSC-sEVs from the failing heart presented a distinct proteomic profile.

To further characterize the proteins from cMSC-sEVs, we analyzed biological pathways using the functional enrichment tool (FunRich) of VesiclePedia using the gene ontology biological processes database.^31^ We found that cMSC-sEVs from the failing heart were enriched in pathways related to the development and progression of cancer (Fig-S5B).^32^ Pathways such as vascular endothelial cell proliferation and mitotic cell cycle phase transition were enriched exclusively in post-MI cMSC-sEVs. Overall, post-MI HF altered the proteomic profile of cMSC- sEVs toward a reparative and tumor-promoting repertoire.

Analysis of EV-free conditioned-medium from cMSCs revealed modest differences in protein profile between cMSCs from failing vs. sham-MI hearts (Fig-1I). Thus, our results suggest that the protein cargo of cMSC-sEVs better reflects the environment of failing hearts, compared with cMSC-EV-free conditioned medium.

#### cMSC-sEVs from the Failing Heart Carry Cytokines with Pro-Tumorigenic Properties

EVs encapsulate various cytokines that might contribute to neoplastic growth.^33,34^ Compared with cMSC-sEVs from sham-MI hearts, cMSC-sEVs from the failing heart carried higher amounts of cytokines involved in pathogenesis of cardiovascular diseases and cancer,^35^ including periostin (3-times); osteopontin (2.7-times), interleukin-6 (IL-6, 4-times), galectin-3 (6.2-times), tumor necrosis factor α (TNFα, 1.4-times) and vascular endothelial growth factor (VEGF, 1.7-times) (Fig-1J). Interestingly, the concentration of sEV-encapsulated galectin-3, a protein implicated in the pathogenesis of cardiovascular disease and cancer,^35^ was 3.5 times higher than that of soluble galectin-3 (Fig-1J, S6B). Post-MI HF did not affect the amount of IL- 1α, IL-1β, IFNγ, and TGF-β within cMSC-sEVs (Fig-1J, S6A-B). However, the concentration of TGF-β within cMSC-sEVs was 31 times higher than that of free TGF-β (Fig-1J, S6B). This finding is noteworthy because TGF-β has been implicated in cancer cell invasion, dissemination, and recruitment of tumor-associated macrophages (TAMs).^36,37^

In parallel, analysis of soluble cytokines in the cMSC-conditioned medium revealed higher quantities of soluble cytokines than cMSC-sEVs (Fig-S6B). However, only periostin and IL-6 maintained higher profile characterized cMSC-sEVs from the post-MI hearts. Collectively, our results indicated that cMSC-sEVs from post-MI failing hearts carried a unique tumor-promoting cytokine signature. Compared with soluble cytokines, the cargo of cMSC-sEVs better reflected the physiological processes within the infarcted and failing myocardium.

#### cMSC-sEVs Encapsulated microRNAs (miRs) with Pro-tumorigenic Properties

Pro-tumorigenic miRs are significantly enriched in sEVs.^16^ To determine whether cMSC-EVs carry tumor-promoting miRs, we first analyzed the proteomic profile of LLC tumors from mice with and without post-MI HF. On day 30 after tumor inoculation, tumor tissues were processed for proteomics. We identified 4,909 tumor proteins, of which 101 were differentially expressed in the post-MI HF group (Fig-S7A). Using TargetScan tool,^38^ we identified 48 genes-encoding proteins with a known miR binding site in their 3’-untranslated region (3’-UTR) and a total of 70 miRs that can bind to those sites. Then, to filter for miR with reported involvement in cancer, we used the miRBase tool. We filtered the list for miR involvement in cardiovascular diseases and found 17 potential tumor-promoting miRs (Fig-S7B).

We next extracted and analyzed total EVs RNA from cMSC-sEVs. Compared with sham-MI sEVs, cMSC-sEVs from the failing heart contained more tumor-promoting miRs, such as miR- 221, that stimulate tumor development, cancer cell proliferation, and migration (Fig-1K). Other miRs that were found more abundantly in cMSC-sEVs from the failing hearts included miR-21, miR-24-1, and miR-214 (Fig-1K). Other tumor-promoting miRs that were detected in similar amounts in cMSCs-EVs from post-MI HF and sham MI included: miR-34a and miR-219-5p. Overall, cMSC-sEVs from the failing heart carry higher amounts of certain tumor-promoting miRs.

#### cMSC-sEVs from Failing Heart Accelerated Proliferation and Migration of Cancer Cells

Activated MSCs may possess pro-neoplastic functions.^26,28^ To assess the pro-neoplastic properties of cMSC-sEVs, we added cMSC-sEVs (10^8^ EVs/mL) or saline to different cultured cancer cells: LLC lung cancer, MC38 colon cancer, EO771 triple-negative breast, and B16 melanoma cell lines. Colorimetric-based proliferation assay showed that cMSC-sEVs from the failing heart stimulated proliferation of LLC and MC38 cells by 1.5 and 1.2 times more than cMSC-sEVs from sham-MI hearts or saline (Fig-2A-B, left column). The proliferative effect of cMSC-sEVs corresponded to a dose-response pattern (Fig S7A-B). Overall, the effects of cMSC-sEVs were tumor-specific: significant on lung and colon cancer cells and modest or null with breast and melanoma cancer cells (Fig-2A-D, left column).

**Figure 2:**
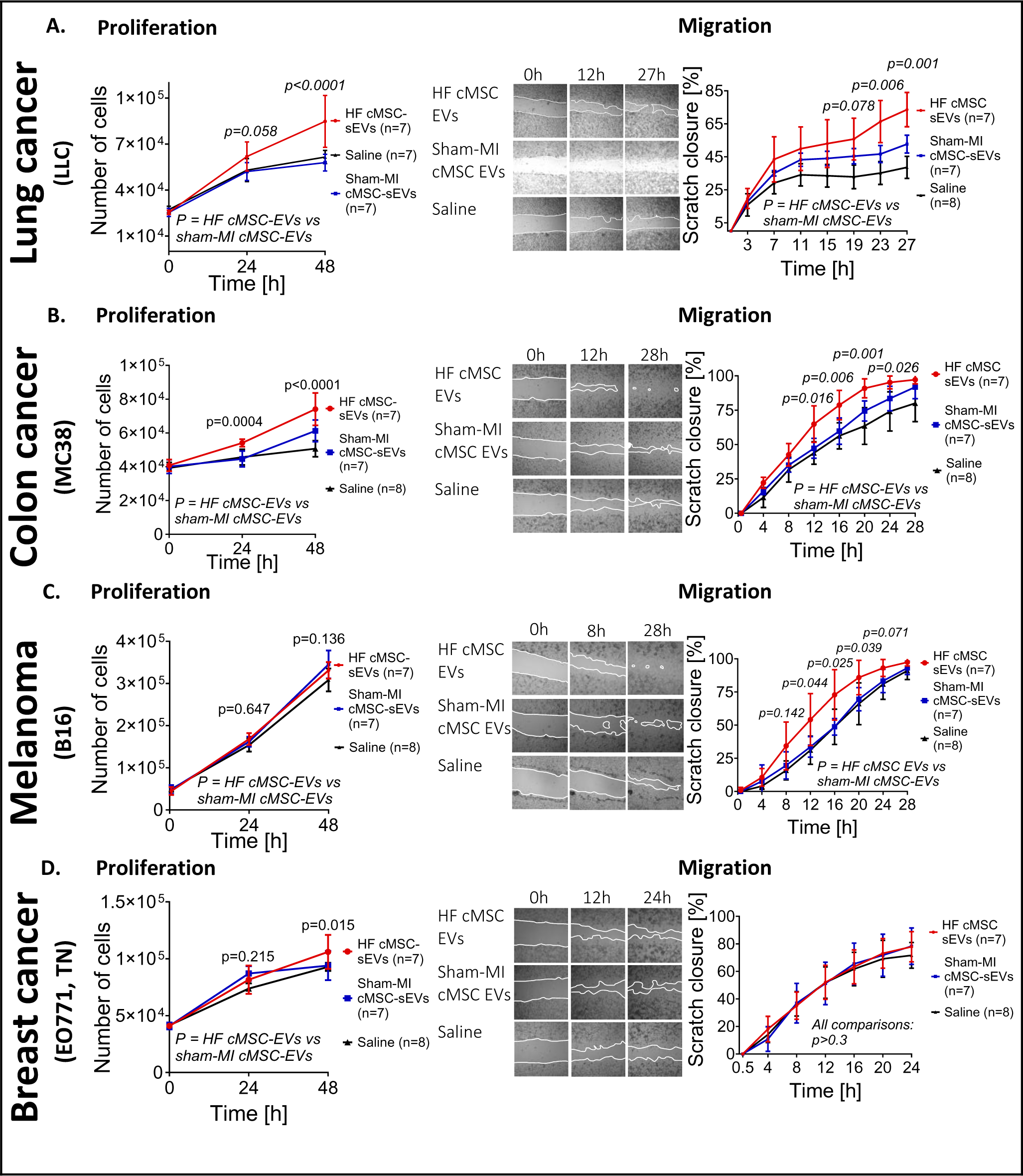
The Effects of cMSC-sEVs from Post-MI Failing Hearts on Cancer Cell Proliferation and Migration. cMSC-sEVs were purified 10 days after MI or sham-MI using SEC. The effects of cMSC-sEVs on cancer cell lines were evaluated by colorimetric cell proliferation and cell migration (scratch) assays. A) cMSC EVs (10^8^ EVs/mL) from failing hearts facilitated LLC cancer cell proliferation. P for cMSC-sEVs <0.0001, p for time < 0.0001, p for interaction = 0.0001. cMSC-sEVs from the failing hearts increased lung cancer cell migration two times compared to sham-MI cMSC-sEVs. P for cMSC-sEVs =0.0002, p for time < 0.0001, p for interaction = 0.0001. B) cMSC-sEVs (10^9^ EVs/mL) from failing hearts facilitated MC38 cancer cell proliferation. P for cMSC-sEVs <0.0001, p for time < 0.0001, p for interaction < 0.0001. cMSC-sEVs from the failing hearts increased MC38 migration compared to sham-MI cMSC-sEVs. P for cMSC-sEVs = 0.0003, p for time < 0.0001, p for interaction = 0.0001. C) cMSC-sEVs accelerated migration but not the proliferation of B16 Melanoma cells. A colorimetric proliferation assay showed that cMSC EVs (10^9^ EVs/mL) from failing hearts did not facilitate B16 cell proliferation. P for cMSC-sEVs = 0.8473, p for time < 0.0001, p for interaction = 0.6641. For migration assay: P for cMSC-sEVs =0.0059, p for time < 0.0001, p for interaction = 0.0001. All samples were assayed in duplicates. D) The effect of cMSC-sEVs (10^9^ EVs/mL) on proliferation (modest) and migration (no effect) of EO771 triple-negative breast cancer cells: P for cMSC-sEVs =0.0211, p for time < 0.0001, p for interaction = 0.0164; and scratch (migration) assay with 10^9^ EVs/mL. P for cMSC-sEVs =9044, p for time < 0.0001, p for interaction = 0.1600. All samples were assayed in duplicates. P values were calculated using two-way ANOVA with Holm-Šídák’s post-test for proliferation assays and P two-way repeated measures ANOVA with Holm-Šídák’s post-test for migration assays. P for HF cMSC-sEVs vs. sham-MI cMSC-sEVs is indicated on the graph. All samples were assayed in duplicates.

Next, we assessed the effect of cMSC-sEVs on migration of cancer cells. We used a scratch assay with several types of cancer cells (Fig-2, middle column). Administration of cMSC-sEVs (10^7^ EVs/mL) from the failing heart accelerated migration of LLC cells compared with cMSC-sEVs from sham-MI or saline (Fig-2A, right column). The magnitude of the effect increased as the concentration of EVs increased (Fig-S8C-D). Surprisingly, the migratory effect on LLC cells decreased with the highest concentration of cMSC-sEVs. Next, we found that cMSC-sEVs from the failing heart accelerated migration of MC38 (colon) and B16 (melanoma) but not EO771 (breast) cancer cell lines (Fig-2B-D, right column). Overall, aggressive cancer cells (e.g., melanoma and breast) were less affected by cMSC-sEVs.

Finally, ECs have been implicated in the development and progression of tumors.^32^ We found that cMSC-sEVs from the failing heart (10^9^ EVs/mL) promoted C166 endothelial cell permeability and migration (Fig-S8E-F).

#### cMSC-sEVs Activated Macrophages into a Pro-Tumorigenic Phenotype

Monocytes and macrophages, whose functions could be regulated by sEVs,^19^ have both tumor-promoting and tumor-suppressive effects.^36,39^ To determine the effect of cMSC-sEVs on macrophage activation and function, we isolated peritoneal macrophages from naive mice (96% purity, Fig-3A).^40^ Adding cMSC-sEVs from post-MI failing heart, sham-operated heart, or saline, we found that cMSC-EVs from the failing heart upregulated the expression of inducible nitric oxide synthase (iNOS) (Fig-3B, top). Moreover, concentrations of nitrite and nitrate, indirect products of iNOS, were higher in the conditioned medium of macrophages treated with sEVs from the failing heart (Fig-3B, bottom). These findings are significant because TAMs are characterized by producing NO and reactive oxygen intermediates, which can cause DNA damage and genetic instability during the initiation phase of tumors.^36^

**Figure 3:**
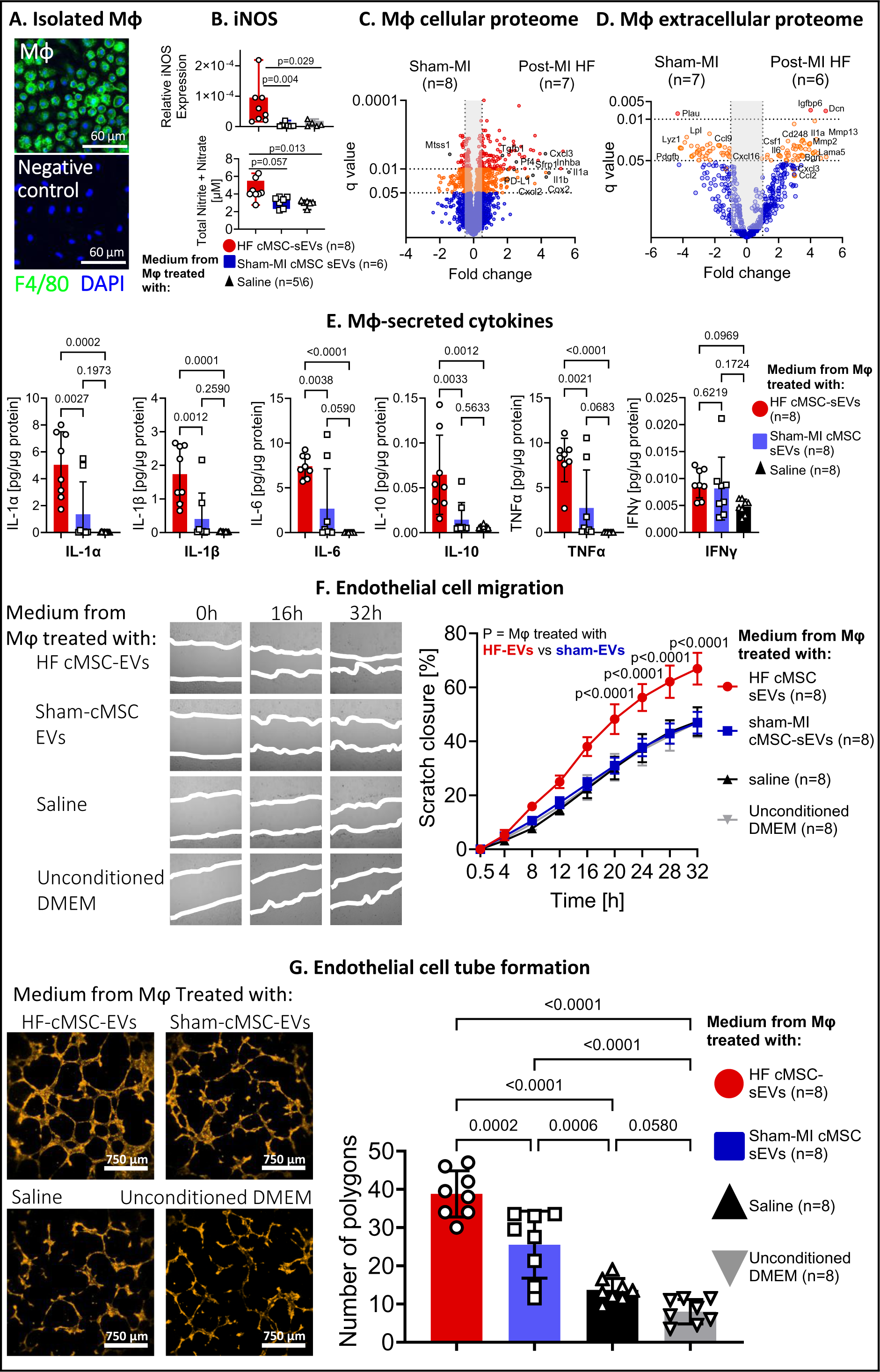
cMSCs from Post-MI Failing Hearts Transform Macrophages into a Tumor-Promoting Phenotype. A) To study the effect of cMSC-sEVs on macrophage activation, we isolated peritoneal macrophages from female C57BL/6 mice using a resistance to trypsinization assay. Staining for F4/80 and DAPI confirmed the purity of isolated macrophages. The purity of isolated macrophages reached 96%. Scale bars: 20 nm for all panels except the right panel, 40 nm. B) To assess the activity of inducible nitric oxide synthase (iNOS) in cMSC-sEVs treated macrophages, we incubated macrophages with cMSC-sEVs (10^9^ EVs/mL) from the failing heart, sham-operated heart, or saline for 24 hours in serum-free conditions. Then, cells were washed and incubated for another 24 hours in a serum-free medium. After 24 hours, we collected the macrophage-conditioned medium and isolated total macrophage RNA. We used RT-PCR to determine the expression iNOS in isolated macrophages and measured the total concentration of nitrite and nitrate in the conditioned medium of incubated macrophages. Expression of both iNOS was upregulated after incubation with cMSC-sEVs from failing hearts, as well as the concentration of nitrites and nitrates in conditioned medium. P values were determined by Kruskal Wallis test with Dunn’s post-test. P for iNOS = 0.0003, P for nitrites and nitrates =0.004. P values for specific comparisons are indicated on the graph. C-D) Next, we incubated macrophages with cMSC-sEVs (10^9^ EVs/mL) from the failing heart, sham-operated heart, or saline for 24 hours in serum-free conditions. Then, cells were washed and incubated for another 24 hours in a serum-free medium. After 24 hours, we collected the macrophage-conditioned medium and lysed the macrophages. The proteomes of macrophages lysate (C) and conditioned medium (D) were analyzed to assess cellular and extracellular proteome. P value was determined using multiple t-tests with FDR correction for multiple comparisons. E) To determine if cMSC-sEVs from post-MI HF induces cytokines secretion, we used ELISA assays. P values were determined by one-way ANOVA with Holm-Šídák’s post-test to account for multiple comparisons. Data passed the D’Agostino-Pearson normality test. F-G) cMSC-sEVs from HF mice (10^9^ EVs/mL) induced macrophage into a pro-angiogenic phenotype. First, the conditioned medium from macrophages treated with post-MI HF cMSC- sEVs stimulated migration of the C166 EC line, compared to cMSC-sEVs from sham-MI, demonstrated by a scratch closure assay (E). P values were calculated using two-way repeated measures ANOVA with Holm-Šídák’s post-test. P for HF cMSC-sEVs < 0.0001, p for time < 0.0001, p for interaction < 0.0001. P for HF cMSC-sEVs vs. sham-MI cMSC-sEVs is indicated on the graph. Next, cMSC-sEVs from HF mice (10^9^ EVs/mL) but not sham-MI stimulated macrophages to induce tube formation in angiogenesis assay with C166 ECs (F). Scale bar = 750 μm. P values were determined by one-way ANOVA with Holm-Šídák’s post-test (p< 0.0001). Data passed the D’Agostino-Pearson normality test. P values for specific comparisons are indicated on the graph. Abbreviations: ANOVA - analysis of variance. cMSC-EV - cardiac mesenchymal stromal cell extracellular vesicles.

To further characterize the changes in macrophages after cMSC-sEVs exposure, we analyzed cellular and extracellular macrophage proteins (Fig-3C-D). Comparative proteomics of intracellular macrophage proteins revealed that cMSC-sEVs from the failing heart enriched pathways related to inflammation, immune modulation, angiogenesis, and remodeling of the extracellular matrix (ECM) (Fig-3C, S9A-H). Notably, activated macrophages upregulated the expression of the immune-checkpoint molecule programmed death-ligand 1 (PD-L1) at mRNA (Fig-S9I) and proteomic levels (Fig-S9E). PD-L1 induces inactivation and apoptosis of cytotoxic T lymphocytes and thus promotes immune suppression and tumor growth. This is a significant characteristic of TAMs.^36,41^

Proteomics of macrophage-conditioned medium revealed that cMSC-EVs from post-MI HF induced significant changes in the profile of macrophage-secreted proteins (Fig-3D, S10A-B). These changes included up-regulation of pro-inflammatory cytokines and chemokines, pro-angiogenic mediators, and down-regulation in anti-angiogenic proteins (Fig-S10C-E). Biological pathway analysis revealed enrichment in processes related to tumor formation: inflammation, angiogenesis, ECM remodeling, and depletion in processes related to regulation of proliferation and cell death (Fig-S10F-G). Together, macrophage proteome indicated that cMSC-EVs from post-MI heart educated macrophages into a pro-tumorigenic state with many characteristics of TAM.^17^

Cytokines are among the most significant messengers and effectors in executing macrophage functions.^33,34^ Cytokine array revealed that macrophages exposed to post-MI cMSC-sEVs secreted higher amounts of cytokines implicated in pathogenesis of cardiovascular diseases and cancer, namely IL-1α and β, IL-6, IL-10, and TNF-α (Fig-3E).^35^ Thus, cMSC-sEVs from post-MI HF educated macrophages to express a unique cytokine profile with pro-tumorigenic properties.

Pro-angiogenic macrophages contribute to tumor angiogenesis, growth, and metastasis.^36^ We found that conditioned medium from educated macrophages stimulated endothelial cell migration (Fig-3F) and enhanced angiogenesis by endothelial tube formation assay (Fig-3G).

Finally, we studied whether macrophages educated by cMSC-EVs from post-MI hearts modulate proliferation or migration of lung (LCC) and colon (MC38) cancer cells. We found that conditioned medium from educated macrophages did not affect the proliferation and migration of LLC cells in vitro (Fig-S11A-B) but slightly reduced MC38 cell proliferation and did not affect migration (Fig-S11C-D). Collectively, cMSC-sEVs from the failing heart drive macrophages toward pro-angiogenic, pro-inflammatory, immune-modulatory, and pro-tumorigenic phenotypes. However, educated macrophages did not affect cancer cell proliferation and migration.

#### Biodistribution and Uptake of cMSC-sEVs

Tracking in vivo destination of cEVs is essential for understanding their contribution to tumor growth. To assess the distribution of cMSC-sEVs in vivo, we used a mouse model of adoptive transfer. We first labeled cMSC-sEVs from MI hearts with a near-IR fluorescent dye. Next, we injected the labeled cMSC-sEVs systematically into the left ventricle (LV) cavity of mice 10 days after MI or sham-MI, bearing heterotopic LLC tumors (Fig-4A-C) and assessed the distribution of cMSC-sEVs by bioluminescent IVIS Spectrum imaging. We found increased fluorescent activity in situ in the chest of mice with infarcted hearts, compared with sham-MI hearts (Fig-4D- E). Next, to improve the signal and reduce interferences, we ex vivo scanned the internal organs, including tumors 24 hours after injection (Fig-4F-G). We found enhanced fluorescent activity of sEVs in the liver, kidneys, lungs, spleen, bones, and tumors (Fig-4F-G).

**Figure 4:**
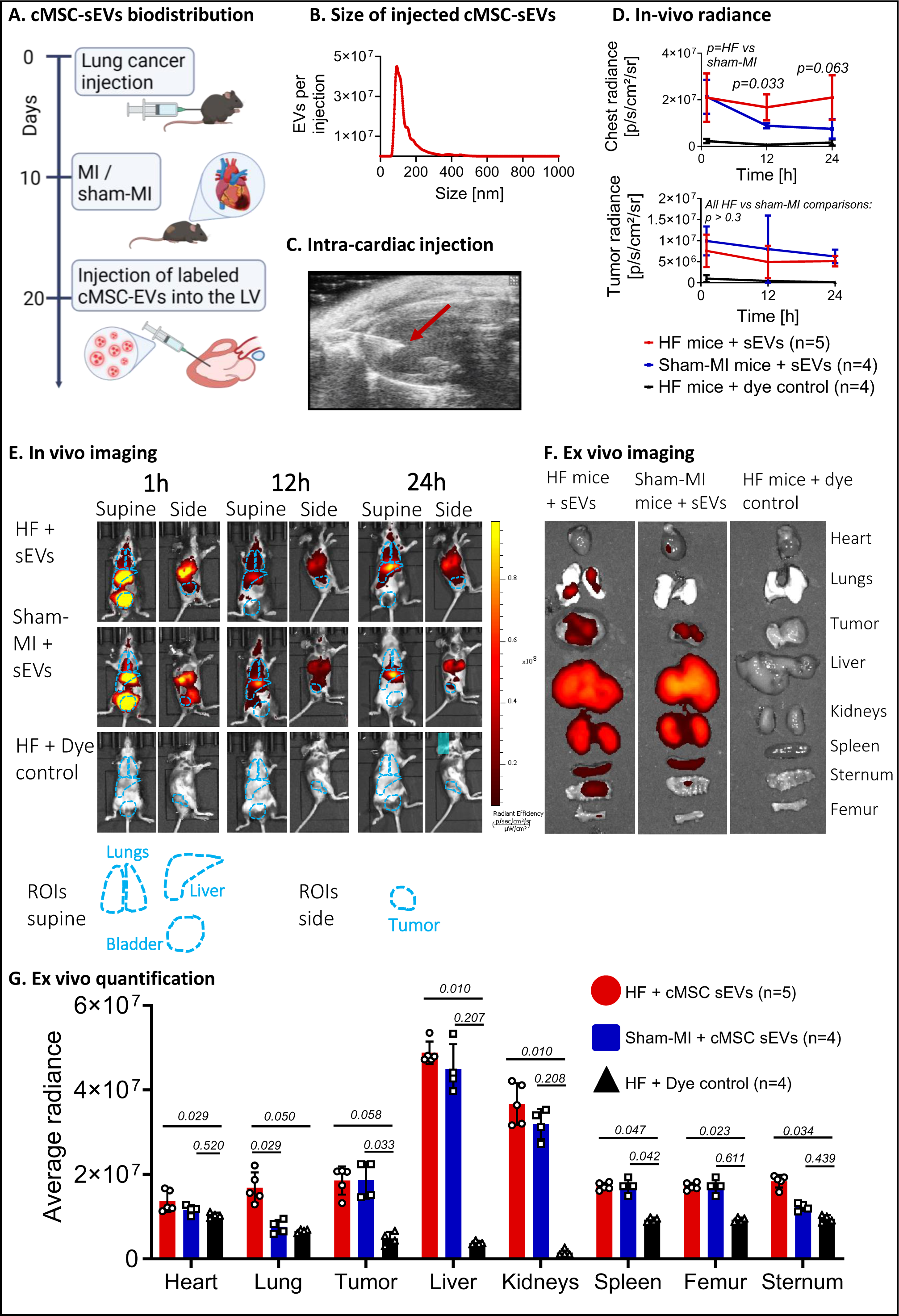
Biodistribution and Uptake of cMSC-sEVs. A) To monitor the biodistribution of cMSC-sEVs, we used IVIS Lumina LT for both in vivo and ex vivo imaging. We isolated cMSC-sEVs 10 days after MI from donor mice, labeled them with near-infra-red dye, and injected them to the left ventricular cavity of tumor-bearing recipient mice after either post-MI HF or sham-MI. We used the same amount of dye for control in a sample without sEVs. B) Size distribution of injected cMSC-sEVs from HF mice was in the small EV range. C) Representative image for echocardiography-guided LV cavity injection of labeled EVs. The red arrow indicates the needle. We confirmed the success of the injection by echocardiography imaging of a bubble jet in the LV cavity during injection. D) Quantification of the fluorescence activity of cMSC-sEVs in vivo. We quantified the radiance of the chest and the tumor area 1, 12 and 24 hours after injection. We found that post-MI HF enhanced the retention of cMSC-sEVs in the area of the lungs. In addition, we found fluorescence activity in the tumor area of both post-MI HF and sham-MI mice. P values were calculated using two-way repeated measures ANOVA with Holm-Šídák’s post-test. For the retention in the lungs: P for post-MI HF = 0.0001, p for time = 0.0785, p for interaction = 0.1357. For the retention in the tumors: P for post-MI HF = 0.0003, p for time = 0.2685, p for interaction= 0.9263. P for post-MI HF vs. sham-MI is indicated on the graph. E) Representative images of fluorescence activity in mice with and without HF after injection of labeled cMSC-sEVs from post-MI HF mice. Post-MI HF facilitated the distribution and retention of cMSC-sEVs in the lungs and we detected fluorescence activity in the tumor area of both post-MI HF and sham-MI mice. Outlines of the lungs, liver, bladder, and tumor are depicted in dotted light blue lines. F) To reduce background fluorescence, enhance fluorescence signal, and evaluate the distribution of cMSC-sEVs from HF mice to specific organs, we performed ex vivo analysis 24 hours after cMSC-sEVs injection. The highest signal was detected in the liver and kidneys, and an intermediate signal in the tumor and spleen. Furthermore, we confirmed our in situ findings and showed that HF promoted cMSC-sEVs distribution to the lungs. G) Quantification of the fluorescence activity of cMSC-sEVs ex vivo. We quantified the radiance of the heart, lungs, tumor, liver, kidneys, spleen, sternum, and femur 24 hours post-injection. P values were calculated using Kruskal-Wallis with Dunn’s post-test. P for heart=0.024, lungs=0.003, tumor=0.008, liver=0.002, kidneys=0.002, spleen=0.009, sternum=0.017 and femur<0.0001. P values for specific comparisons are indicated on the graph. Illustration created with BioRender.com.

Interestingly, after MI, the accumulation of post-MI HF cMSC-sEVs in the congested lungs was greater than in the lungs of sham-MI mice (Fig-4F-G). Together, our biodistribution studies indicated that cMSC-sEVs biodistributed to various organs and tumors. MI promoted cMSC- sEVs accumulation in congested lungs.

#### cMSC-sEVs from the Failing Heart Accelerated Tumor Growth

Lung cancer is a frequent and leading cause of death in patients with HF.^6-8,42^ To investigate the role of cMSC-sEVs in tumor growth in vivo, we first inoculated LLC cells in the dorsal aspect of the limb of female C57BL/6 mice to create a heterotopic lung cancer model. Then,10 days later, we randomized mice bearing LLC tumors to MI or sham-MI (Fig-S2B). The mortality rate was 11% (1 out of 9) or null after MI or sham-MI. Consistent with previous reports,^10,11^ we found that MI and LV dysfunction were associated with accelerated tumor growth (Fig-S2D-E).

To dissect the effect of cMSC-sEVs from other factors released from the post-MI failing heart, we used a model of adoptive transfer of cMSC-sEVs to LLC tumor-bearing mice with normal hearts (Fig-5A). We subcutaneously injected cMSC-sEVs (2μg of EV protein, Fig-S12A) or saline into the cancer cell inoculation site. We found that transfer of cMSC-sEVs from failing hearts promoted the appearance of visible tumors (Fig-S12B). Moreover, cMSC-sEVs from the failing heart accelerated tumor growth compared with sham-MI EVs or saline (Fig-5B), as also confirmed by heavier tumors in mice treated with cMSC-sEVs from failing hearts (Fig-S12E). Finally, histological analysis for expression of Ki67, a marker of cell cycle activity and tumorigenesis, revealed a higher percentage of Ki67 in tumors treated with cMSC-sEVs from the failing heart at 16 days (before differences in tumor volumes accelerated) and 28 days after tumor inoculation (Fig-5C,D, S12C). Mechanistically, we found that the percentage of Ki67 correlated with tumor volume at day 28, but not at day 16 (Fig-5C, S12F-G). Thus, changes in tumor cell proliferation preceded the acceleration in tumor growth. Overall, cMSC-sEVs from the failing heart stimulated tumor cell proliferation and growth.

**Figure 5:**
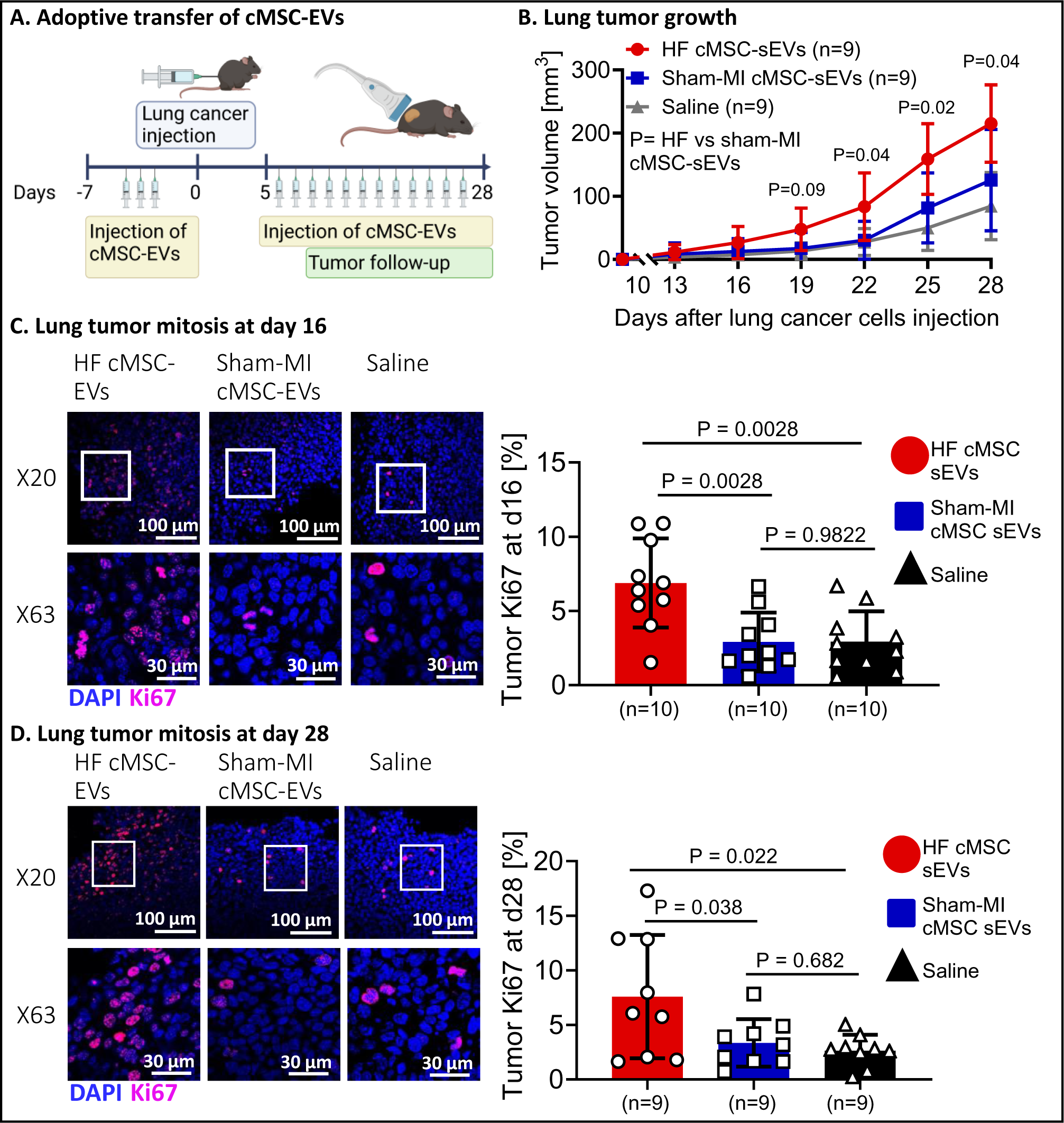
cMSC-sEVs from Post-MI Failing Hearts Accelerated Tumor Growth. A) To investigate the effects of cMSC-sEVs from other soluble factors, we randomized mice to receive an equal amount of cMSC-sEVs from post-MI HF, sham-MI (2 μg of EVs’ protein), or saline every 48 hours. Each mouse received 3 subcutaneous injections to the inoculation site during the week before LLC inoculation (750,000 cells in 100 µL saline) and another 12 injections resuming 5 days after inoculation. A) We monitored tumor growth with serial ultrasound examinations. Mice receiving SC injections of cMSC-sEVs from failing hearts developed larger tumors than mice treated with sham-MI cMSC-sEVs. P values were determined by two-way repeated measures ANOVA with Holm-Šídák’s post-test. P for cMSC-sEVs from HF = 0.0023, p for time < 0.0001, p for interaction < 0.0001. P for HF cMSC-sEVs vs. sham-MI cMSC-sEVs indicated on the graph. C-D) To determine if cMSC-sEVs from the failing hearts facilitated cancer cell proliferation, we stained tumor sections for Ki67 and assessed the number of mitoses. We found a higher percentage of mitoses from tumors of mice who received cMSC-sEVs from failing hearts, compared with sham-MI cMSC-sEVs or saline. This higher rate of tumor cell mitosis was detected on day 16 (D), before any differences in tumor volume, and on day 28 (C). P values were determined by one-way ANOVA with Holm-Šídák’s post-test. P for day 28 = 0.0154, p for day 16 = 0.0009. P values for specific comparisons are indicated on the graph. Normality was tested by the D’Agostino-Pearson omnibus test. Scale bars: 100 μm (upper panels) and 30 μm (lower panels). Abbreviations: cMSC - cardiac mesenchymal stromal cells. EVs - extracellular vesicles. LLC- Lewis lung cancer. LV - left ventricle. NTA - nanoparticle tracking analysis. SC - subcutaneous. Illustration created with BioRender.com.

#### EV Depletion Reduced the Tumor-Promoting Effects of Post-MI HF

Interfering with EV secretion and uptake reduces tumor growth, impedes metastatic progression, and inhibits the systemic effects of cancer.^16^ To determine whether EV depletion would reduce the effects of post-MI HF on tumor growth, we used GW4869, an inhibitor of the enzyme that converts membrane sphingomyelin into ceramide, which is required for formation of EVs.^43^ LLC cells were inoculated to the hindlimb of mice, and 10 days later, mice were randomized for MI or sham-MI operation. Starting 3 days after MI or sham-MI, mice from each group were further randomized to receive intraperitoneal (IP) injections of GW4869 or DMSO (vehicle) every 48 hours (Fig-6A). We confirmed effective EV depletion in isolated sEVs from the whole heart tissue at the end of the experiment (20 days after MI) (Fig-6B). Significantly, EV depletion attenuated tumor growth after MI (Fig-6C) and to lesser extent after sham-MI (Fig-6C, S13A), suggesting that GW4869 also affects pro-tumorigenic sEVs from sources other than heart. Still, the marked reduction in tumor growth in post-MI mice during EV depletion suggested that inhibiting cardiac sEVs after MI contributed to reduced tumor growth.

**Figure 6:**
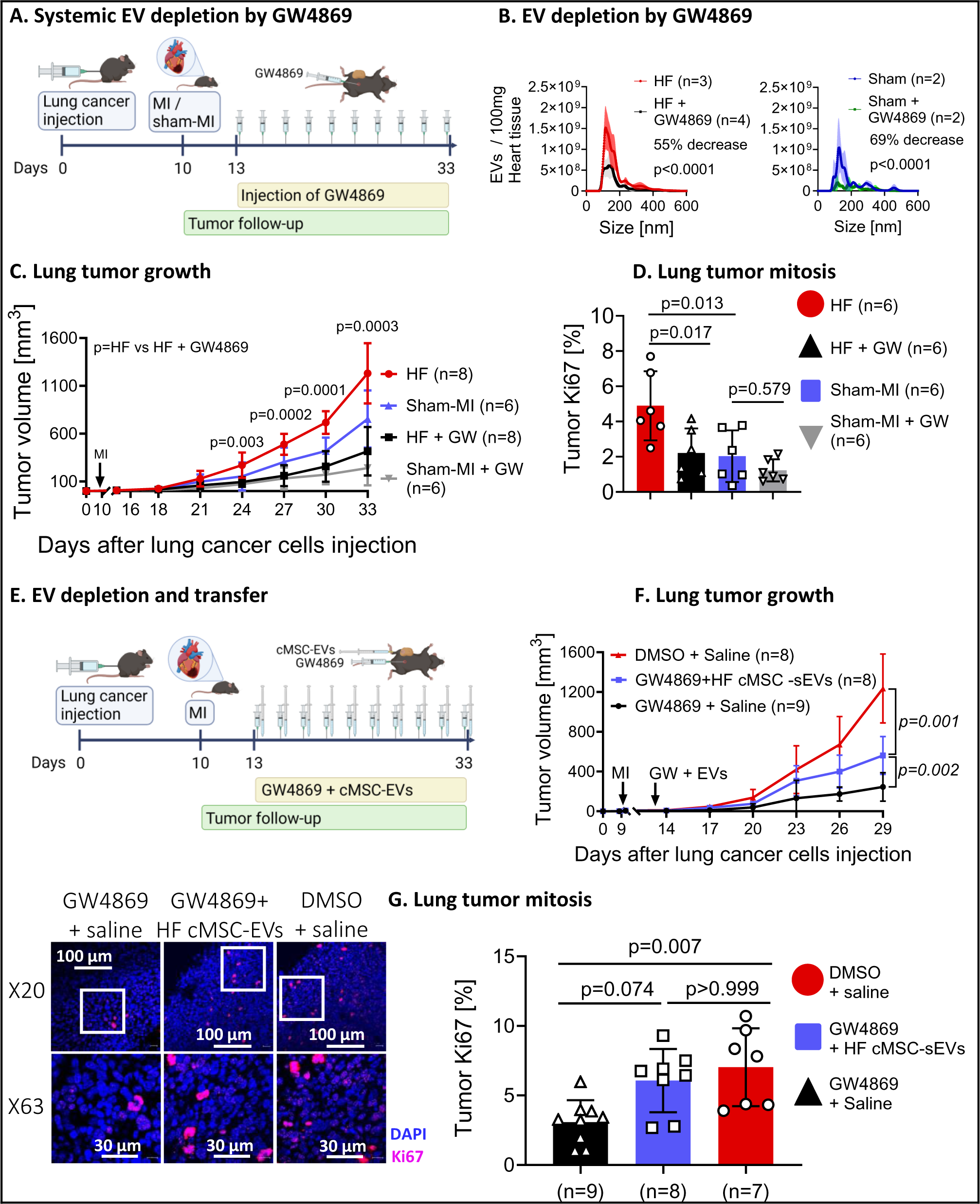
sEV Depletion Attenuated Tumor Growth and Transfer of post-MI cMSC-sEVs and Restored Accelerated Tumor Growth. A) LLC cancer cells (750,000 cells in 100 µL saline) were inoculated into the hindlimb of mice. Ten days later, mice were randomized for MI or sham-MI operation. Starting three days after MI or sham-MI, mice from each group were further randomized to receive intraperitoneal (IP) injections of GW4869 (2.5 mg/kg) or DMSO (GW4869 vehicle as control) every 48 hours. Tumor growth and heart function were assessed by ultrasound and echocardiography. B) To validate that IP injections of GW4869 depleted cardiac EV production, we separated EVs directly from cardiac tissue and analyzed them with NTA. GW4869 successfully reduced the number of EVs in failing hearts (B) and sham-MI hearts (C). P values and percentage change were determined by zero-inflated negative binomial regression. C) Mice with post-MI HF developed larger tumors than sham-MI mice, and EV depletion attenuated the tumor-trophic effects of HF. P values were determined by two-way repeated measures ANOVA with Holm-Šídák’s post-test. P for GW4869, for time, and for interaction are < 0.0001. P values indicated on the graph for post-MI HF with vs. without GW4869. D) To validate our findings, we stained the tumors for Ki67 and assessed tumor cell mitoses. We found a higher percentage of mitoses in tumors from mice with post-MI HF. EV depletion reduced the neoplastic effect of post MI HF. P value determined with two-way ANOVA with Holm-Šídák’s post-test. P for GW4869 = 0.0079, p for post-MI HF = 0.0038, p for interaction = 0.1266. E) Transfer of cMSC-sEVs during systemic EV depletion. We inoculated LLC cells (750,000 in 100 µL PBS) in female C57BL/6 mice and subjected them to MI 10 days later. Three days after MI, we randomized the tumor-bearing mice to 3 treatment groups: GW4869 (intraperitoneal, 2.5 mg/kg), and then 2 μg of cMSC-sEVs from the failing heart, subcutaneously above the tumor (n=8) every 48 hours; 2) GW4869 (intraperitoneal, 2.5 mg/kg) and saline (n=8) every 48 hours; or 3) DMSO (the vehicle of GW4869 as control) and saline (n=8) every 48 hours. We monitored cardiac function and tumor volume with echocardiography and ultrasound. F) Transfer of cMSC-sEVs from the failing heart accelerated tumor growth despite systemic EV depletion. P values were determined by two-way repeated measures ANOVA with Holm-Šídák’s post-test. P for HF cMSC-sEVs, for time, and for interaction are < 0.0001. P for specific comparisons indicated on the graph. G) We stained tumor sections for Ki67 and assessed tumor cell mitosis. We found thattransfer of cMSC-sEVs from the failing hearts increased the number of mitoses in the tumor despite systemic EV depletion. P value was determined by Kruskal Wallis (p=0.0063) and Dunn’s post-test. P values for specific comparisons are indicated on the graph. Scale bars: 100 μm (upper panels) and 30 μm (lower panels). Abbreviations: DMSO - dimethyl sulfoxide. IP - intraperitoneal. LLC - Lewis lung cancer. LV - left ventricle. Illustration created with BioRender.com

Staining of tumor slides for Ki67 confirmed increased cell cycle activity in tumors from mice with post-MI HF. Significantly, GW4869 reduced the percentage of Ki67 in tumor cells (Fig-6D, S13B). Thus, EV depletion reduced tumor growth, at least in part, by reducing cancer cell proliferation. Moreover, sEV depletion reduced the power of the inverse correlation between LVEF and tumor volume (Fig-S13C). Overall, our data suggested that EV depletion decreases the association between post-MI HF and lung cancer growth.

The inhibitory effect of the GW4869 on the growth of tumors could be linked to a systemic depletion of EVs and other mechanisms independent of the action of cEVs. To dissect the role of cEVs in accelerating tumor growth and differentiate it from EVs from other sources, we created a new model of systemic EV depletion with transfer of cMSC-sEVs in mice with HF (Fig-6E). Again, GW4869 depleted EVs in the hearts (Fig-S13D-E), and while systemic EVs depletion suppressed LLC tumor growth, transfer of cMSC-sEVs from the failing heart re-accelerated tumor growth after MI (Fig-6F, S13F). The percentage of Ki67-positive cells in tumor sections (Fig-6G), confirmed the pro-tumorigenic effects of cMSC-sEVs. We thus showed that cMSC-sEVs contribute to the tumor-promoting effects of post-MI HF independently of EVs from other sources.

#### cMSC-sEVs Promoted Growth of Orthotopic Lung Tumors

Studying cancer in its natural environment can provide unique insights into tumor biology, tumor-host interactions, and the factors influencing tumor progression. Thus, we performed orthotopic LLC cell transplantation, injecting luciferase-expressing LLC cells into the tail vein and tracked the spreading LLC cells into lungs (also called pulmonary metastasis assay).^9^ We assessed tumor spreading and growth by bioluminescence and micro-CT imaging, with and without a nanoparticle-based approach, utilizing glucose-functionalized gold nanoparticles (GF- GNPs) as a metabolically targeted CT contrast agent (glucose coating enhances uptake by cancer cells).^44^

Consistent with the heterotopic model, post-MI HF promoted tumor growth (Fig-S14A-B). We found that cMSC-sEVs from the post-MI HF hearts accelerated colonization and growth of LLC cells in the lungs of mice after MI (Fig-7A-G). Significantly, systemic sEV depletion by GW4869 suppressed the tumor-promoting effects of post-MI HF by a factor of 4.5 (Fig-7C-F). Transfer of cMSC-sEVs from post-MI hearts, but not from sham-MI, partially re-established the tumor-promoting effects of post-MI HF (Fig-7C-G). These findings were supported by CT imaging of lungs with GF-GNPs (Fig-S15 A-B). Together, we confirmed and supported our earlier findings that post-MI HF cMSC-sEVs contributed to tumor growth.

**Figure 7:**
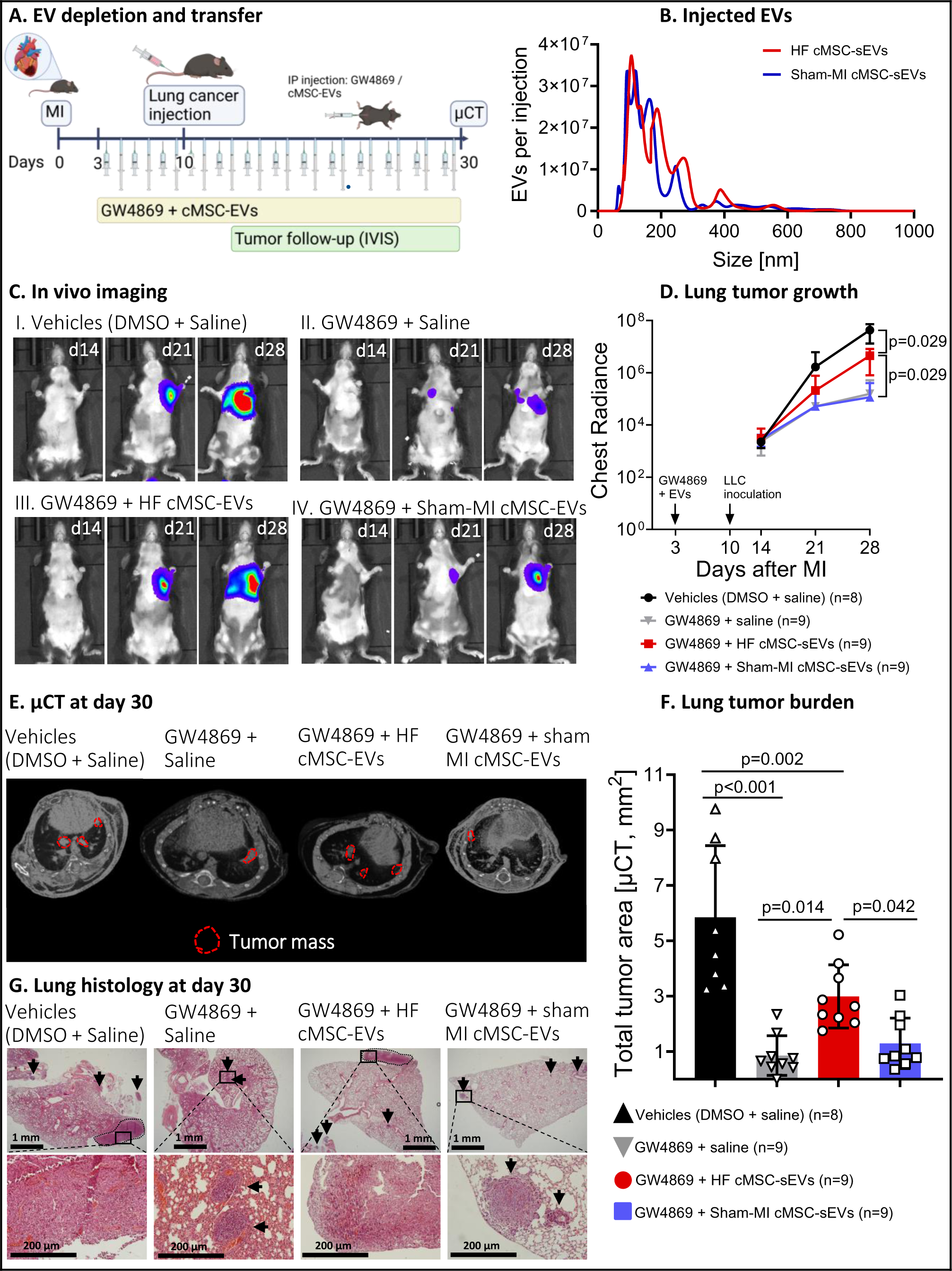
cMSC-sEVs from Post-MI Failing Hearts Promoted Growth of Orthotopic Lung Tumors. A) To investigate the role of cMSC-sEVs in spreading and growth of lung tumors, we used an orthotopic lung cancer model, EV depletion by GW4869, and cMSC-sEVs transfer from either post-MI HF or sham-MI mice. We induced post-MI HF in female C57BL/6 mice, and three days after MI, we randomized the tumor-bearing mice to 4 treatment groups: GW4869 (intraperitoneal, 2.5 mg/kg), and then 10 μg of cMSC-sEVs from the failing heart, IP (n=9) every 48 hours; 2) GW4869 (intraperitoneal, 2.5 mg/kg), and then 10 μg of cMSC-sEVs from sham-MI, IP (n=9) every 48 hours. 3) GW4869 (intraperitoneal, 2.5 mg/kg) and saline (n=9) every 48 hours; or 4) DMSO (the vehicle of GW4869 as control) and saline (n=8) every 48 hours. Treatment with GW4869 started on day 3, and cMSC-sEVs transfer started on day 4 to avoid mixing cMSC-sEVs with the GW4869 solution containing DMSO. Then, we injected luciferase-expressing LLC cells (750,000 in 100 µL PBS) into the tail vein on day 10 after MI. We monitored cardiac function with echocardiography and the development of lung tumors with IVIS on days 14, 21, and 28. In addition, we scanned the lungs with and without injection of GF- GNPs at day 30 using micro-CT. B) Size distribution of injected cMSC-sEVs from post-MI HF or sham-MI mice was mostly in the small EV range. C) Representative images of the bioluminescence of the mice at days 14, 21, and 28 post-MI. D) Quantification of chest bioluminescence after MI and cMSC-sEV transfer. GW4869 reduced the growth of LLC tumors and to a lesser extent after transfer of cMSC-sEVs from the post-MI HF heart. Sham-MI cMSC-sEVs did not affect LLC tumor growth. P values were determined by two-way repeated measures ANOVA with Holm-Šídák’s post-test. P for cMSC-sEVs from HF <0.0001, p for time < 0.0001, p for interaction < 0.0001. P for specific comparisons indicated on the graph. E) Representative images of tumor mass in the lungs by micro-CT at day 30. Individual lung tumors are circled with a dotted red line. F) Blinded analysis of total tumor area (tumor burden). cMSC-sEVs from post-MI HF but not from sham-MI mice promoted the growth of tumor masses in the lungs despite systemic EV depletion. Micro CT data was analyzed by a technician blinded to the allocation of mice into experimental groups. All tumor masses identified in the axial plane were confirmed and discriminated from blood vessels using sagittal and coronal planes. P values by one-way ANOVA with Holm-Šídák’s post-test. P < 0.0001. P values for specific comparisons are indicated on the graph. G) Representative images of lung tumor masses by histological staining for hematoxylin and eosin at day 30. The arrows indicate small individual lung tumors. The dotted black line circles the larger tumors, and the box outlines the higher magnification insert below. Scale bars for upper panel (X2 magnification) = 1 mm. Scale bars for lower panel (X20 magnification) = 200 μm. Illustration created with BioRender.com.

We also established an orthotopic model of more malignant, triple-negative breast cancer tumors using the 4T1 cell line implanted in the mammary pad of syngeneic female Balb/c mice (Fig-S16A). We found that post-MI HF accelerated tumor growth modestly (Fig-S16B).

Furthermore, EV depletion by GW4869 had minimal effect on tumor growth (Fig-S16B). Thus, the effect of post-MI HF and the contribution of cardiac sEVs were limited to certain types of cancers.

#### Post-MI Spironolactone Restrained Tumor Growth

Post-MI HF is characterized by maladaptive neurohormonal activation, particularly the renin-angiotensin-aldosterone system (RAAS). To determine if post-MI HF treatment by RAAS inhibitor would affect cMSC-sEVs and tumor growth, we used spironolactone, an aldosterone receptor antagonist (Fig-8A).^45,46^ We found that post-MI spironolactone attenuated post-MI LV remodeling and improved global longitudinal strain (Fig-S17A-C, Fig-S17G-K). Significantly, while post-MI HF accelerated growth of LLC tumors, spironolactone attenuated this effect (Fig-8B, S17F). Moreover, we found that reduced LV remodeling, as indicated by LV dimension, was correlated with smaller tumor volumes in mice with post-MI HF (Fig-S17D). Notably, spironolactone treatment did not reduce tumor volume or weight in sham-MI mice (Fig-8B, Fig S17F), suggesting that spironolactone lacked direct anti-tumor properties. Finally, we found that spironolactone reduced the rate of cell cycle activity (Ki67) in tumor cells in mice with post-MI HF but not in sham-MI mice (Fig-8C, S17E). Collectively, our findings suggest that post-MI spironolactone reduced post-MI tumor growth.

**Figure 8:**
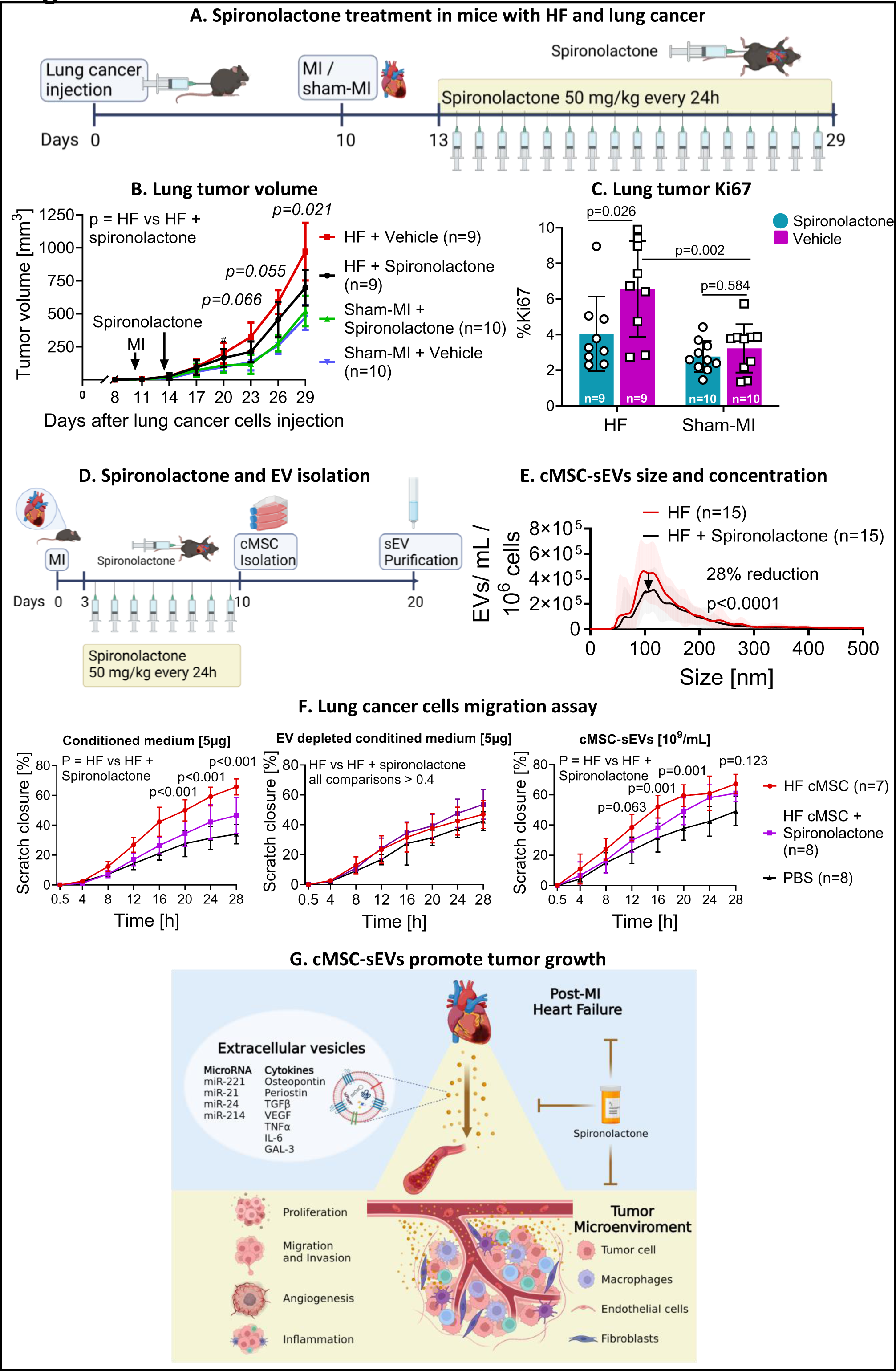
Post-MI Spironolactone Reduced the Tumor-Promoting Effects of MI and HF. A) We inoculated cancer cells (750,000 in 100 µL PBS) into female C57BL/6 mice with LLC cancer cells (750,000 in 100 µL PBS) and randomized them to either MI or sham-MI 10 days later. Three days after MI, we started spironolactone treatment (50 mg/kg, daily) or its vehicle, injected subcutaneously. Finally, we performed serial echocardiographic and ultrasound measurements to assess cardiac remodeling, LV function, and tumor growth. B) While post-MI HF accelerated the growth of heterotopic lung cancer, spironolactone significantly reduced the effect of MI and HF on tumor growth but had no effects on the tumors of sham-MI-operated mice. P values were determined by two-way repeated measures ANOVA with Holm-Šídák’s post-test. P for spironolactone < 0.0001, p for time < 0.0001, p for interaction < 0.0001. P for HF vs HF + spironolactone is indicated on the graph. C) To confirm our findings, we stained the tumors for Ki67 and assessed tumor cell proliferation. We found that spironolactone reduced the number of proliferating cells in tumors from mice with HF but had no effect on the tumors of sham-operated mice. P value determined by two-way ANOVA with Holm-Šídák’s post-test. P for spironolactone = 0.018, p for post-MI HF = 0.0005, p for interaction = 0.0935. P for specific comparisons indicated on the graph. D) To determine if spironolactone will reduce the number or function of cMSC-sEVs secreted by the failing heart, we subjected female C57BL/6 mice to MI and started spironolactone treatment by SC injection (50 mg/kg, daily) three days later. On day 10, we harvested the hearts and isolated cMSCs. E) We purified sEVs from the cMSC-conditioned medium and counted them by NTA. We found that spironolactone reduced EV release by cMSC by 28%. P values and percentage change were determined by zero-inflated negative binomial regression. F) To test the effects of post-MI spironolactone on the ability of cMSC-sEVs to induce lung cancer cell migration, we used a scratch assay. To examine the effects of cMSC-sEVs from soluble factors, we tested the conditioned medium (5 μg of proteins), purified cMSC-sEVs (10^9^/mL), and EV-depleted medium (5 μg of proteins). P value determined by two-way ANOVA with Holm-Šídák’s post-test. For conditioned medium: P for spironolactone < 0.0001, p for time < 0.0001, p for interaction < 0.0001. For purified cMSC-sEVs: P for spironolactone < 0.0001, p for time < 0.0001, p for interaction = 0.0430. For EV depletion medium: P for spironolactone = 0.1145, p for time < 0.0001, p for interaction = 0.0167. P for HF vs HF + spironolactone is indicated on the graph. G) A schematic presentation describes how cardiac cMSC-sEVs transmit signals from the failing heart and accelerate tumor growth. This mechanism could be a target for risk stratification and treatment. Spironolactone treatment reduces the release of cMSC-sEVs and tumor growth. Abbreviations: TGFβ - transforming growth factor beta. TNFα - tumor necrosis factor-alpha. IL-6- interleukin 6. VEGF - vascular endothelial growth factor. Illustration created with BioRender.com.

To further dissect the specific function of cMSC-sEVs during post-MI spironolactone treatment, we assessed the effects of spironolactone treatment on the generation of cMSC- sEVs (Fig-8D). We found that post-MI spironolactone reduced the number of cMSC-sEVs by 28% (Fig-8E). Then, we used a conditioned medium of cMSCs from post-MI HF hearts and found post-MI spironolactone attenuated the effects on LCC cell migration (Fig-8F, left), but not proliferation (Fig. S18A). Next, we isolated cMSC-EVs from the conditioned medium and tested equal amounts of cMSC-sEVs and EV-depleted medium (Fig-S18B). We found that the sEV- depleted medium did not accelerate lung cancer cell migration, and the inhibitory effects of post-MI spironolactone vanished (Fig-8F, middle). In contrast, post-MI spironolactone attenuated the effects of pure cMSC-sEVs on accelerated LLC cell migration and proliferation (Fig-8F, right, Fig S18C). Together, the effect of post-MI spironolactone to reduce accelerated LCC tumor growth was mediated by modulating cMSC-sEVs.

Notably, post-MI spironolactone treatment did not reduce the effects of cMSC-sEVs on proliferation and migration of MC38 colon cancer cells (Fig-S18D-G). Additionally, post-MI spironolactone did not modify the effects of cMSC-sEVs on C166 EC migration (Fig-S18H-I) but mildly reduced EC permeability (Fig-S18J). Together, our data suggested that the anti-tumor effects of post-MI spironolactone were primarily mediated by reducing the number of pro-neoplastic cMSC-sEVs.

## DISCUSSION

Our work provides several new findings. First, we present evidence that post-MI failing hearts, specifically cMSCs, secrete sEVs with pro-tumorigenic properties. cMSC-sEVs transfer multiple factors, including proteins, cytokines, and miRs that stimulate neoplastic growth and activate pro-tumorigenic macrophages (Fig-8G). Second, the pro-tumorigenic effects of cMSC-sEVs are tumor-dependent: highest with lung cancer and modest or absent with other types of cancer. Third, EV depletion reduces the pro-tumorigenic effects of post-MI HF. Finally, post-MI spironolactone treatment decreases the number of cMSC-sEVs and suppress tumor growth. Together, we add new insight into the evolving field of reverse cardio-oncology, with potential clinical implications.

### Comparison with previous reports

Our work builds upon prior studies.^9-11,47^ The inflammatory and reparative responses to myocardial injury and failure may promote neoplastic growth by secreting inflammatory, reparative, immunomodulatory, and tumor-promoting mediators. For example, aortic constriction promotes growth of breast and lung tumors in mice.^9^ Periostin has been identified as a potentially important mediator, and eliminating plasma periostin abolished the effects on tumor cell proliferation in vitro.^9^ However, inhibition of soluble factors, including periostin, has never been studied in vivo. We found that periostin levels were higher in both cMSC-sEVs and cMSC-conditioned medium from post-MI failing hearts. Furthermore, our work identifies other tumor-promoting factors, including cytokines and miRs within cMSC-sEVs. Whether sEVs carry sufficient quantities of cytokines and miRs to prompt changes in cancer cells has been questioned.^16^ Here, specific mediators, such as TGF-β, Gal-3, and miRs, were enriched in sEVs. Such mediators can influence tumor progression by promoting pre-metastatic niche formation and metastasis.^16^ Overall, we suggest that rather than a single factor, multiple factors encapsulated by cMSC-sEVs contribute to the link between heart disease and cancer.

A recent report has suggested another mechanism linking sEVs to accelerated tumor growth. In post-MI HF, plasma sEVs suppress the sensitivity of cancer cells to ferroptosis, an iron-dependent form of apoptosis, thereby promoting tumor growth.^48^ The authors proposed that miR-22-3p-enriched EVs from cardiomyocytes are pivotal in this mechanism.^48^

Notably, a previous report showed that MI and HF did not accelerate growth of renal cancer tumors.^47^ This finding dovetails with our findings and clinical observations, suggesting that the effects of heart disease on tumor growth cannot be generalized to all types of cancer.

Macrophages play a significant role in tumor initiation, progression as well as cytotoxic response.^36,39^ Koelwyn et al. have shown that MI induced a sustained increase in circulating monocytes that targeted and accelerated tumor growth.^11^ Significantly, monocyte depletion inhibited tumor growth in mice subjected to MI.^11^ Here, we show that cMSC-sEVs from post-MI heart educate macrophages toward pro-tumorigenic mode. The educated macrophages share specific characteristics, such as pro-inflammatory, angiogenic, ECM remodeling, and immunosuppressive properties with TAM.^36^ Interestingly, educated macrophages upregulated the expression of the checkpoint molecule programmed death-ligand 1(PD-L1), which interferes with the ability of the immune system to eliminate cancer cells and subsequently accelerates tumor growth.

In terms of translation, we show that post-MI spironolactone reduces adverse LV remodeling, number of cMSC-EVs, and tumor growth. Our findings are significant because they may help to improve the outcome of patients with HF and cancer. A recent retrospective analysis has shown that RAAS inhibitors reduced mortality and tumor recurrence in hypertensive patients after cancer surgery.^49^ Thus, effective HF therapy may reduce the tumorigenic effects of post-MI HF.

#### Limitations and Strengths

First, we used GW4869 to inhibit the production of cEVs. However, GW4869 induces systemic EV depletion and perhaps some other off-target effects. Moreover, GW4869 may also exert its effects via mechanisms other than EV depletion. Today, however, there are no specific, safe inhibitors of EV production, release, or uptake. Still, by cMSC-EV transfer, we demonstrated the independent tumor-promoting effects of cMSC-sEVs in the presence of systemic EV depletion. Second, the methods of infusing sEVs in experimental animals or adding them to cell cultures have been questioned.^18^ The administration of exogenous EVs has the limitation of not fully replicating the natural release of cardiac sEVs. However, because the failing heart secretes additional factors, such as soluble proteins,^9-11^ our approach allows us to study and dissect the role of cMSC-sEVs independent of other secreted factors. Third, although our results suggest a significant contribution of cMSCs-EVs to tumor growth, we acknowledge the need to investigate the role of sEVs from other cardiac cells in tumor growth.

#### Summary, Implications, and Future Research

For the first time, we show that cMSCs from post-MI failing heart secret sEVs that are enriched with multiple pro-tumorigenic factors. The sEVs were uptaken by cancer cells and accelerated tumor growth. Our findings that post-MI spironolactone attenuated the association between post-MI HF and tumor growth may have important translational implications. In addition, our results may be relevant to the association between cancer and other cardiovascular diseases, such as atherosclerosis and aortic stenosis. By better understanding the biology and physiology of cardiac sEVs, we may be able to identify new biomarkers and develop more effective treatments for cancer associated with cardiovascular diseases.

## Abbreviations

cEV: cardiac extracellular vesicles.
cMSC: cardiac mesenchymal stromal cells.
cMSC-sEVs: cardiac mesenchymal stromal cells small extracellular vesicles.
COL1α: Collagen 1 alpha.
DMSO: dimethyl sulfoxide.
ECM: extracellular matrix
ELISA: enzyme-linked immunosorbent assay.
EV: extracellular vesicles.
FDR: false discovery rate.
GF: GNPs-glucose-functionalized gold nanoparticles
HF: heart failure.
IL: interleukin.
IRR: incidence rate ratio.
LLC: Lewis lung carcinoma.
LV: left ventricle.
LVIDs: left ventricular diameter during systole. MI- myocardial infarction
NT-proBNP: N-terminal pro-brain natriuretic peptide
NTA: nanoparticle tracking analysis
miR: micro-RNA.
NTA: nanoparticle tracking analysis.
SEC: size exclusion chromatography.
sEVs: small extracellular vesicles.
TEM: transmission electron microscopy.
TGFβ: Transforming growth factor beta.
TNFα: tumor necrosis factor-alpha,
TSG101: tumor susceptibility gene 101.
VEGF: vascular endothelial growth factor.

## Acknowledgments

We thank N. Ziv-Crispel and Nir Lewis for skillful English-language editing. We thank Dr. Doron Aronson for helpful statistical advice. This work was performed in partial fulfillment of requirements for the PhD degree of Tal Caller, Faculty of Medicine, Tel Aviv University, Israel.

