## supplementary materials for "Small Extracellular Vesicles from Failing Heart Accelerate Tumor Growth"

**Data Supplement**

**Detailed Methods**

The data that support our findings are available from the corresponding author upon reasonable request.

All animal experiments were conducted in accordance with the National Institutes of Health (NIH) Guide for the Care and Use of Laboratory Animals and complied with the standards and approval of the Sheba Medical Center Institutional Animal Care and Use Committee.

**Model of MI and Sham-MI**

Our model of MI in mice has been previously described.[1,2] In brief, female C57BL/6 mice (ENVIGO, Jerusalem, Israel) were anesthetized with inhalation of 2% isoflurane, the chest was opened by gentle dissection, and the pericardium was removed to provide access to the left ventricular free wall. A 9-0 prolene suture was used to ligate the left anterior descending (LAD) coronary artery at the lower border of the left atrium. MI was confirmed by visual blanching distal to the occlusion site and by echocardiography 48 hours after ligation of the LAD. For sham-MI, the suture was passed around the LAD and removed without ligation. The thoracic incisions were closed with 5-0 silk sutures.

**Mice Models of Heart Failure and Cancer**

<u>Heterotopic lung cancer model:</u> Lewis lung cancer (LLC) cells were inoculated heterotopically to the subcutis of the right hindlimb of female C57BL/6 mice (750,000 cells in 100 µL PBS), 10 days before MI or sham-MI. We used small animal ultrasound (Vevo 2100 Imaging System, VisualSonics Inc, Toronto, Ontario, Canada) to measure tumor volume every 3 days.

Orthotopic lung cancer model: Female C57BL/6 mice were randomized for either MI or sham-MI procedures. LLC cancer cells expressing stable luciferase (1,500,000 cells in 100 µL PBS) were injected into the tail vein 10 days after the procedure. We monitored the formation and growth of lung tumors by serial luminescence measurements and micro-CT.

Orthotopic triple-negative breast cancer model: Female Balb/C mice were randomized for either MI or sham-MI. At day 10, 4T1 cancer cells (250,000 cells in 100 µL saline) were inoculated into the mammary pad of the mice. Tumor growth and heart function were assessed by ultrasound and echocardiography.

**Echocardiography and Tumor Ultrasound**

To monitor LV remodeling, function, as well as tumor growth, we used a small animal ultrasound system (Vevo 2100 Imaging System, VisualSonics Inc, Toronto, Ontario, Canada) equipped with a 22- to 55-MHz linear-array transducer (MS550D MicroScan Transducer). The echocardiography protocol was guided by the position paper of the ESC Working Group on Myocardial Function.[3] Light anesthesia was induced by inhalation of 2% isoflurane/98% O2 and subsequently maintained by 1% to 1.5% isoflurane. We controlled the isoflurane flow to maintain the heart rate at >500 bpm. All measurements were averaged for 3 consecutive cardiac cycles. To improve the sensitivity to detect regional wall motion abnormalities in adult mice after MI, we used speckle tracking-based strain analysis to quantify strain in the long axis. Echocardiographic parasternal long-axis images were acquired at a frame rate of 310 frames per second. Three consecutive cardiac cycles were selected, and the endocardial and epicardial borders were traced. If needed, borders were corrected to preserve as precise tracking as possible throughout each cine loop. Analysis performed by Vevo Lab Software version 1.7.1, including Vevo Strain package (VisualSonics Inc, Toronto, Ontario, Canada).

For tumor imaging, we used the same system equipped with a 22- to 55-MHz linear-array transducer (MS250D MicroScan Transducer). Tumor volume was calculated by L·W·D·π/6 (L- tumor longest diameter, W- tumor longest diameter perpendicular to L, D- tumor depth in the plane of L). During ultrasound and echocardiography measurements, mice were anesthetized with inhalation of 2% isoflurane, placed on a warm (37°C) platform, and warm (37°C) ultrasound gel was applied. The heart rate was monitored and kept between 430-500 beats per minute (bpm).

**Isolation and Culture of cMSCs**

cMSCs from the hearts of mice with and without MI were isolated 10 days after MI or sham-MI using enzymatic digestion as previously described.[2] In brief, mice were euthanized, and the heart was extracted and chopped using a scalpel. Cells were extracted with an enzymatic digestion mixture using 3 cycles of incubation at 37°C for 10 minutes on an orbital shaker. Plastic-adhered cells were incubated at 37°C in humid air with 5% $CO_2$ and grown in DMEM (Gibco, Grand Island, New York) with 15% FBS (Gibco, Grand Island, New York), 1% penicillin-streptomycin (Biological Industries, Beit Haemek, Israel), 1% L-glutathione (Sigma-Aldrich, St. Louis, MO, USA), 1% MEM nonessential amino acids (Gibco, Grand Island, New York), and 0.1 mM β-mercaptoethanol (Gibco, Grand Island, New York). The medium was changed 48 hours after plating and subsequently every 3 or 4 days. In each experiment group, we used cultured cMSCs at passage 2. To collect sEVs from cMSC, we only used conditioned medium from fresh (not frozen and thawed) cells. cMSCs were allowed to grow to 85% confluence, and the medium was replaced with serum-free conditions after two subsequent washes. The conditioned medium was collected after 72 hours, and the cells were counted with a hemacytometer.

**Characterization of cMSCs by Flow Cytometry**

To characterize the cells used for cEVs collection and isolation, we used flow cytometry (Cytoflex, BECKMAN COULTER, Pasadena, CA, USA) after staining the cells for fibroblasts markers: COL1α (Collagen 1 Polyclonal Antibody, FITC Conjugated, Bioss, MA, USA) and isotype control (Rabbit IgG Isotype Control, FITC Conjugated, Bioss, MA, USA). CD90.2 (FITC anti-mouse CD90.2 (Thy1.2) Antibody, Biolegend, San Diego, CA, USA) and isotype control (FITC Rat IgG2b, κ Isotype Ctrl Antibody, Biolegend, San Diego, CA, USA). mEF-SK4 (Feeder Cells Antibody, anti-mouse, PE, Miltenyi Biotec, Germany) and isotype control (Isotype Control Antibody, rat IgG1, PE, Miltenyi Biotec, Germany). PDGFRα (APC anti-mouse CD140a Antibody, Biolegend, San Diego, CA, USA) and isotype control (APC Rat IgG2a, κ Isotype Ctrl Antibody, Biolegend, San Diego, CA, USA). Macrophages marker: F4/80 (Alexa Fluor® 647 anti-mouse F4/80 Antibody, Biolegend, San Diego, CA, USA) and isotype control (Alexa Fluor® 647 Rat IgG2a, κ Isotype Ctrl Antibody, Biolegend, San Diego, CA, USA), and the endothelial cells marker CD31 (Alexa Fluor® 647 anti-mouse CD31 Antibody, Biolegend, San Diego, CA, USA) and isotype control (Alexa Fluor® 647 Rat IgG2a, κ Isotype Ctrl Antibody, Biolegend, San Diego, CA, USA). Briefly, cMSCs were lifted with trypsin-EDTA (TrypLE, Gibco, Grand Island, New York), centrifuged, counted, and resuspended in flow cytometry buffer containing 1% bovine serum albumin (Sigma-Aldrich, St. Louis, MO, USA) and 0.1% sodium azid (Sigma-Aldrich, St. Louis, MO, USA) in PBS (Biological Industries, Beit Haemek, Israel). Then, the cells were incubated with polyclonal IgG, washed and incubated with a specific primary antibody or isotype control, washed twice, and resuspended in a flow cytometry buffer containing CellFix (BD Biosciences, CA, USA) for fixation and analysis. For intracellular staining (COL1a), we first used CellFix for fixation, then 0.5% Tween-20 (Sigma-Aldrich, St. Louis, MO, USA) in PBS for permeabilization. Our gating strategy is shown for each experiment and includes the exclusion of debris (FSC-A vs SSC-A), inclusion of single cells (singlets) and not clusters of cells (FSC-A vs. FSC-H), and then gating for positive and negative staining using an appropriate isotype control antibody. The data were analyzed using CytExpert v2.4 (BECKMAN

COULTER, Pasadena, CA, USA). To assess the viability of cMSCs after culture in serum-free condition, we used trypan blue (Gibco, Grand Island, New York) staining.

**Purification of sEVs from the Myocardium**

Our EV separation protocol was guided by the recent position statement of the International Society for Extracellular Vesicles (MISEV2018).[4] The information about EV isolation and separation has been uploaded to the EV-TRACK knowledge base. Readers may access the experimental parameters in the following URL: http://evtrack.org/review.php (EV-TRACK ID: EV230012, Caller, Tal). Small EVs were isolated from myocardial tissue according to modifications from previous reports.[5,6] Briefly, mice were euthanized with isoflurane inhalation, and the hearts were perfused with ice-cold PBS for 5 minutes. The hearts were extracted, chopped, and mixed with scalpel blades for 30 seconds on ice. Then, ice-cold PBS was applied to the tissue, and the mixture was passed through a 100-μm cell strainer (Corning, NY, USA), centrifuged at 1500g for 5 minutes using Heraeus Labofuge 400 centrifuge (Thermo-Fisher Scientific, Waltham, MA, USA) to remove any cells and large particles. The supernatant was collected and then passed through a 28-mm syringe filter, 0.80 μm, SFCA membrane (Corning, NY, USA) and centrifuged at 10000g at 4°C for 10 min, using Sorvall LYNX 4000 Superspeed Centrifuge (Thermo-Fisher Scientific, Waltham, MA, USA). The supernatant was collected and then loaded onto the loading frit of the IZON qEV10-35nm legacy SEC column (IZON, Oxford). The sample was allowed to run into the column according to manufacturer’s instructions, and the eluate was collected in 40 sequential fractions of 2 mL to determine the EV fraction and the protein fraction. The pooled EVs’ fraction was then concentrated by ultrafiltration with AMICON 15 10-kDa tubes (Millipore, Burlington, Massachusetts).

##### Purification of cMSC-sEVs

To separate sEVs from cMSC-conditioned media, we used IZON qEV10-35 size exclusion chromatography columns (IZON, Oxford).[7] Briefly, cMSC-conditioned medium was centrifuged at 1500g for 5 minutes, 4°C using Heraeus Labofuge 400 centrifuge (Thermo-Fisher Scientific, Waltham, MA, USA) to remove any cells and large particles. The supernatant was collected and then passed through a 28-mm syringe filter, 0.80 µm, SFCA membrane (Corning, NY, USA) and centrifuged at 10000g at 4°C for 10 min, using Sorvall LYNX 4000 Superspeed Centrifuge (Thermo-Fisher Scientific, Waltham, MA, USA). The supernatant was collected and then loaded onto the loading frit of the size exclusion chromatography (SEC) column. The sample was allowed to run into the column according to the manufacturer's instructions, and the eluate was collected in 40 sequential fractions of 2 mL to determine the EV fraction and the protein fraction. The pooled EVs fraction was then concentrated by ultrafiltration with AMICON 15 10-kDa tubes (Millipore, Burlington, Massachusetts).

##### Nanoparticle Tracking Analysis (NTA)

The size distribution and concentration of isolated EVs were examined by the Malvern NanoSight NS300 device (Malvern PANalytical, Malvern, UK). The samples were diluted at 1:30 and 1:1 in sterile-filtered PBS and analyzed. The measurements were based on 3 one-minute-long videos, with a screen gain of 10, camera level of 12, and the analysis parameters: detection threshold of 4 and screen gain of 10. Results were analyzed using the software package (version v3.00) nanoparticle tracking analysis.

**Negative Staining with Transmission Electron Microscopy**

cMSC-sEVs of mice with and without MI were loaded onto formvar carbon-coated grids, fixed in 2% paraformaldehyde and washed. The EVs were post-fixed in 2.5% glutaraldehyde, washed, contrasted in 2% uranyl acetate embedded in a mixture of uranyl acetate (0.4%) and methyl cellulose (0.13%), and examined in a JEM-1400Plus Transmission Electron Microscope (JEOL, Tokyo, Japan).

**Cryogenic Electron Microscopy**

For the cryo-EM measurement, 3 μl of SEC-purified sEVs samples were loaded on a glow discharged (EmiTech K100 machine) lacey grid that was blotted and plunged into liquid ethane using a Gatan CP3 automated plunger and stored in liquid nitrogen until use. Frozen specimens (samples with sEVs embedded in vitreous ice) were transferred to Gatan 914 cryo-holder and maintained at temperatures below -176°C inside the microscope. Samples were inspected with a Tecnai G2 microscope (FEI – Thermo-Fisher Scientific, Waltham, MA, USA) with an acceleration voltage of 120 kV, which is equipped with a cryobox decontaminator. Images were taken using Digital Micrograph with a Mulitiscan Camera model 794 (Gatan, CA, USA) in different resolutions.

**Protein Assay**

Isolated EVs were treated with radioimmunoprecipitation assay (RIPA) buffer (Sigma-Aldrich, St. Louis, MO, USA) supplemented with PHOSstop phosphatases inhibitor (Roche, Basel, Switzerland) and cOmplete proteases inhibitor mixture (Roche, Basel, Switzerland). The protein content of isolated EVs was measured by BCA Pierce Protein Assay Kit (Thermo-Fisher Scientific, Waltham, MA, USA) according to the manufacturer's protocol.

#### EV Markers by Western Blot

sEVs were analyzed for surface markers by western blot. Samples were probed for EV markers CD81 and CD9, and exosomal marker TSG101[4] (Santa Cruz Biotechnology, Dallas TX, USA). Appropriate secondary antibodies were used (Santa Cruz Biotechnology, Dallas TX, USA). Briefly, EVs (8 µg of protein) were treated with laemmli X5 sample buffer according to protocol, under reducing conditions to probe for TSG101 and non-reducing conditions to probe for CD81 and CD9. The samples were loaded onto a 12% SGS-PAGE (Bio-RAD, Hercules, CA, USA), run for 15 minutes on 150V, and then the voltage was increased to 200V. We used the wet transfer method (Bio-RAD, Hercules, CA, USA) to blot the proteins onto a 0.22-µm PVDF membrane (Frogga Bio, ON, Canada). The membrane was blocked with 5% BSA and probed with primary antibody and then secondary antibody. Detection was carried out using the enhanced chemiluminescent reagent WESTAR NOVA 2.0 (CYANAGEN, Bologna, Italy). Full membrane images were added to the supplementary material (Figure 7 in the Data Supplement).

#### Proteomic Analysis

For sample preparation, sEVs were lysed in lysis buffer (containing 1.5 M urea, 0.5 M thiourea, 25 mM Tris pH 8.5) and sonicated. For preparations of LLC tumors, lysis buffer containing 4% SDS, 100 mM Tris/HCl pH 7.6 was added to the tumor tissue. Samples were then homogenized, centrifuged at 14,000 rpm, 4°C, 15 minutes, and the supernatant was collected. Supernatant Samples were sonicated, diluted 1:5 in SDS buffer (4% SDS, 100 mM Tris/HCl pH 7.6), and heated to 95°C for 30 minutes. Lysates were precipitated using methanol/chloroform and resuspended in Urea /Thiourea buffer (containing 6 M urea, 2 M thiourea, 100 mM Tris pH 8.5). For EV-depleted secretome of cMSCs, the medium was collected, centrifuged at 1500 g for 5 minutes, 4°C and then centrifuged for 10 minutes at 10000g at 4°C and 70 minutes at 100,000 g (40833rpm) at 4°C. The supernatant was collected. Supernatant samples were concentrated by Amicon Ultra 3K centrifugal filter (Millipore), and their buffer was exchanged to Urea /Thiourea buffer (containing 6M urea, 2M thiourea, 100 mM Tris pH 8.5) on the same Amicon filter by two washes with the buffer. For the proteomic and secretome analysis of macrophages, the conditioned medium was centrifuged at 1500g for 5 minutes, 4°C and the supernatant was collected. Supernatant samples were concentrated by Amicon Ultra 3K centrifugal filter (Millipore), and their buffer was exchanged for Urea /Thiourea buffer on the same Amicon filter by two washes with the buffer. Then, macrophages were lysed in lysis buffer containing 6 M urea, 2 M thiourea, and 100 mM Tris (pH=8), and lysates were incubated for 15 min at RT in thermomixer (1000rpm). Crude lysates were centrifuged for 30 min at 14,000rpm, and the supernatant was collected. 5 µg (EVs, secretome) or 10 ug (LLC tumors) of protein extract from each sample was reduced with DTT, alkylated with IAA, and subjected to digestion with LysC and trypsin. The samples were quenched with 1% TFA and purified on C18 StageTips (Pierce c18 spin tips 84850). The retrieved peptides were analyzed by mass-spectrometry at the Proteomic Unit in Sheba Medical Center. Purified peptides were separated on EasySpray columns (50-cm-long 0.75 µm 803A PepMap) using a 100 or 165 min water-acetonitrile gradient and were injected into the Q-Exactive HF mass spectrometer (Thermo-Fisher Scientific, Waltham, MA, USA) via the EasySpray ionization source. MS analysis was performed using data-dependent acquisition, with fragmentation of the top 10 proteins from each MS spectrum. MS spectra were acquired with a 60,000 resolution, and MS/MS spectra were acquired with a 15,000 resolution. MS raw files were analyzed with MaxQuant version (1.6.5) and the Andromeda search engine using the Uniprot database (UP000000589 2019). The search engine included oxidation and N-term acetyl as variable modifications and carbamidomethyl as fixed modifications and used an FDR of 0.01 for PSM and protein identification. The label-free quantification (LFQ) algorithm in MaxQuant was used for the relative quantification of proteins. The protein list was filtered to eliminate the identifications from the reverse database, common contaminants, and single peptide identifications. In addition, proteins were filtered based on valid values. After log2 transformation, the missing values were imputed with values that form a normal distribution with a width of 0.3 and a downshift of 1.8 standard deviations of the overall protein intensity distribution.

The mass spectrometry proteomics data have been submitted to the ProteomeXchange Consortium via the PRIDE[8-9] partner repository (https://www.proteomexchange.org/) with the following dataset identifiers: for cMSC EV-depletion secretome: PXD040631. For LLC tumor proteomics: PXD040636. For cMSC-sEVs proteomics: PXD040630. For cellular and extracellular proteome of cMSC-sEVs treated macrophages: PXD044593.

**Enzyme-Linked Immunosorbent Assay (ELISA) on Conditioned Medium and Purified EVs**

To compare cEV-encapsulated and secreted cytokines from MI and sham-MI, we treated isolated EVs with RIPA buffer, and specific cytokines were measured with commercially available kits (FineTest Biotech, Hubei, China) of sandwich ELISAs or multiplex ELISA (Quansys Biosciences, Logan, UT, USA)[1] according to the manufacturer's protocol. OD measurements taken by an Infinite F50 Absorbance Microplate Reader (TECAN, Männedorf, Switzerland).[2] All samples were assayed in duplicates, and cytokines concentration was normalized to total protein concentration in the sample.

**miRNA Extraction and Real-Time PCR of EVs**

To determine whether cMSC-sEV-derived miRs exert tumor-promoting functions, we first analyzed the proteomic profile of LLC tumors from mice with and without HF. On day 30 after tumor inoculation (20 days after MI or sham-MI), tumor tissues were extracted and processed for proteomics analysis. We identified 4,909 tumor proteins and, using the TargetScan tool,[10] we identify 48 genes-encoding proteins with a known miR binding site in their 3'-untranslated region

(3'-UTR) and a total of 70 miRs that can bind to those sites. Then, to filter for miRs with reported involvement in cancer, we used the miRBase tool.[11] Next, we further filtered the list for miR involvement in cardiovascular diseases and found 17 potential tumor-promoting miRs.[12] Total RNA was extracted from the purified EVs using the Total EV RNA and Protein Isolation kit (Invitrogen, Carlsbad, CA, USA). The kit contains a strong chaotropic lysis buffer that denatures all proteins and protects the RNA from degradation. Due to the denaturation of the sample, neither the argonaut protein nor any other protein will be bound to the miRNAs upon extraction. cDNA was synthesized from 2 ng of total RNA using the qScript micro-RNA cDNA Synthesis kit (Quanta bio, MA, USA). Quantitative real-time PCR was performed using the PerfeCta SYBR Green kit (Quanta bio, MA, USA). All samples were assayed in duplicates, CT values were normalized to RNU6, and relative micro-RNA expression was calculated with the $2^{-\Delta CT}$ method**.**

#### Culture of Cancer Cells

Mouse LLC cell line was grown in T75 flasks (Thermo-Fisher Scientific, Waltham, MA, USA) with 12 mL of RPMI 1640 (Biological Industries, Beit Haemek, Israel) supplemented with 1% penicillin-streptomycin (Biological Industries, Beit Haemek, Israel), 1% glutamine (Biological Industries, Beit Haemek, Israel), 10% FBS (Gibco, Grand Island, New York). Cells were not allowed to reach 80% confluence and were not grown in vitro for more than 2 weeks. 4T1 and EO771 triple-negative breast cancer cells were grown in T75 flasks with 12 mL of RPMI 1640 (Biological Industries, Beit Haemek, Israel) supplemented with 1% penicillin-streptomycin (Biological Industries, Beit Haemek, Israel), 1% glutamine (Biological Industries, Beit Haemek, Israel), 10% FBS (Gibco, Grand Island, New York), 10mM HEPES buffer (Sigma-Aldrich, St. Louis, MO, USA), 1% sodium pyruvate (Biological Industries, Beit Haemek, Israel) and 0.5% glucose (Sigma-Aldrich, St. Louis, MO, USA). MC38 colon cancer cells and B16 melanoma cells were grown in T75 flasks with 12 mL of DMEM (Gibco, Grand Island, New York) with 10% FBS

(Gibco, Grand Island, New York), 1% penicillin-streptomycin (Biological Industries, Beit Haemek, Israel), 1% L-glutamine (Biological Industries, Beit Haemek, Israel). Cells were not allowed to reach 80% confluence and were not grown in vitro for more than 2 weeks.

**Culture of C166 Mouse Endothelial Cells**

C166 mouse endothelial cells were grown in T75 flasks with 12 mL of DMEM (Gibco, Grand Island, New York) with 10% FBS (Gibco, Grand Island, New York), 1% penicillin-streptomycin (Biological Industries, Beit Haemek, Israel). Cells were not allowed to reach 80% confluence and were not grown in vitro for more than 2 weeks.

**Isolation and Characterization of Primary Lung Fibroblasts**

Primary lung fibroblasts were isolated from healthy lung tissue of naïve, 12-week-old, female C57BL/6 mice, using an enzymatic digestion as previously described.[2] In brief, mice were euthanized, and the lungs were extracted and chopped using a scalpel. Cells were extracted with an enzymatic digestion mixture similar to the extraction of cMSCs. cells were allowed to grow to 85% confluence before each passage up to passage 2.

To characterize the cells used for cEVs collection and isolation, we used flow cytometry (Cytoflex, BECKMAN COULTER, Pasadena, CA, USA) after staining the cells for fibroblasts markers: COL1α (Collagen 1 Polyclonal Antibody, FITC Conjugated, Bioss, MA, USA) and isotype control (Rabbit IgG Isotype Control, FITC Conjugated, Bioss, MA, USA). CD90.2 (FITC anti-mouse CD90.2 (Thy1.2) Antibody, Biolegend, San Diego, CA, USA) and isotype control (FITC Rat IgG2b, κ Isotype Ctrl Antibody, Biolegend, San Diego, CA, USA). mEF-SK4 (Feeder Cells Antibody, anti-mouse, PE, Miltenyi Biotec, Germany) and isotype control (Isotype Control

Antibody, rat IgG1, PE, Miltenyi Biotec, Germany). Macrophages marker: F4/80 (Alexa Fluor® 647 anti-mouse F4/80 Antibody, Biolegend, San Diego, CA, USA) and isotype control (Alexa Fluor® 647 Rat IgG2a, κ Isotype Ctrl Antibody, Biolegend, San Diego, CA, USA), and the endothelial cells marker CD31 (Alexa Fluor® 647 anti-mouse CD31 Antibody, Biolegend, San Diego, CA, USA) and isotype control (Alexa Fluor® 647 Rat IgG2a, κ Isotype Ctrl Antibody, Biolegend, San Diego, CA, USA). The staining protocol and analysis of flow cytometry data were similar to the characterization of cMSCs.

**Activation and Characterization of Macrophages**

To determine the effect of cMSC-sEVs from the failing heart on macrophage polarization, we cultured peritoneal macrophages isolated by the method of resistance to trypsinization[13] and incubated them with cMSC-sEVs either from HF, sham-MI, or saline. Briefly, the peritoneal cavity of 12-week-old female mice was washed with 5 mL PBS. Then, peritoneal fluid was extracted and centrifuged at 500g for 5 minutes, 4°C, and the cells were resuspended in DMEM (Biological Industries, Beit Haemek, Israel) supplemented with 10% fetal bovine serum (Gibco, Grand Island, New York) and 1% penicillin-streptomycin (Biological Industries, Beit Haemek, Israel). The cells were seeded in a 24-well plate (2,000,000 cells in a well) and allowed to adhere for 16 hours, washed with a complete medium, and allowed to recover for 6 hours. Then, the wells were washed with PBS, and the cells were incubated with 1 mL of trypsin-EDTA solution (Gibco, Grand Island, New York) for 5 minutes at 37°C. After trypsinization, we discarded the fluid and applied a complete growth medium to adherent cells. The cells were incubated again with a complete medium for 20 hours, washed twice with serum-free medium, and incubated for 24 hours with cMSC-sEVs from MI, sham-MI, or saline ($10^9$ EVs/ mL). Following 24 hours of incubation with EVs, the wells were washed twice with serum-free medium, and the macrophages were incubated again with serum-free medium for another 24 hours. Then, we collected the conditioned medium and extracted total RNA from the macrophages. To determine the purity of isolated macrophages, we fixed and stained the cells for F4/80 and DAPI using a rat anti-mouse F4/80 primary antibody (Bio-RAD, Hercules, CA, USA) and a goat anti-rat conjugated to Alexa fluor 488 secondary antibody (Jackson ImmunoResearch Laboratories, West Grove, PA, USA). Next, we imaged the cells using an LSM 700 confocal microscope (ZEISS, Jena, Germany). The purity of macrophages was determined by the ratio of F4/80 positive cells to the total number of nuclei.

**Total RNA Extraction and rtPCR of Macrophages**

Peritoneal macrophages were isolated and incubated with cMSC-sEVs as described above. For total RNA extraction, we used the Extracta Plus RNA extraction kit (Quanta bio, MA, USA), and for cDNA synthesis, we used the qScript cDNA Synthesis kit (Quanta bio, MA, USA). Quantitative real-time PCR was performed using PerfeCTa SYBR Green FastMix, HIGH ROX (Quanta bio, MA, USA) with 5 ng of equivalent cDNA. All samples were assayed in duplicates, CT values were normalized to GAPDH, and relative gene expression was calculated with the $2^{-\Delta CT}$ method**.**

**Total Nitrite and Nitrate Concentration Assay**

To measure the activity of nitric oxide synthase in macrophages incubated with cMSC-sEVs, we used Total nitrite and nitrate concentration assay (Cayman Chemical, Michigan, USA) according to the manufacturer’s protocol. All samples were assayed in duplicates.

#### Scratch Migration Assay

To assess the effect of cMSC-sEVs or condition medium of macrophages incubated with cMSC-sEVs, on cancer cells, lung fibroblasts, and endothelial cells, we used a migration (“scratch”) assay. Cells were cultured in a 96-well plate until full confluence, and the medium was replaced with serum-free conditions. Cells were scratched with a sterile 10-µl pipette tip, washed, and incubated with increased concentrations of cMSC-sEVs from the infarcted or healthy heart up to $10^9$ EVs/mL, or with PBS as control. For incubation with condition medium from macrophages treated with cMSC-sEVs, we used 5 µg of total protein and DMEM medium as control. Then, we acquired images of the scratched area every 4 hours using an LSM 700 confocal microscope (ZEISS, Jena, Germany) with a special incubation chamber set to 37°C and 5% $CO_2$. All samples were assayed in duplicates, and scratch area was measured using ImageJ software version 1.54e.[14]

#### Proliferation Assay

To investigate if Post-MI HF cMSC-sEV or condition medium of macrophages incubated with cMSC-sEVs, facilitates cancer cells or lung fibroblasts proliferation, we used a colorimetric assay based on 3-(4,5-dimethylthiazol-2-yl)-5-(3-carboxymethoxyphenyl)-2-(4-sulfophenyl)-2H-tetrazolium (MTS) (Abcam, Cambridge, UK). Briefly, $40 \cdot 10^3$ LLC, MC38 cells or $10 \cdot 10^3$ 4T1, EO771, B16 cells or primary lung fibroblasts were seeded into a 96-well plate and incubated in full growth medium for 20h. Then, the medium was replaced to serum-free condition for 4 hours, washed, and the cells were incubated with increased concentrations of cMSC-sEVs from the infarcted or healthy heart up to $10^9$ EVs/mL or with PBS as control. For incubation with condition medium from macrophages treated with cMSC-sEVs, we used 5 µg of total protein and DMEM medium as control. After 0, 24, or 48 hours of incubation, the wells were washed with 4.5% glucose in PBS to minimize the medium effect on the assay,[15] and incubated for 2 hours In 10% MTS solution (Abcam, Cambridge, UK) according to the manufacturer's protocol. For

$OD_{490nm}$ measurements, we used an Infinite F50 Absorbance Microplate Reader (TECAN, Männedorf, Switzerland).[2] All samples were assayed in duplicates, and viable cell number was calculated by linear regression with a calibration curve.

#### Angiogenesis (Tube) Formation Assay

To assess the pro-angiogenic properties of conditioned medium from cMSC-EV-treated macrophages, we conducted a tube formation assay. C166 endothelial cells (ATCC, VA, USA) stained with CellTrackerTM Red CMTPX Dye (Invitrogen, MA, USA) according to the manufacturer's protocol. Then, 20,000 cells were seeded into a 96-well plate coated with CellMatrix Basement Membrane Gel (ATCC, VA, USA) in the presence of conditioned medium from macrophages (5 µg protein). Then, we acquired images of polygons after 8 hours, using an LSM 700 confocal microscope (ZEISS, Jena, Germany) with a special incubation chamber set to 37°C and 5% $CO_2$. All samples were assayed in duplicates, and the scratch area was measured using ImageJ software version 1.54e.

#### Endothelial Permeability Assay

To determine if cMSC-EVs from failing hearts promote endothelial dysfunction and permeability, we used a permeability (diffusion) assay. Briefly, we coated transwell inserts (1-µm pore size, Sigma-Aldrich, St. Louis, MO, USA) with fibronectin (Sigma-Aldrich, St. Louis, MO, USA) and then seeded C166 endothelial cells (ATCC, VA, USA). The cells were allowed to form a monolayer. Then, the culture medium was replaced with a serum-free medium, and the cells were incubated with cMSC-EVs for 24 hours. After 24 hours, we added horseradish peroxidase (HRP, 0.288µM, Sigma-Aldrich, St. Louis, MO, USA) as a test molecule that we can detect after diffusing to the lower chamber of the transwell. Then, we measured the concentration of HRP in the lower chambers after 0, 0.5, and 2 hours using TMB substrate (Biolegend, San Diego, CA, USA) according to manufacturer instructions.[16]

#### In Vivo Biodistribution and Uptake of cMSC-EVs

For tracking of labeled EVs, we labeled 100 µg of cMSC-EVs' protein in each reaction with the ExoGlow vivo EV labeling kit (system biosciences, Palo Alto, CA) according to the manufacturer's protocol. Then, we washed the labeled EVs 3 times by ultrafiltration with AMICON 15 10-kDa tubes (Millipore, Burlington, Massachusetts). For control, we used the same amount of dye in PBS with the same protocol. The labeled EVs (20 µg per mouse) were immediately injected into the left ventricular cavity under echocardiography guidance, and we monitored their biodistribution using IVIS Lumina LT series III (PerkinElmer, MA, USA) at 1, 12, 24 hours. After 24 hours, we euthanized the mice and examined the heart, lungs, liver, kidneys, spleen, femur, sternum, and tumor ex vivo. For quantitative analysis of the average radiance, we used Living Image version 4.7.2 (PerkinElmer, MA, USA).

#### Adoptive Transfer of cMSC-sEVs

MI or sham-MI was induced in donor mice as described above**.** We isolated cMSCs 10 days after MI, grew them in vitro, and purified the sEVs that they shed to the medium. EV dosing was determined by protein concentration (2 µg of EV protein per injection), and EV count was validated with NTA. Recipient mice received 3 injections of MI cMSC-sEVs, sham-MI cMSC-sEVs, or saline, to the subcutaneous tissue at the inoculation site, before inoculation. We inoculated LLC cells (750,000 cells in 100 µL PBS) to the hindlimb of the mice 7 days after the first injection, and tumor development was evaluated daily by visualization and palpation. On day 5, we resumed the injections (MI cMSC-sEVs, sham-MI cMSC-sEVs or saline) every 48 hours to the subcutaneous tissue adjacent to the tumor and monitored tumor growth with ultrasound as previously described.

**EV Depletion**

For EV inhibition in vivo, we used GW4869 (Sigma-Aldrich, St. Louis, MO, USA), an inhibitor for neutral sphingomyelin phosphodiesterase (N-SMase), the enzyme that converts membrane sphingomyelin into ceramide, which is required for EV formation. We dissolved GW4869 in Dimethyl Sulfoxide (DMSO, Sigma-Aldrich, St. Louis, MO, USA) to a stock concentration of 0.5 mg/mL and kept it -20°C. A stock solution was used within two weeks of dissolution. We delivered the GW4869 or DMSO (control), diluted to 10% in PBS (v/v), by intraperitoneal injection every 2 days (2.5 mg/kg), starting 3 days after MI and throughout the follow-up period.[17-19]

**Adoptive Transfer of cMSC-sEVs During Systemic EV Depletion**

For EV depletion, we used GW4869 as described above. We generated 2 models of cMSC-EV transfer during systemic EV depletion: a heterotopic lung cancer model and an orthotopic lung cancer model as described above. For adoptive transfer, we isolated cMSC-EVs from donor mice 10 days after MI or sham-MI as described above.

For the heterotopic model, LLC lung cancer cells were inoculated 10 days before MI, and we started the administration of GW4869 (2.5 mg/kg, IP) and cMSC-EVs from HF donor mice (2μg of EV protein per injection, SC) three days after MI, every 48 hours until the end of the follow-up period.

For the orthotopic model, we started administration of GW4869 (2.5 mg/kg, IP) on day 3, every 48 hours, and cMSC-EVs from either HF or sham-MI mice at day 4 (10 μg of EV protein per injection, IP) every 48 hours until the end of the follow-up period. In this orthotopic model, the mortality of mice was 27% (3/11) for the group that received GW4869 vehicle (DMSO) and EV vehicle (PBS). The mortality for the other groups was 18% (2/11). LLC lung cancer cells were injected into the tail vein of mice at day 10 after MI.

**In Vivo Imaging of Lung Tumors**

To monitor orthotopic lung cancer, we used IVIS Lumina LT series III system (PerkinElmer, MA, USA) for the detection of a signal emitted from luciferase-expressing LLC cells. For each measurement, a total of 150 mg/kg luciferin (Promega, WI, USA) was injected intraperitoneally 15 minutes before the signal was measured. For analysis, we used Living Image software version 4.7.2 (PerkinElmer, MA, USA).

**Micro-CT Imaging of Lung Tumors**

To determine the number of lung tumors and the total tumor area in the lungs (tumor burden), we used micro-CT (SkyScan 1176, Bruker Biopsin, MA, USA).[20] We scanned the mice under 2% isoflurane anesthesia at day 30 using the following settings: nominal resolution of 35 μm, a 0.2 mm aluminum filter, a tube voltage of 45 kV, Source Current (uA)=556, Rotation Step (deg)=0.700. For reconstruction, we used NRecon software version 1.7.4.6 (Bruker Biopsin, MA, UAS) with a modified Feldkamp algorithm64 accelerated by GPU. Ring artifact reduction, Gaussian smoothing (3%), and beam hardening correction (20%) were applied. Then, the number of tumors and tumor burden were measured by a technician (MN) who was blinded to the allocation of mice to the various treatments. Each tumor mass in the lung was identified in the axial plane and discriminated from vessels, and confirmed by both sagittal and coronal planes.

##### In Vivo Tracking of Glucose-Coated Gold Nano-Particles

To evaluate both spreading and metabolism of LLC tumor in the lungs during systemic EV depletion, we used glucose-coated gold nanoparticles (GNPs). GNPs are specifically taken by cells with high glucose metabolism, such as tumor cells.[20] GNPs were synthesized as previously described.[20] In brief, to synthesize 20 nm particles, a total of 0.414 mL of 1.4 M HAuCl4 solution in 200 mL of water was added to a 250 mL single-neck round-bottom flask. The solution was stirred in an oil bath on a hot plate until it boiled, then 4.04 mL of a 10% sodium citrate solution of 0.39 M sodium citrate tribasic dihydrate 98% (Sigma-Aldrich, St. Louis, MO, USA) was added and stirred for 5 additional minutes. The flask was then removed from the hot oil. To prevent aggregation and stabilize the particles in physiological solutions, SH-PEG-COOH 1KDa (Sigma-Aldrich, St. Louis, MO, USA) was absorbed onto the GNPs. First, the solution was centrifuged to dispose of excess citrate. SH-PEG-COOH 1KDa solution ($2.26 \times 10^{-3}$ g) was then added to the GNP solution, and the mixture was stirred overnight and subsequently centrifuged. Next, excess EDC (N-ethyl-N-(3-(dimethylamino)- propyl) carbodiimide ($1.87 \times 10^{-3}$ g) and NHS (N-hydroxysuccinimide) (Thermo-Fisher Scientific, Waltham, MA, USA) ($2.12 \times 10^{-3}$ g) were added to the solution, followed by the addition of $1.75 \times 10^{3}$ g glucose-2 (2GF): D-(β)-glucosamine hydrochloride, (Sigma-Aldrich, St. Louis, MO, USA). NHS and EDC form an active ester intermediate with the −COOH functional groups, which can then undergo an amidation reaction with the glucose $NH_2$ group. 3 mg glucosamine molecule C-2 (2GF-GNP): D-(β)-glucosamine hydrochloride $1.75 \times 10^{3}$ g was added to the activated linker-coated GNPs.

Next, we scanned the mice using micro-CT as described above, and 3-4 mice from each group were randomly chosen to receive an injection of 6 mg of GNPs into the tail vein; after 3 hours the mice were euthanized and perfused with 20 mL saline by inferior vena cava cannulation. Then, we scan the mice again using micro-CT to track the GNPs in the chest and lungs. Enhancement of tumor tissue by GNPs was quantified by analysis of the mean gray value (0-255) in the micro-CT scans of tumor tissue in the lungs. Finally, to quantify the exact among of gold in the lungs, we used inductively coupled plasma spectrometry (ICP-OES 710, Agilent Technologies, CA, USA). Lung tissues were melted with 1 mL of aqua regia acid and then evaporated and diluted to a total volume of 4 mL. After filtration of the samples, gold concentrations were determined according to absorbance values, with correlation to calibration curves, constructed from solutions with known gold concentrations (0, 0.5, 2, and 5 mg/L). The detection limit of ICP is 50 ppb.

**Anti-Remodeling and HF therapy with Spironolactone**

MI was induced as described above. LLC cancer cells were inoculated heterotopically to the subcutis of the right hindlimb of female C57BL/6 mice (750,000 cells in 100 µL PBS), 10 days before MI or sham-MI. Ultrasound measurements of the tumor using a small animal echocardiography system were performed every 3 days. For anti-remodeling therapy, we used spironolactone (Sigma-Aldrich, St. Louis, MO, USA) dissolved in Dimethyl Sulfoxide (DMSO, Sigma-Aldrich, St. Louis, MO, USA) to a stock concentration of 40 mg/mL. Then, we diluted the working solution 1:3 in corn oil. We injected the working solution of spironolactone (50 mg/kg) or vehicle to the subcutis of the mice. Spironolactone was dissolved in DMSO and, within 30 minutes, was injected subcutaneously into the back of the neck of mice. On day 29, mice were euthanized with an isoflurane overdose.

#### Lung Histology

To evaluate lung congestion after MI and HF, we stained lung sections from mice with either MI or sham-MI, hematoxylin and eosin. Briefly, the lungs were removed 10 days after MI or sham-MI, washed with PBS, and fixed with 4% PFA for 48 hours at 4°C. Then, the fixed tissue was paraffinized, embedded, and sectioned into 5μm slices. After hematoxylin and eosin staining (Sigma-Aldrich, St. Louis, MO, USA), sections of each lung were examined by brightfield microscopy.

#### Morphometry of Heart Sections

To further assess LV remodeling, mice were euthanized by excessive inhalation of isoflurane, and the chest opened. Then, we perfused the heart with 5 mL of 14.9% KCl and then 5 mL of 4% formaldehyde. After perfusion, the heart was removed and fixed with 4% PFA for 48 hours, at 4°C. The fixed tissue was then paraffinized, embedded, and sectioned into 5-μm slices. Next, we used picrosirius red (Direct Red 80, Sigma-Aldrich, St. Louis, MO, USA) to detect scarring and fibrosis. Each heart was photographed at the level of the papillary muscles and analyzed with planimetry software (Sigma Scan Pro 5, Systat Software, San Jose, California, USA).

We measured average scar thickness from 3 measurements of scar thickness, average wall thickness from 3 measurements of septum thickness, LV muscle area, LV cavity area, whole LV area, epicardial scar length, and endocardial scar length. Relative scar thickness was calculated as average scar thickness divided by average wall thickness. The expansion index was calculated as [LV cavity area/whole LV area]/relative scar thickness.[13]

#### Ki67 Staining of Tumor Sections

To determine the effect of MI and cEVs on tumor cell proliferation in vivo, we used Ki67 staining. Briefly, the tumors were removed, weighted, washed with PBS, and fixed with 4% PFA for 48 hours, 4°C. The fixed tissue was then paraffinized, embedded, and sectioned into 5-μm slices. Then, we rehydrated the slides, used the heat antigen retrieval method, and blocked them with normal donkey serum (Jackson ImmunoResearch Laboratories, West Grove, PA, USA). For staining, we used rabbit anti-mouse Ki67 (Abcam, Cambridge, UK), Biotin-SP AffiniPure Donkey Anti-Rabbit, and Cy3 Streptavidin (Jackson ImmunoResearch Laboratories, West Grove, PA, USA) and DAPI (Sigma-Aldrich, St. Louis, MO, USA). For mounting, we used Lab Vision PermaFluor Aqueous Mounting Medium (Thermo-Fisher Scientific, Waltham, MA, USA). The percentage of $Ki67^+$ cells was calculated by dividing the number of $Ki67^+$ nuclei by the total number of nuclei and multiplied by 100. For each tumor, the percentage of $Ki67^+$ was calculated from 5-10 fields collected randomly from the tumor periphery.

#### Statistical Analysis

Experimental data are expressed as mean ± standard deviation (SD), and specific statistical tests are detailed in the figure legends. Statistical analyses were performed with GraphPad Prism version 9 (GraphPad Software) unless otherwise stated. Statistical analyses of the identification and quantization of the proteomic results were done using Perseus 1.6.7.0 software, and statistical analysis for NTA data was performed using STATA BE version 17.

Two-tailed student's t-test was performed to compare normally distributed continuous variables and a two-tailed Mann-Whitney U test for non-normal distribution (tested by the D'Agostino-Pearson omnibus normality test). When the experiment design included more than one comparison, we used multiple t-tests or multiple Mann-Whitney tests to account for multiple comparisons. For experiments with more than 2 groups, the comparison was performed with one-way or two-way ANOVA with Holm-Šídák's post-test. Repeated measures tests were used if the same subject was sampled at different time points. To compare two frequency distributions (NTA data), we used zero-inflated negative binomial regression.

**Supplementary Table 1. A List of Primers for qRT-PCR**

| Target | Forward primer | Reverse primer |
|---|---|---|
| RNU6 | *GTGCTCGCTTCGGCAGCACATATACTAAAAT*<br>*TGGAACGATACAGAGAAGAT*<br>*TAGCATGGCCCCTGCGCAAGGAT*<br>*GACACGCAAATTCGTGAAGCGTTCCATATTTTT* | Quanta Bio universal primer |
| miR-221 | *AGCUACAUUGUCUGCUGGGUUUC* | Quanta Bio universal primer |
| miR-21 | *UAGCUUAUCAGACUGAUGUUGA* | Quanta Bio universal primer |
| miR-24-1 | *UGGCUCAGUUCAGCAGGAACAG* | Quanta Bio universal primer |
| miR-24-2 | *UGCCUACUGAGCUGAAACAGU* | Quanta Bio universal primer |
| miR-214 | *ACAGCAGGCACAGACAGGCAGU* | Quanta Bio universal primer |
| miR-34a | *UGGCAGUGUCUUAGCUGGUUGU* | Quanta Bio universal primer |
| miR-219-5p | *GAUUGUCCAAACGCAAUUCU* | Quanta Bio universal primer |
| miR-208a-3p | *AUAAGACGAGCAAAAAGCUUGU* | Quanta Bio universal primer |
| miR-208a-5p | *GAGCUUUUGGCCCGGGUUAUAC* | Quanta Bio universal primer |
| GAPDH | *GGTCGGTGTGAACGGATTTGG* | *AGACCATGTAGTTGAGGTCAATGAA* |
| iNOS | *ACCAAGATGGCCTGGAGGAATG* | *GTTCCGAGCGTCAAAGACCT* |
| PD-L1 | *CACATCCTCCACAGAACAGGACT* | *TCTCCACATCTAGCATCCTCACT* |

### Figure S1.

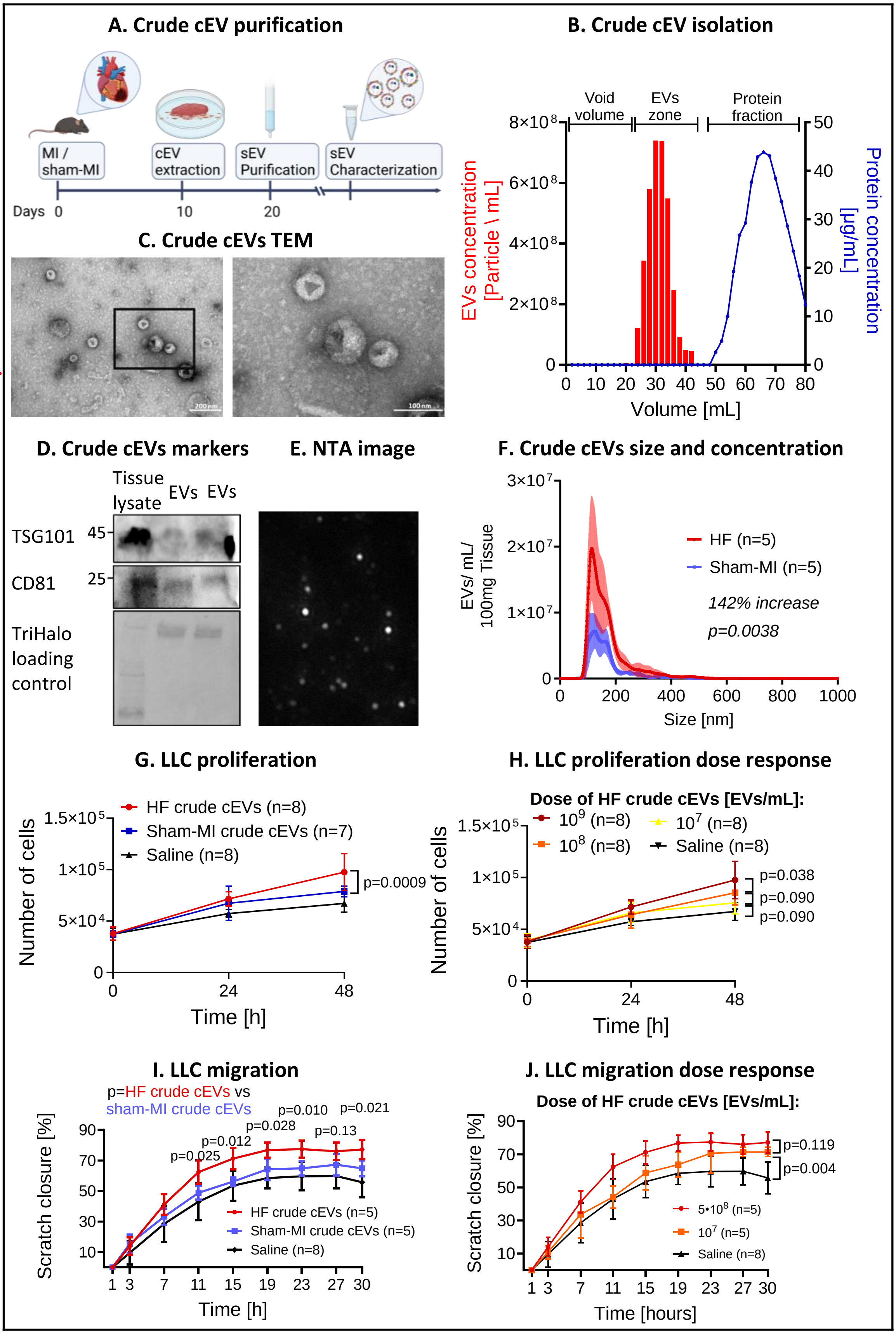
A. Crude cEV purification
MI / sham-MI
cEV extraction
sEV Purification
sEV Characterization
Days 0 10 20
B. Crude cEV isolation
Void volume
EVs zone
Protein fraction
EVs concentration [Particle \ mL]
Protein concentration [µg/mL]
Volume [mL]
C. Crude cEVs TEM
D. Crude cEVs markers
Tissue lysate
EVs
EVs
TSG101
CD81
TriHalo loading control
E. NTA image
F. Crude cEVs size and concentration
HF (n=5)
Sham-MI (n=5)
142% increase
p=0.0038
EVs/ mL/ 100mg Tissue
Size [nm]
G. LLC proliferation
HF crude cEVs (n=8)
Sham-MI crude cEVs (n=7)
Saline (n=8)
p=0.0009
Number of cells
Time [h]
H. LLC proliferation dose response
Dose of HF crude cEVs [EVs/mL]:
10^9 (n=8)
10^7 (n=8)
10^8 (n=8)
Saline (n=8)
p=0.038
p=0.090
p=0.090
Number of cells
Time [h]
I. LLC migration
p=HF crude cEVs vs sham-MI crude cEVs
p=0.025
p=0.012
p=0.028
p=0.010
p=0.021
p=0.13
HF crude cEVs (n=5)
Sham-MI crude cEVs (n=5)
Saline (n=8)
Scratch closure [%]
Time [h]
J. LLC migration dose response
Dose of HF crude cEVs [EVs/mL]:
p=0.119
p=0.004
5•10^8 (n=5)
10^7 (n=5)
Saline (n=8)
Scratch closure [%]
Time [hours]

### Figure S2.

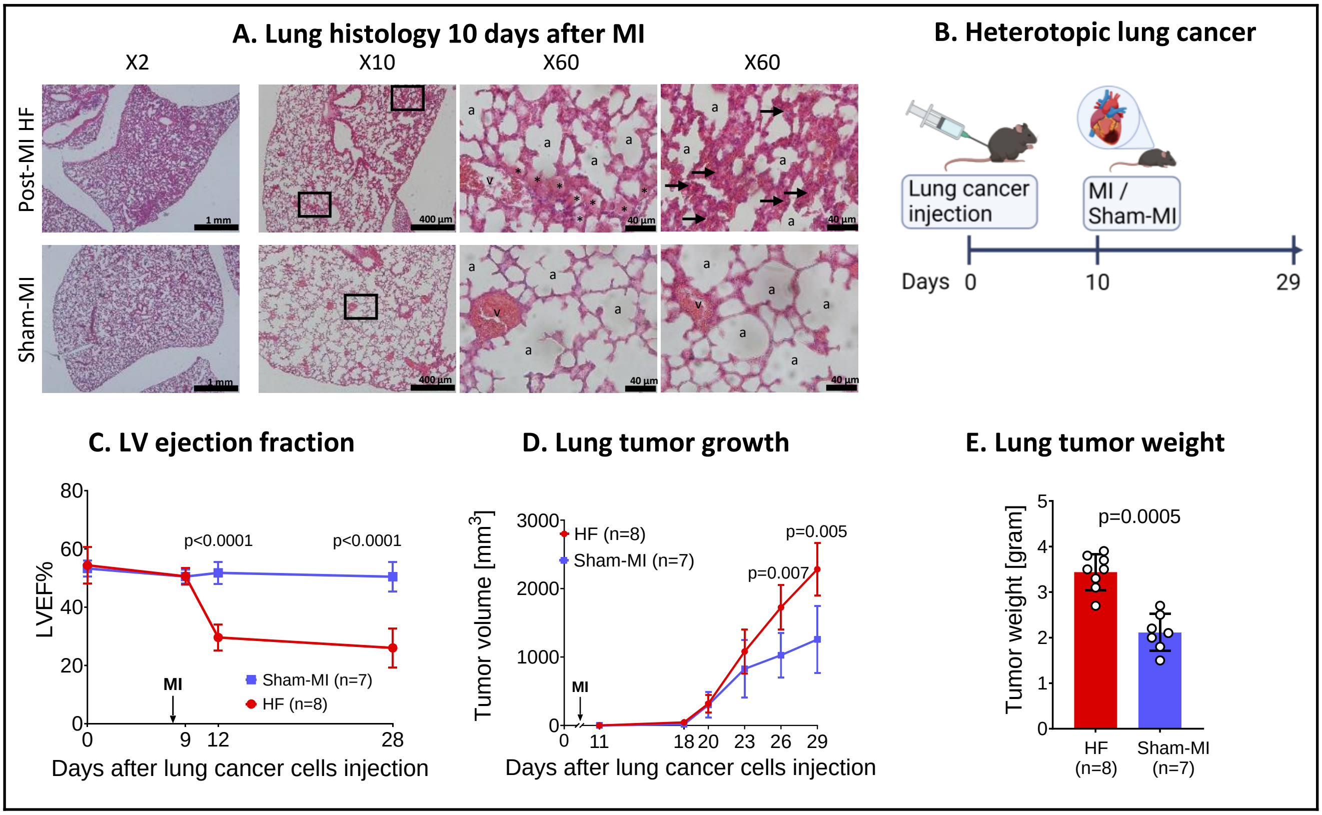
A. Lung histology 10 days after MI
X2
X10
X60
X60
Post-MI HF
Sham-MI
1 mm
400 µm
40 µm
B. Heterotopic lung cancer
Lung cancer injection
MI / Sham-MI
Days 0
10
29
C. LV ejection fraction
LVEF%
p<0.0001
p<0.0001
MI
Sham-MI (n=7)
HF (n=8)
Days after lung cancer cells injection
D. Lung tumor growth
Tumor volume [mm3]
HF (n=8)
Sham-MI (n=7)
p=0.005
p=0.007
MI
Days after lung cancer cells injection
E. Lung tumor weight
Tumor weight [gram]
p=0.0005
HF (n=8)
Sham-MI (n=7)

#### Figure S3.

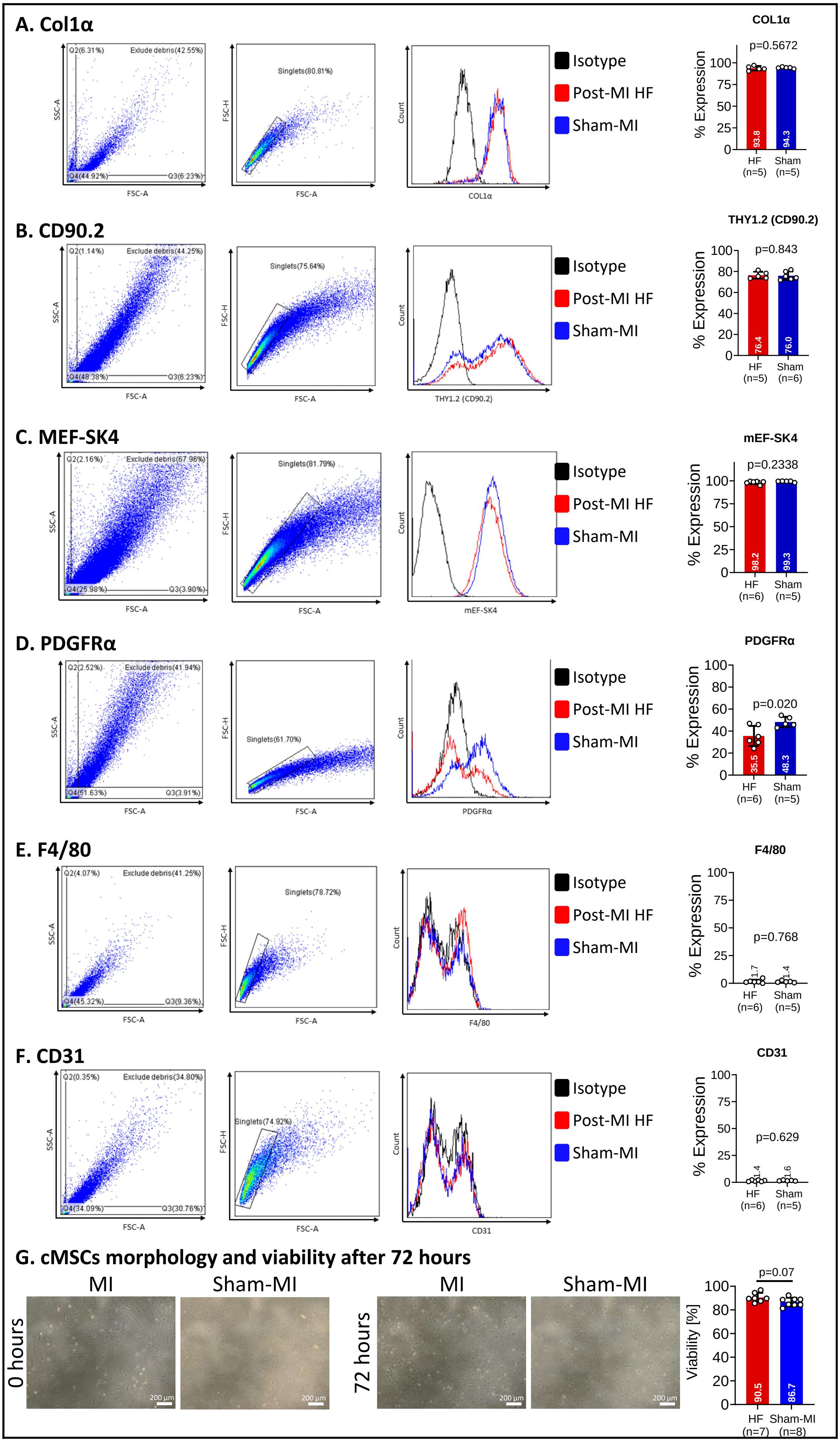

A. Col1α
B. CD90.2
C. MEF-SK4
D. PDGFRα
E. F4/80
F. CD31
G. cMSCs morphology and viability after 72 hours
Isotype
Post-MI HF
Sham-MI
COL1α
p=0.5672
THY1.2 (CD90.2)
p=0.843
mEF-SK4
p=0.2338
PDGFRα
p=0.020
F4/80
p=0.768
CD31
p=0.629
p=0.07
% Expression
Viability [%]
HF
Sham
Sham-MI
MI
0 hours
72 hours

### Figure S4.

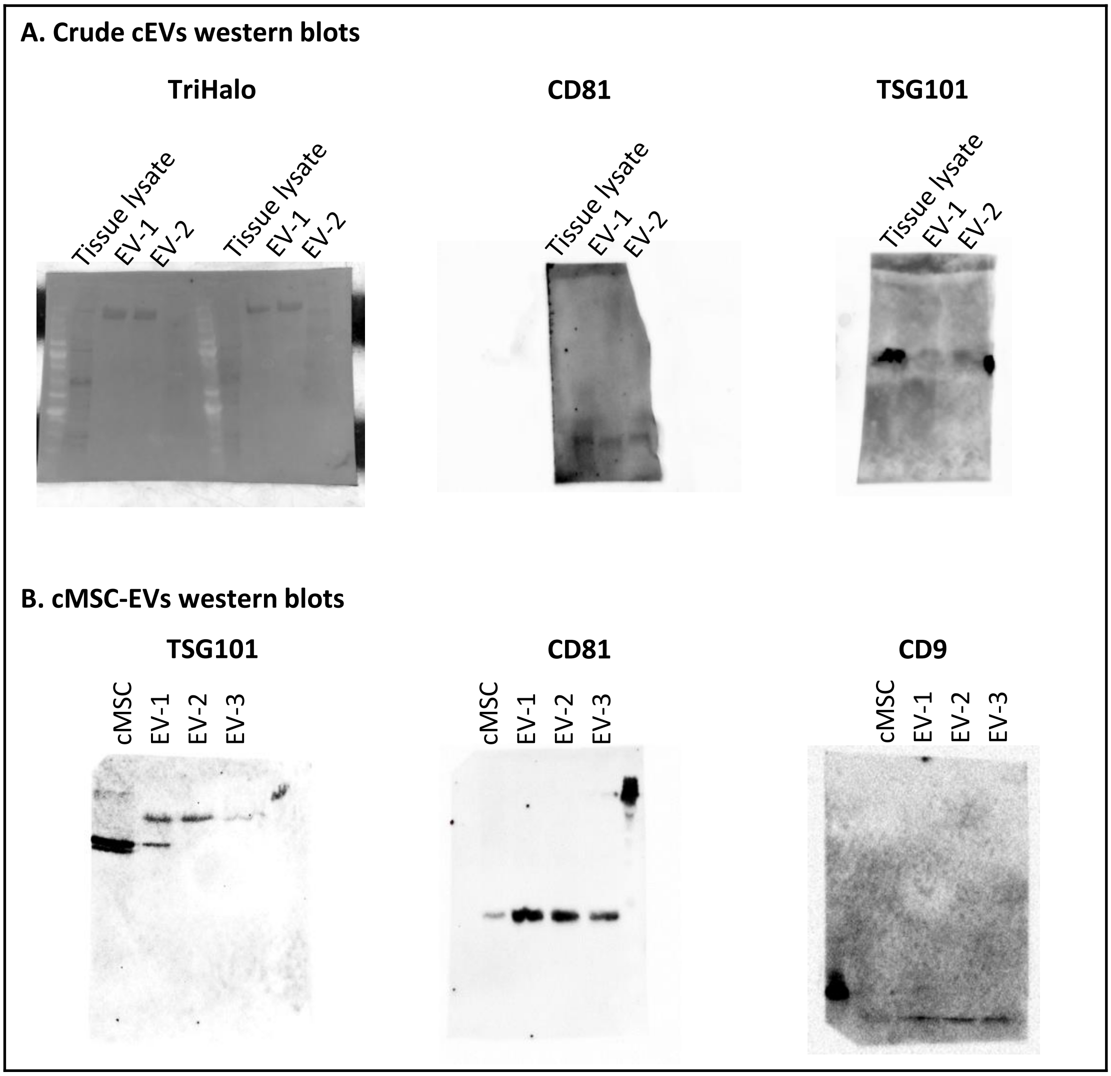
A. Crude cEVs western blots
TriHalo
CD81
TSG101
Tissue lysate
EV-1
EV-2
Tissue lysate
EV-1
EV-2
Tissue lysate
EV-1
EV-2
Tissue lysate
EV-1
EV-2
B. cMSC-EVs western blots
TSG101
CD81
CD9
cMSC
EV-1
EV-2
EV-3
cMSC
EV-1
EV-2
EV-3
cMSC
EV-1
EV-2
EV-3

### Figure S5.

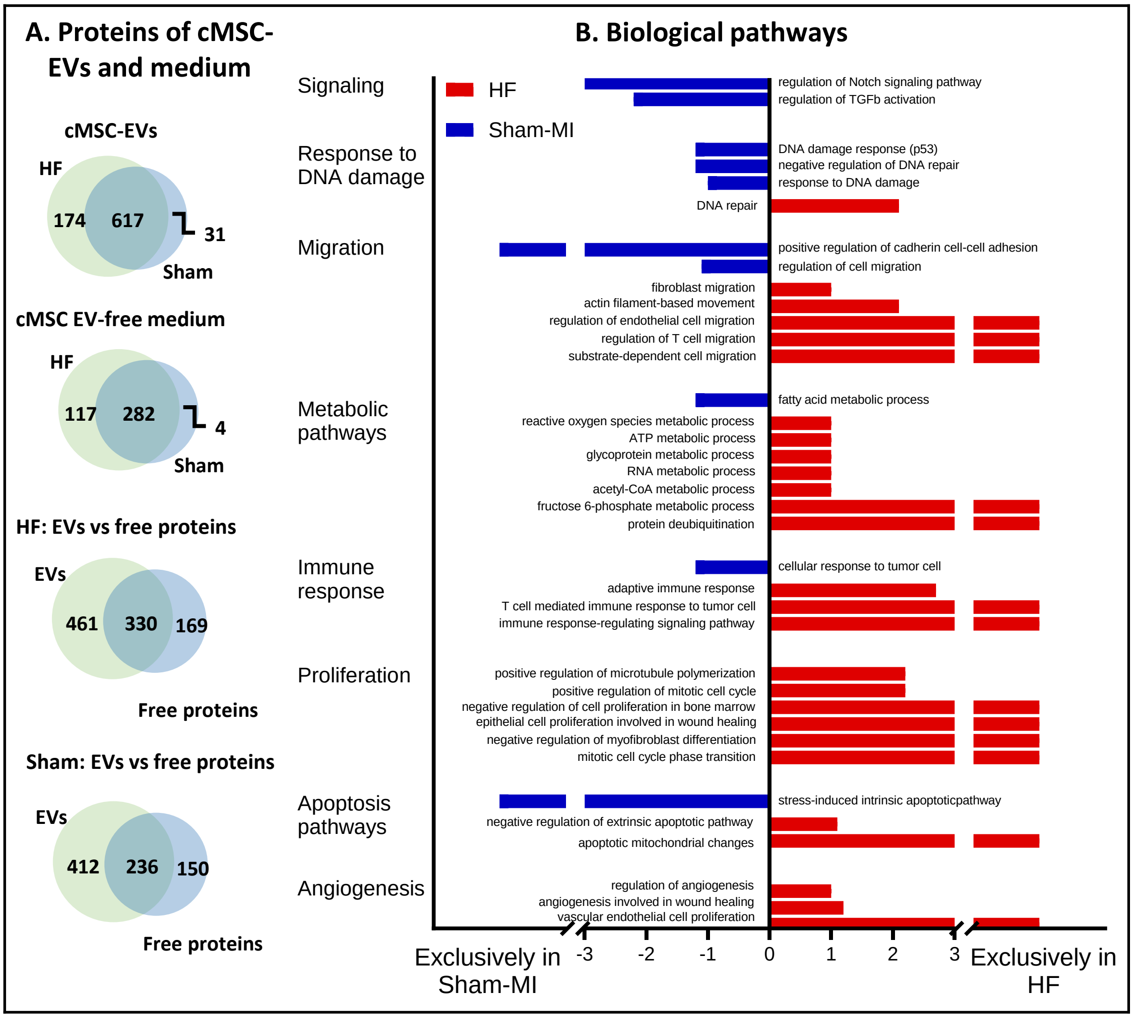

A. Proteins of cMSC-EVs and medium
cMSC-EVs
HF
174
617
31
Sham
cMSC EV-free medium
HF
117
282
4
Sham
HF: EVs vs free proteins
EVs
461
330
169
Free proteins
Sham: EVs vs free proteins
EVs
412
236
150
Free proteins
B. Biological pathways
HF
Sham-MI
Signaling
regulation of Notch signaling pathway
regulation of TGFb activation
Response to DNA damage
DNA damage response (p53)
negative regulation of DNA repair
response to DNA damage
DNA repair
Migration
positive regulation of cadherin cell-cell adhesion
regulation of cell migration
fibroblast migration
actin filament-based movement
regulation of endothelial cell migration
regulation of T cell migration
substrate-dependent cell migration
Metabolic pathways
fatty acid metabolic process
reactive oxygen species metabolic process
ATP metabolic process
glycoprotein metabolic process
RNA metabolic process
acetyl-CoA metabolic process
fructose 6-phosphate metabolic process
protein deubiquitination
Immune response
cellular response to tumor cell
adaptive immune response
T cell mediated immune response to tumor cell
immune response-regulating signaling pathway
Proliferation
positive regulation of microtubule polymerization
positive regulation of mitotic cell cycle
negative regulation of cell proliferation in bone marrow
epithelial cell proliferation involved in wound healing
negative regulation of myofibroblast differentiation
mitotic cell cycle phase transition
Apoptosis pathways
stress-induced intrinsic apoptoticpathway
negative regulation of extrinsic apoptotic pathway
apoptotic mitochondrial changes
Angiogenesis
regulation of angiogenesis
angiogenesis involved in wound healing
vascular endothelial cell proliferation
Exclusively in Sham-MI
-3
-2
-1
0
1
2
3
Exclusively in HF

### Figure S6.

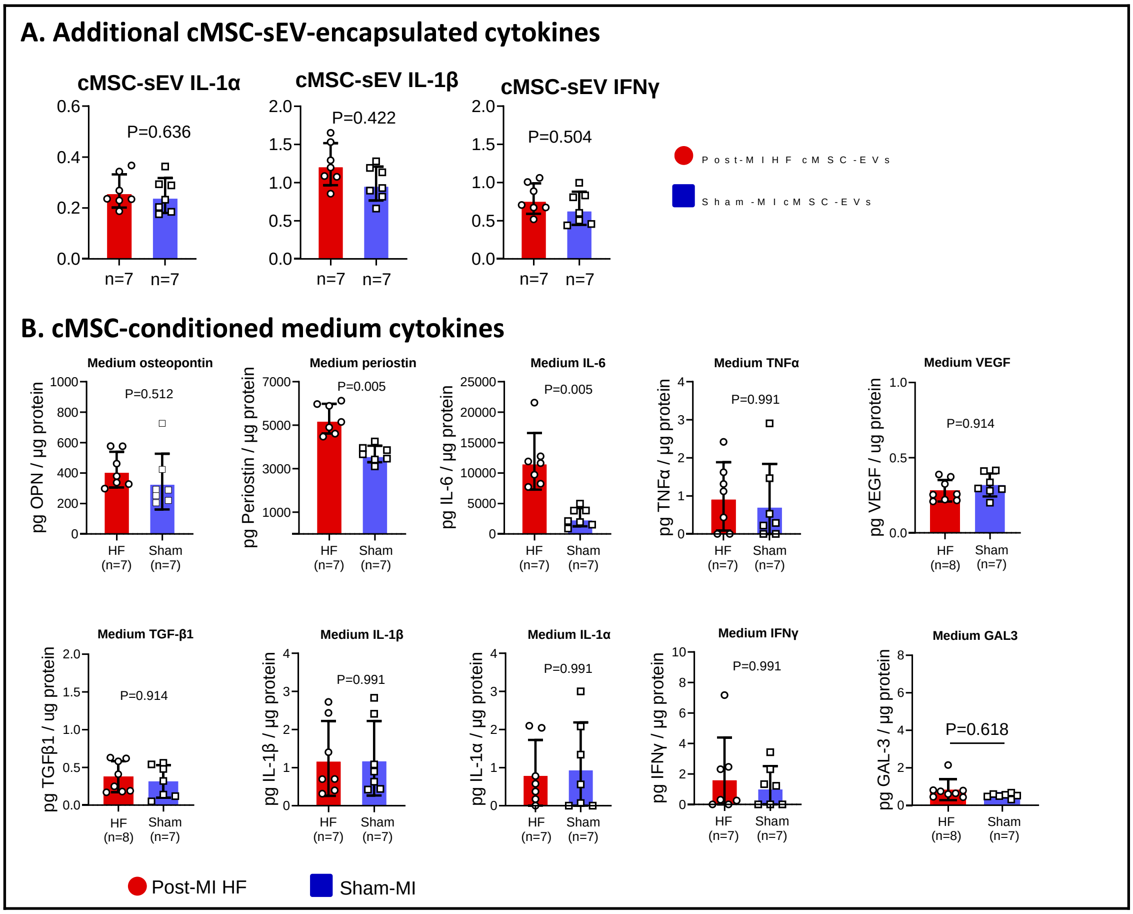
A. Additional cMSC-sEV-encapsulated cytokines
cMSC-sEV IL-1α
P=0.636
cMSC-sEV IL-1β
P=0.422
cMSC-sEV IFNγ
P=0.504
n=7 n=7
Post-MI HF cMSC-EVs
Sham-MI cMSC-EVs
B. cMSC-conditioned medium cytokines
Medium osteopontin
P=0.512
pg OPN / µg protein
Medium periostin
P=0.005
pg Periostin / µg protein
Medium IL-6
P=0.005
pg IL-6 / µg protein
Medium TNFα
P=0.991
pg TNFα / µg protein
Medium VEGF
P=0.914
pg VEGF / ug protein
Medium TGF-β1
P=0.914
pg TGFβ1 / ug protein
Medium IL-1β
P=0.991
pg IL-1β / µg protein
Medium IL-1α
P=0.991
pg IL-1α / µg protein
Medium IFNγ
P=0.991
pg IFNγ / µg protein
Medium GAL3
P=0.618
pg GAL-3 / µg protein
HF (n=7) Sham (n=7)
HF (n=8) Sham (n=7)
Post-MI HF
Sham-MI

### Figure S7.

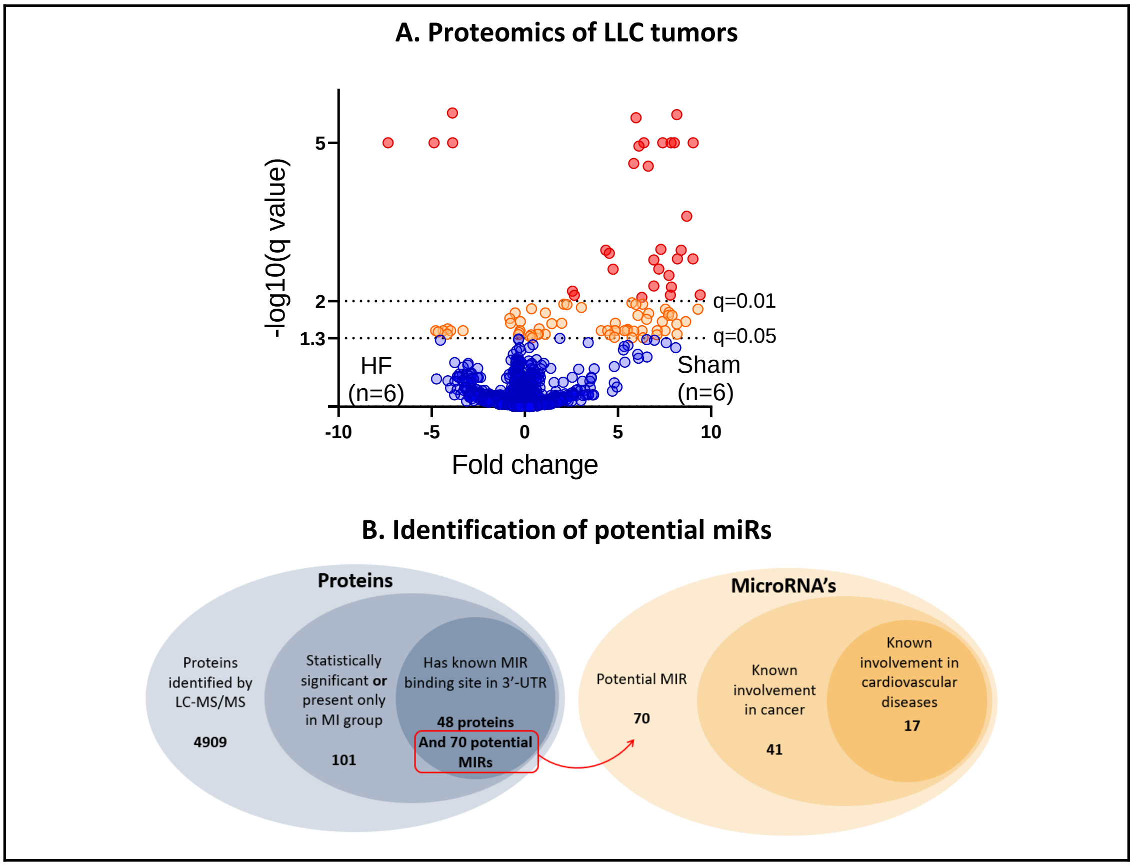
A. Proteomics of LLC tumors
-log10(q value)
5
2
1.3
q=0.01
q=0.05
HF
(n=6)
Sham
(n=6)
-10
-5
0
5
10
Fold change
B. Identification of potential miRs
Proteins
Proteins identified by LC-MS/MS
4909
Statistically significant or present only in MI group
101
Has known MIR binding site in 3'-UTR
48 proteins
And 70 potential MIRs
MicroRNA's
Potential MIR
70
Known involvement in cancer
41
Known involvement in cardiovascular diseases
17

### Figure S8.

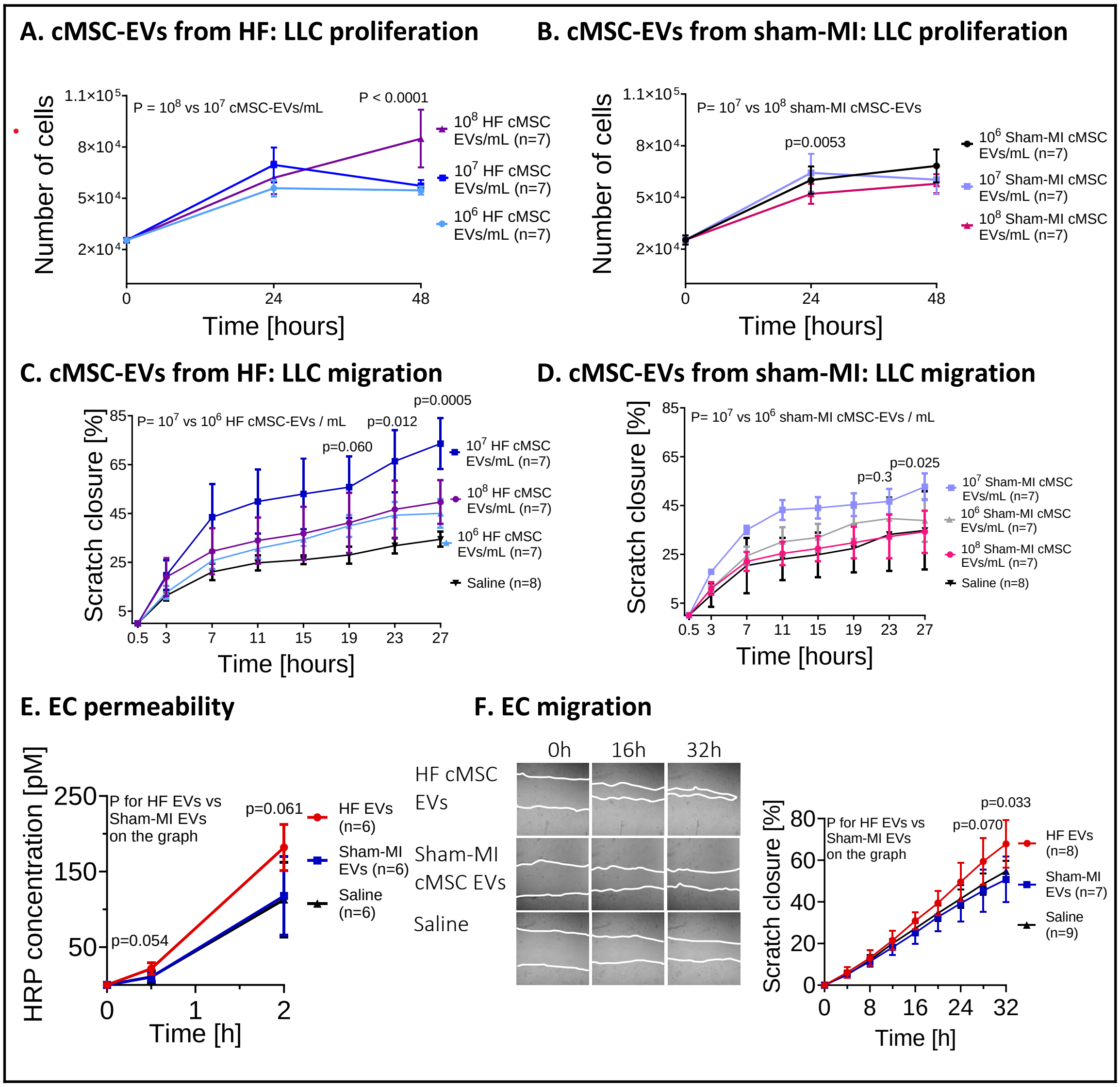
A. cMSC-EVs from HF: LLC proliferation
B. cMSC-EVs from sham-MI: LLC proliferation
Number of cells
Time [hours]
P = 10^8 vs 10^7 cMSC-EVs/mL
P < 0.0001
10^8 HF cMSC EVs/mL (n=7)
10^7 HF cMSC EVs/mL (n=7)
10^6 HF cMSC EVs/mL (n=7)
P= 10^7 vs 10^8 sham-MI cMSC-EVs
p=0.0053
10^6 Sham-MI cMSC EVs/mL (n=7)
10^7 Sham-MI cMSC EVs/mL (n=7)
10^8 Sham-MI cMSC EVs/mL (n=7)
C. cMSC-EVs from HF: LLC migration
D. cMSC-EVs from sham-MI: LLC migration
Scratch closure [%]
P= 10^7 vs 10^6 HF cMSC-EVs / mL
p=0.060
p=0.012
p=0.0005
10^7 HF cMSC EVs/mL (n=7)
10^8 HF cMSC EVs/mL (n=7)
10^6 HF cMSC EVs/mL (n=7)
Saline (n=8)
P= 10^7 vs 10^6 sham-MI cMSC-EVs / mL
p=0.3
p=0.025
10^7 Sham-MI cMSC EVs/mL (n=7)
10^6 Sham-MI cMSC EVs/mL (n=7)
10^8 Sham-MI cMSC EVs/mL (n=7)
Saline (n=8)
E. EC permeability
F. EC migration
HRP concentration [pM]
Time [h]
P for HF EVs vs Sham-MI EVs on the graph
p=0.054
p=0.061
HF EVs (n=6)
Sham-MI EVs (n=6)
Saline (n=6)
0h
16h
32h
HF cMSC EVs
Sham-MI cMSC EVs
Saline
p=0.070
p=0.033
HF EVs (n=8)
Sham-MI EVs (n=7)
Saline (n=9)

### Figure S9.

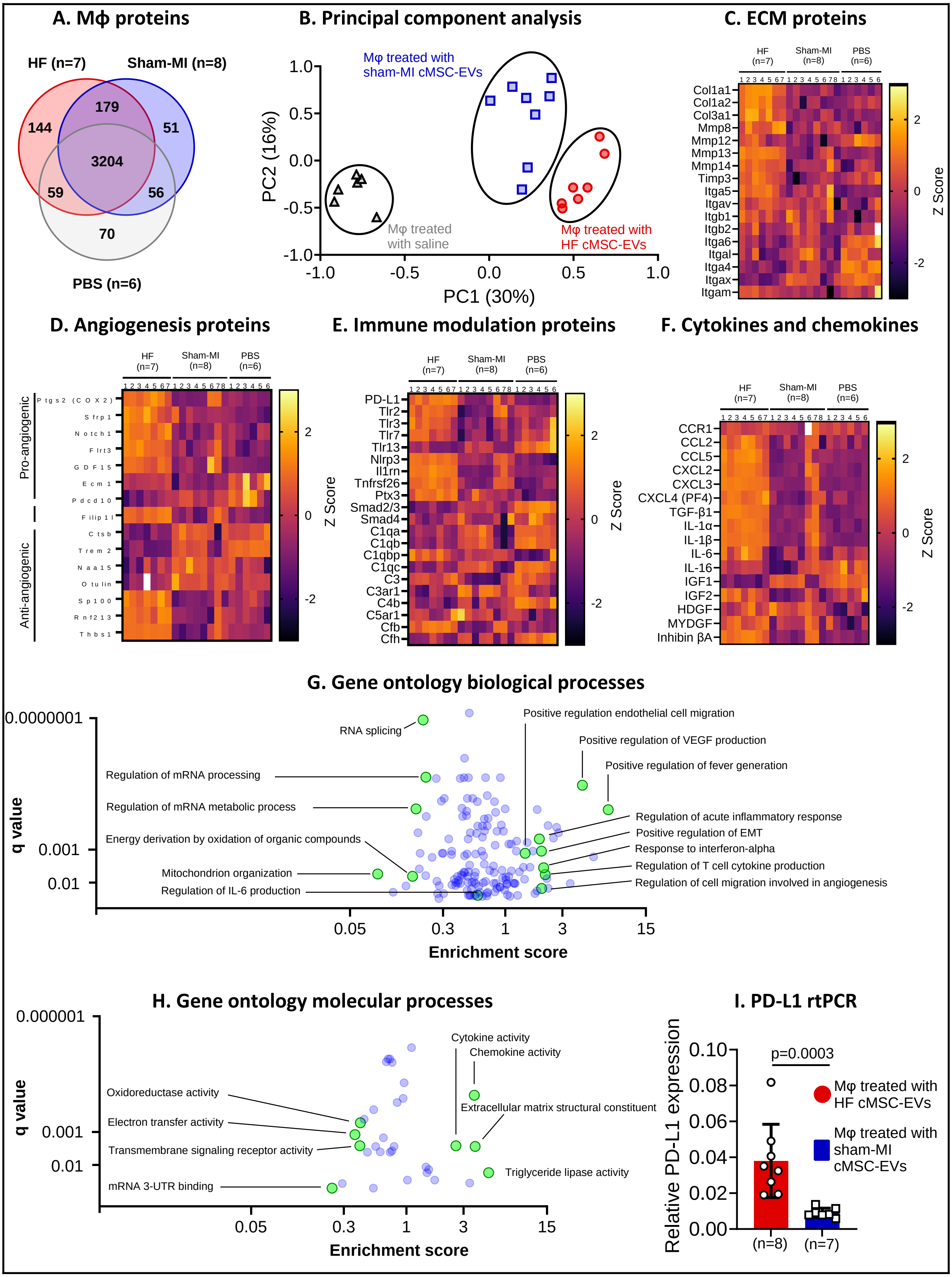

A. Mϕ proteins
HF (n=7)
Sham-MI (n=8)
PBS (n=6)
144
179
51
3204
59
56
70
B. Principal component analysis
Mφ treated with sham-MI cMSC-EVs
Mφ treated with saline
Mφ treated with HF cMSC-EVs
PC1 (30%)
PC2 (16%)
C. ECM proteins
Col1a1
Col1a2
Col3a1
Mmp8
Mmp12
Mmp13
Mmp14
Timp3
Itga5
Itgav
Itgb1
Itgb2
Itga6
Itgal
Itga4
Itgax
Itgam
Z Score
D. Angiogenesis proteins
Pro-angiogenic
Anti-angiogenic
Ptgs2 (COX2)
Sfrp1
Notch1
Flrt3
GDF15
Ecm1
Pdcd10
Filip1l
Ctsb
Trem2
Naa15
Otulin
Sp100
Rnf213
Thbs1
E. Immune modulation proteins
PD-L1
Tlr2
Tlr3
Tlr7
Tlr13
Nlrp3
Il1rn
Tnfrsf26
Ptx3
Smad2/3
Smad4
C1qa
C1qb
C1qbp
C1qc
C3
C3ar1
C4b
C5ar1
Cfb
Cfh
F. Cytokines and chemokines
CCR1
CCL2
CCL5
CXCL2
CXCL3
CXCL4 (PF4)
TGF-β1
IL-1α
IL-1β
IL-6
IL-16
IGF1
IGF2
HDGF
MYDGF
Inhibin βA
G. Gene ontology biological processes
RNA splicing
Positive regulation endothelial cell migration
Positive regulation of VEGF production
Positive regulation of fever generation
Regulation of mRNA processing
Regulation of mRNA metabolic process
Energy derivation by oxidation of organic compounds
Mitochondrion organization
Regulation of IL-6 production
Regulation of acute inflammatory response
Positive regulation of EMT
Response to interferon-alpha
Regulation of T cell cytokine production
Regulation of cell migration involved in angiogenesis
q value
Enrichment score
H. Gene ontology molecular processes
Cytokine activity
Chemokine activity
Oxidoreductase activity
Electron transfer activity
Transmembrane signaling receptor activity
mRNA 3-UTR binding
Extracellular matrix structural constituent
Triglyceride lipase activity
q value
Enrichment score
I. PD-L1 rtPCR
p=0.0003
Relative PD-L1 expression
Mφ treated with HF cMSC-EVs
Mφ treated with sham-MI cMSC-EVs
(n=8)
(n=7)

### Figure S10.

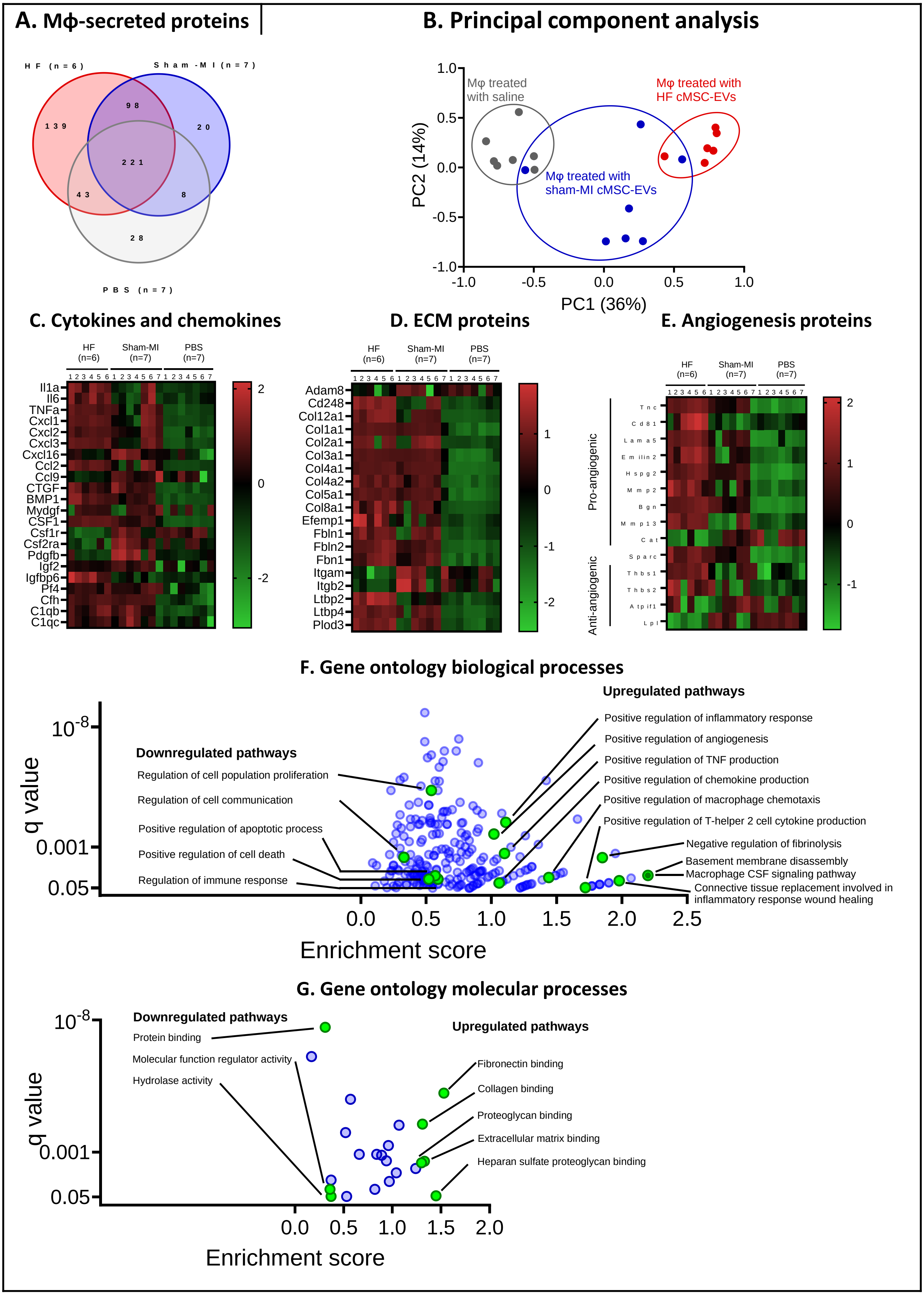
A. Mϕ-secreted proteins
HF (n=6)
Sham-MI (n=7)
PBS (n=7)
139
98
20
221
43
8
28
B. Principal component analysis
Mφ treated with saline
Mφ treated with HF cMSC-EVs
Mφ treated with sham-MI cMSC-EVs
PC2 (14%)
PC1 (36%)
1.0
0.5
0.0
-0.5
-1.0
C. Cytokines and chemokines
HF (n=6)
Sham-MI (n=7)
PBS (n=7)
Il1a
Il6
TNFa
Cxcl1
Cxcl2
Cxcl3
Cxcl16
Ccl2
Ccl9
CTGF
BMP1
Mydgf
CSF1
Csf1r
Csf2ra
Pdgfb
Igf2
Igfbp6
Pf4
Cfh
C1qb
C1qc
D. ECM proteins
Adam8
Cd248
Col12a1
Col1a1
Col2a1
Col3a1
Col4a1
Col4a2
Col5a1
Col8a1
Efemp1
Fbln1
Fbln2
Fbn1
Itgam
Itgb2
Ltbp2
Ltbp4
Plod3
E. Angiogenesis proteins
Pro-angiogenic
Anti-angiogenic
Tnc
Cd81
Lama5
Emilin2
Hspg2
Mmp2
Bgn
Mmp13
Cat
Sparc
Thbs1
Thbs2
Atpif1
Lpl
F. Gene ontology biological processes
Upregulated pathways
Positive regulation of inflammatory response
Positive regulation of angiogenesis
Positive regulation of TNF production
Positive regulation of chemokine production
Positive regulation of macrophage chemotaxis
Positive regulation of T-helper 2 cell cytokine production
Negative regulation of fibrinolysis
Basement membrane disassembly
Macrophage CSF signaling pathway
Connective tissue replacement involved in inflammatory response wound healing
Downregulated pathways
Regulation of cell population proliferation
Regulation of cell communication
Positive regulation of apoptotic process
Positive regulation of cell death
Regulation of immune response
q value
Enrichment score
G. Gene ontology molecular processes
Downregulated pathways
Protein binding
Molecular function regulator activity
Hydrolase activity
Upregulated pathways
Fibronectin binding
Collagen binding
Proteoglycan binding
Extracellular matrix binding
Heparan sulfate proteoglycan binding
q value
Enrichment score

### Figure S11.

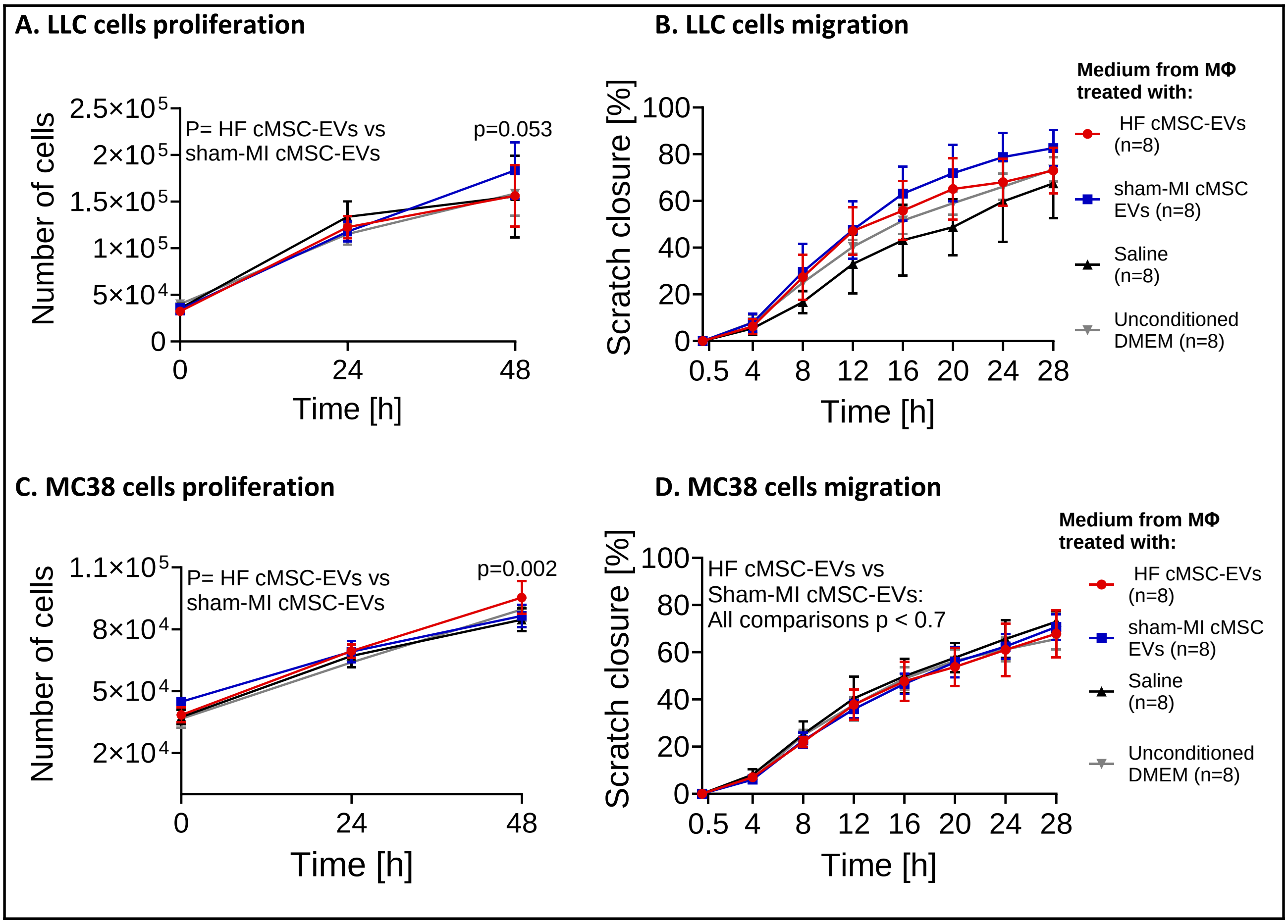
A. LLC cells proliferation
B. LLC cells migration
C. MC38 cells proliferation
D. MC38 cells migration
P= HF cMSC-EVs vs sham-MI cMSC-EVs
p=0.053
p=0.002
HF cMSC-EVs vs Sham-MI cMSC-EVs: All comparisons p < 0.7
Number of cells
Scratch closure [%]
Time [h]
Medium from MΦ treated with:
HF cMSC-EVs (n=8)
sham-MI cMSC EVs (n=8)
Saline (n=8)
Unconditioned DMEM (n=8)

**Figure S12.**

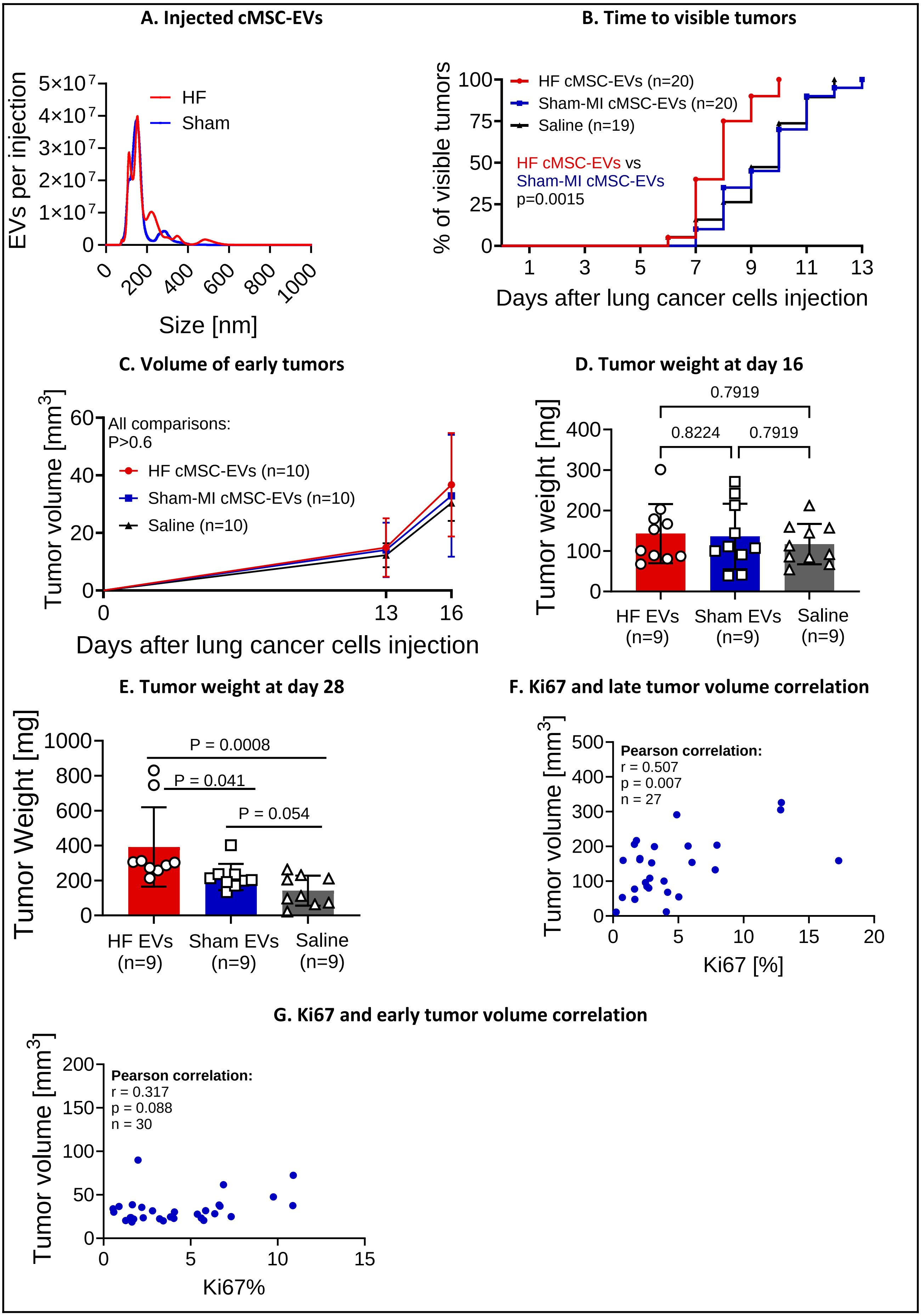
A. Injected cMSC-EVs
EVs per injection
5×10^7
4×10^7
3×10^7
2×10^7
1×10^7
0
HF
Sham
0
200
400
600
800
1000
Size [nm]
B. Time to visible tumors
% of visible tumors
100
75
50
25
0
HF cMSC-EVs (n=20)
Sham-MI cMSC-EVs (n=20)
Saline (n=19)
HF cMSC-EVs vs
Sham-MI cMSC-EVs
p=0.0015
1
3
5
7
9
11
13
Days after lung cancer cells injection
C. Volume of early tumors
Tumor volume [mm3]
60
40
20
0
All comparisons:
P>0.6
HF cMSC-EVs (n=10)
Sham-MI cMSC-EVs (n=10)
Saline (n=10)
0
13
16
Days after lung cancer cells injection
D. Tumor weight at day 16
Tumor weight [mg]
400
300
200
100
0
0.7919
0.8224
0.7919
HF EVs (n=9)
Sham EVs (n=9)
Saline (n=9)
E. Tumor weight at day 28
Tumor Weight [mg]
1000
800
600
400
200
0
P = 0.0008
P = 0.041
P = 0.054
HF EVs (n=9)
Sham EVs (n=9)
Saline (n=9)
F. Ki67 and late tumor volume correlation
Tumor volume [mm3]
500
400
300
200
100
0
Pearson correlation:
r = 0.507
p = 0.007
n = 27
0
5
10
15
20
Ki67 [%]
G. Ki67 and early tumor volume correlation
Tumor volume [mm3]
200
150
100
50
0
Pearson correlation:
r = 0.317
p = 0.088
n = 30
0
5
10
15
Ki67%

### Figure S13.

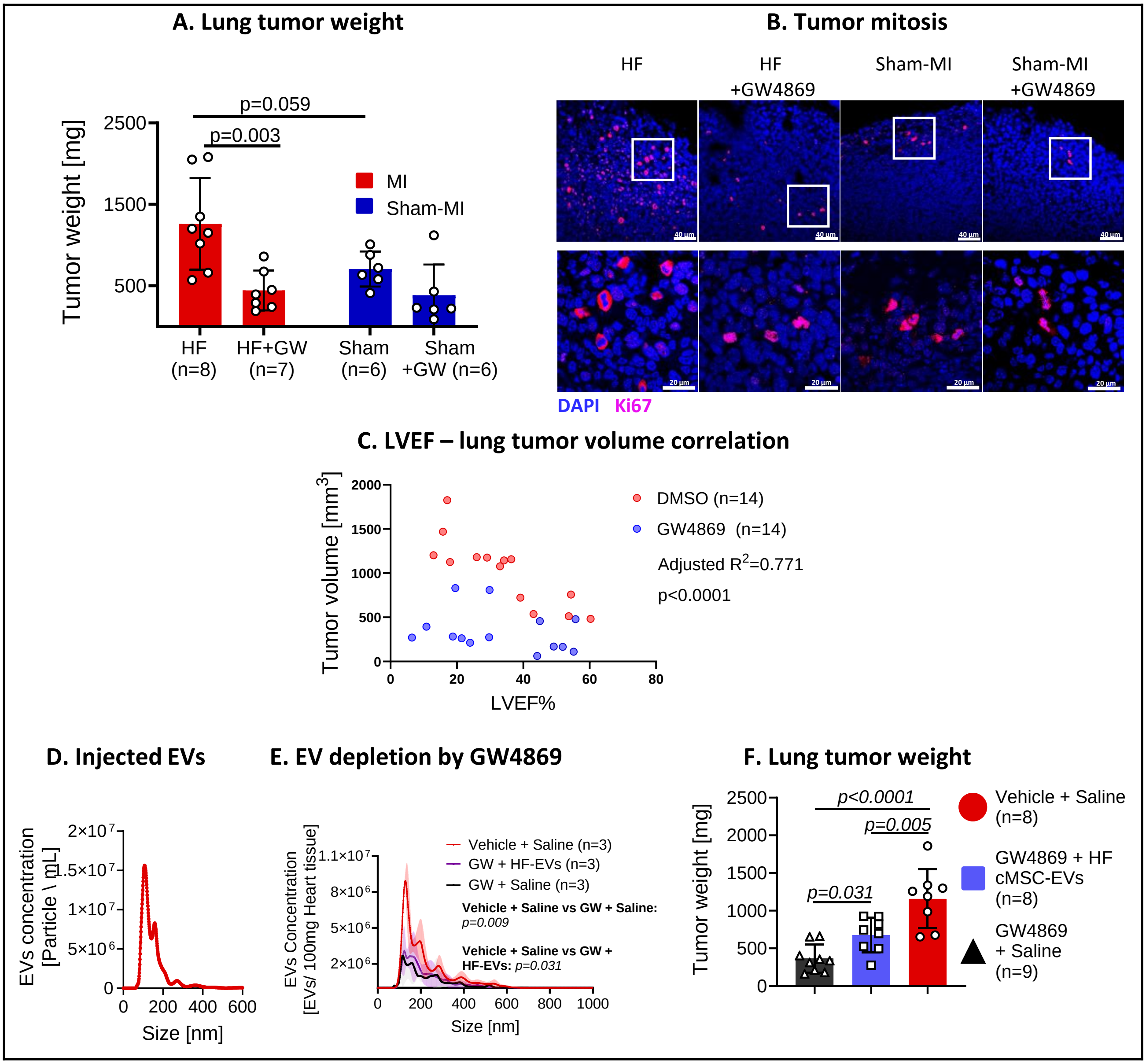

A. Lung tumor weight
Tumor weight [mg]
p=0.059
p=0.003
MI
Sham-MI
HF (n=8)
HF+GW (n=7)
Sham (n=6)
Sham +GW (n=6)
B. Tumor mitosis
HF
HF +GW4869
Sham-MI
Sham-MI +GW4869
DAPI
Ki67
C. LVEF – lung tumor volume correlation
Tumor volume [mm3]
DMSO (n=14)
GW4869 (n=14)
Adjusted R2=0.771
p<0.0001
LVEF%
D. Injected EVs
EVs concentration [Particle \ mL]
Size [nm]
E. EV depletion by GW4869
EVs Concentration [EVs/ 100mg Heart tissue]
Vehicle + Saline (n=3)
GW + HF-EVs (n=3)
GW + Saline (n=3)
Vehicle + Saline vs GW + Saline: p=0.009
Vehicle + Saline vs GW + HF-EVs: p=0.031
Size [nm]
F. Lung tumor weight
Tumor weight [mg]
p<0.0001
p=0.005
p=0.031
Vehicle + Saline (n=8)
GW4869 + HF cMSC-EVs (n=8)
GW4869 + Saline (n=9)

#### Figure S14.

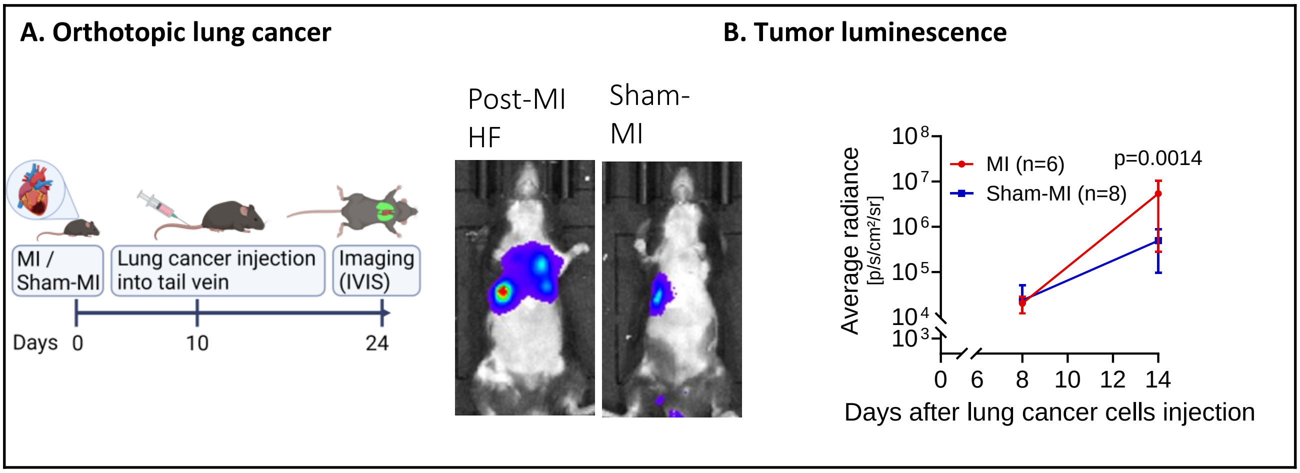

A. Orthotopic lung cancer
B. Tumor luminescence
MI / Sham-MI
Lung cancer injection into tail vein
Imaging (IVIS)
Days 0
10
24
Post-MI HF
Sham-MI
Average radiance [p/s/cm²/sr]
10^8
10^7
10^6
10^5
10^4
10^3
MI (n=6)
Sham-MI (n=8)
p=0.0014
0
6
8
10
12
14
Days after lung cancer cells injection

### Figure S15.

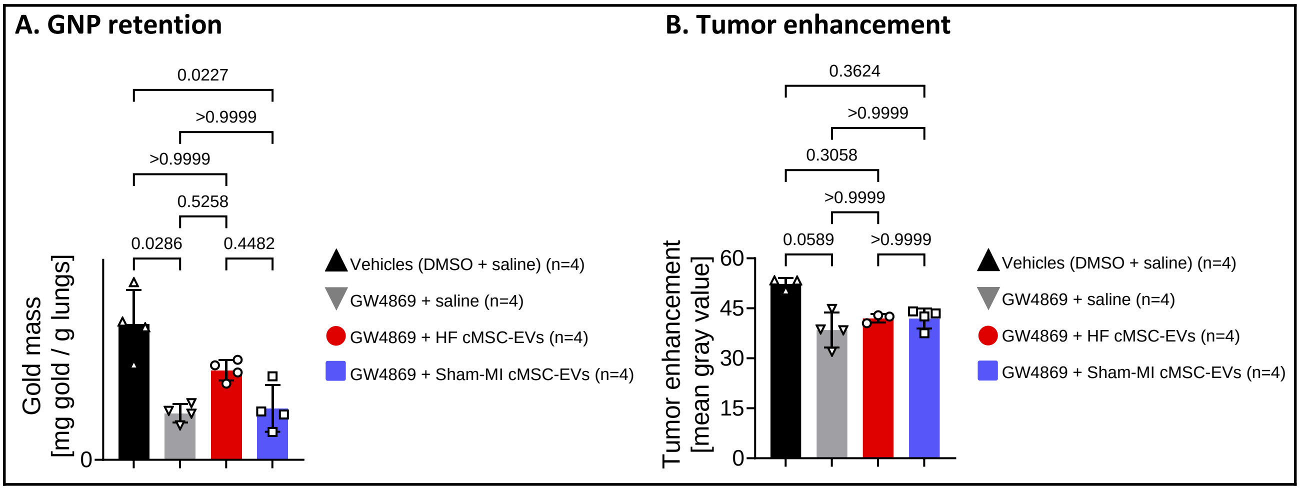
A. GNP retention
0.0227
>0.9999
>0.9999
0.5258
0.0286
0.4482
Gold mass
[mg gold / g lungs]
0
Vehicles (DMSO + saline) (n=4)
GW4869 + saline (n=4)
GW4869 + HF cMSC-EVs (n=4)
GW4869 + Sham-MI cMSC-EVs (n=4)
B. Tumor enhancement
0.3624
>0.9999
0.3058
>0.9999
0.0589
>0.9999
Tumor enhancement
[mean gray value]
60
45
30
15
0
Vehicles (DMSO + saline) (n=4)
GW4869 + saline (n=4)
GW4869 + HF cMSC-EVs (n=4)
GW4869 + Sham-MI cMSC-EVs (n=4)

#### Figure S16.

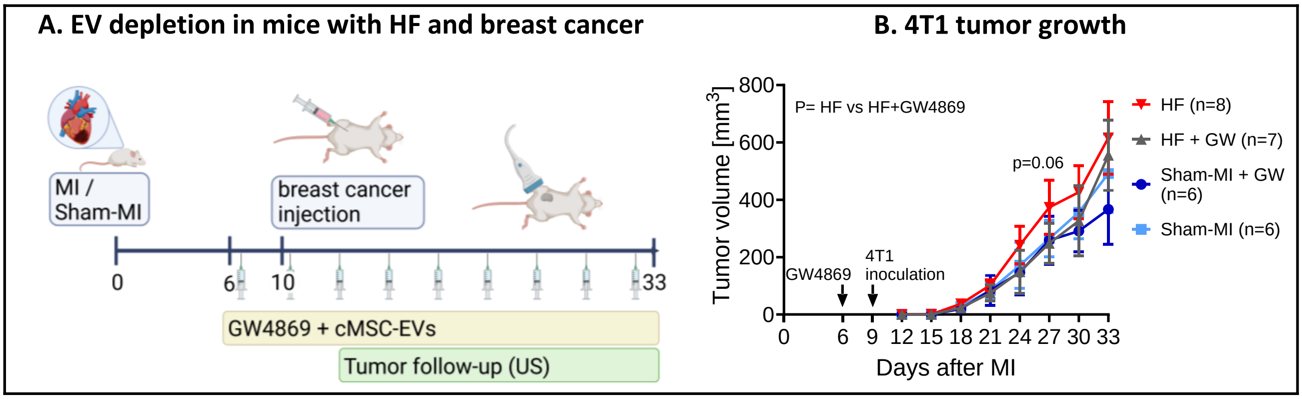
A. EV depletion in mice with HF and breast cancer
MI /
Sham-MI
breast cancer
injection
0
6
10
33
GW4869 + cMSC-EVs
Tumor follow-up (US)
B. 4T1 tumor growth
Tumor volume [mm3]
800
600
400
200
0
P= HF vs HF+GW4869
p=0.06
GW4869
4T1
inoculation
0 6 9 12 15 18 21 24 27 30 33
Days after MI
HF (n=8)
HF + GW (n=7)
Sham-MI + GW
(n=6)
Sham-MI (n=6)

### Figure S17.

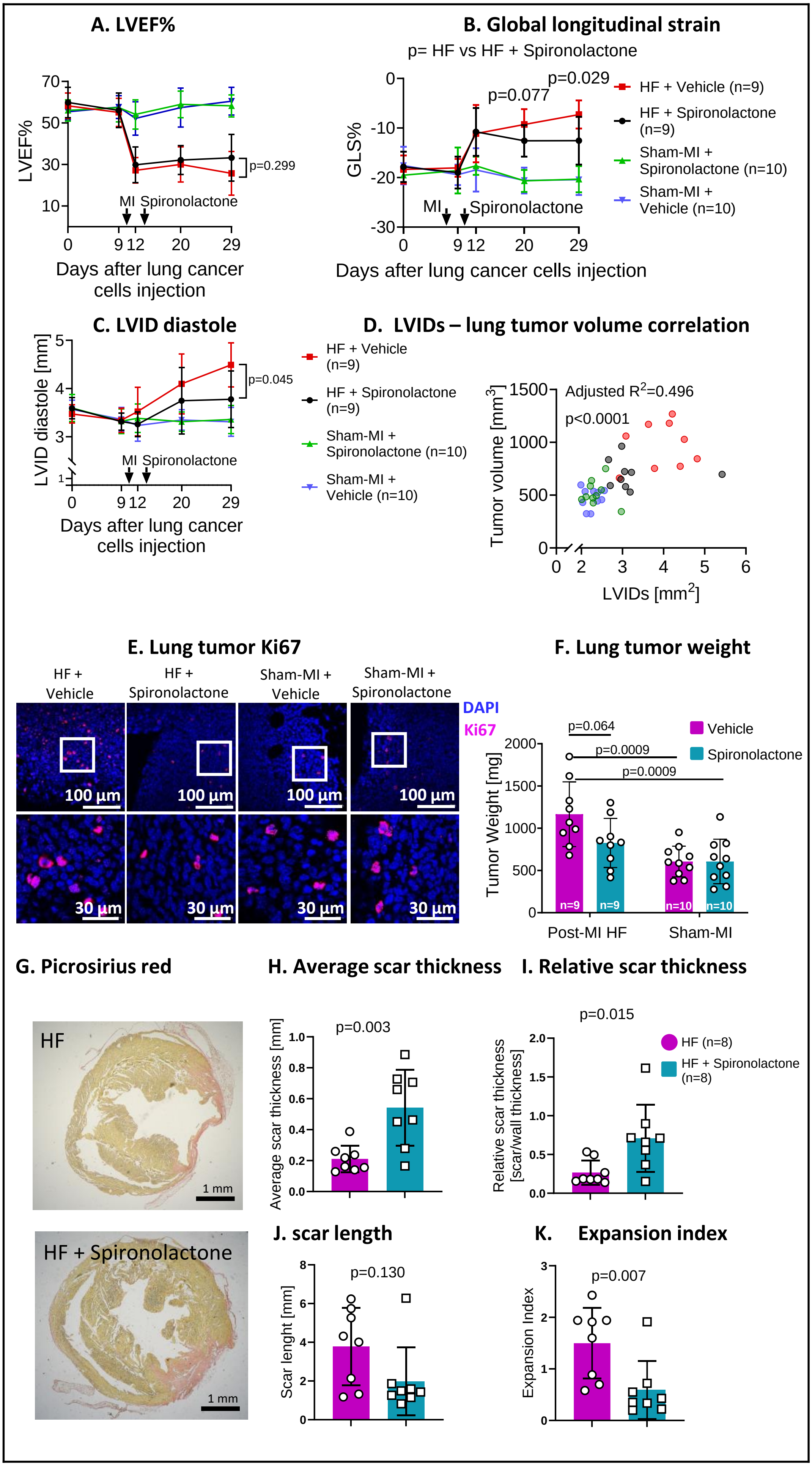

A. LVEF%
LVEF%
p=0.299
MI Spironolactone
Days after lung cancer cells injection
B. Global longitudinal strain
p= HF vs HF + Spironolactone
p=0.029
p=0.077
GLS%
MI
Spironolactone
Days after lung cancer cells injection
HF + Vehicle (n=9)
HF + Spironolactone (n=9)
Sham-MI + Spironolactone (n=10)
Sham-MI + Vehicle (n=10)
C. LVID diastole
LVID diastole [mm]
p=0.045
MI Spironolactone
Days after lung cancer cells injection
D. LVIDs – lung tumor volume correlation
HF + Vehicle (n=9)
HF + Spironolactone (n=9)
Sham-MI + Spironolactone (n=10)
Sham-MI + Vehicle (n=10)
Adjusted R2=0.496
p<0.0001
Tumor volume [mm3]
LVIDs [mm2]
E. Lung tumor Ki67
HF + Vehicle
HF + Spironolactone
Sham-MI + Vehicle
Sham-MI + Spironolactone
DAPI
Ki67
100 µm
30 µm
F. Lung tumor weight
p=0.064
p=0.0009
p=0.0009
Vehicle
Spironolactone
Tumor Weight [mg]
n=9
n=10
Post-MI HF
Sham-MI
G. Picrosirius red
HF
HF + Spironolactone
1 mm
H. Average scar thickness
p=0.003
Average scar thickness [mm]
I. Relative scar thickness
p=0.015
Relative scar thickness [scar/wall thickness]
HF (n=8)
HF + Spironolactone (n=8)
J. scar length
p=0.130
Scar lenght [mm]
K. Expansion index
p=0.007
Expansion Index

### Figure S18.

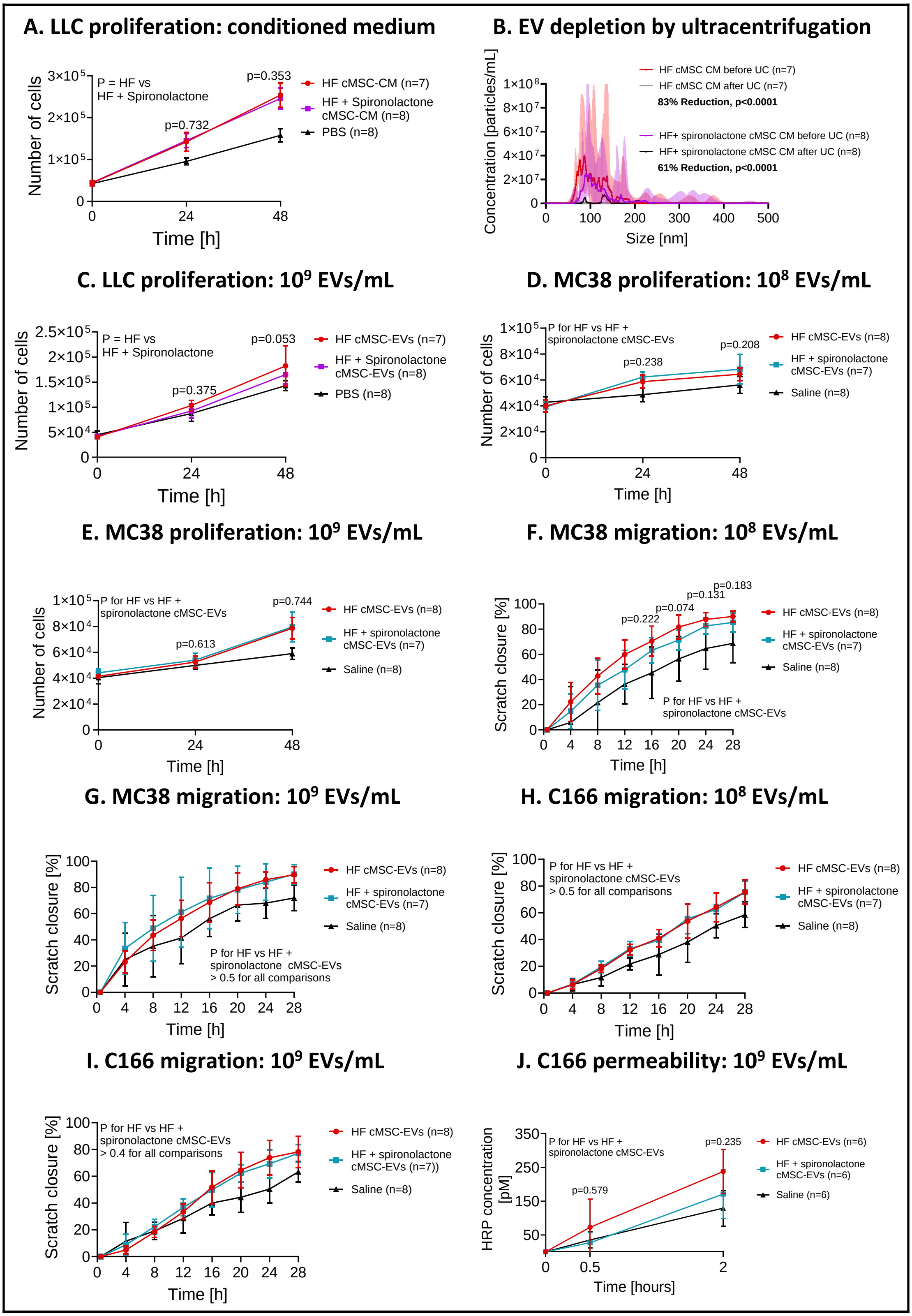

A. LLC proliferation: conditioned medium
P = HF vs HF + Spironolactone
p=0.732
p=0.353
HF cMSC-CM (n=7)
HF + Spironolactone cMSC-CM (n=8)
PBS (n=8)
Number of cells
Time [h]
B. EV depletion by ultracentrifugation
HF cMSC CM before UC (n=7)
HF cMSC CM after UC (n=7)
83% Reduction, p<0.0001
HF+ spironolactone cMSC CM before UC (n=8)
HF+ spironolactone cMSC CM after UC (n=8)
61% Reduction, p<0.0001
Concentration [particles/mL]
Size [nm]
C. LLC proliferation: 10^9 EVs/mL
P = HF vs HF + Spironolactone
p=0.375
p=0.053
HF cMSC-EVs (n=7)
HF + Spironolactone cMSC-EVs (n=8)
PBS (n=8)
Number of cells
Time [h]
D. MC38 proliferation: 10^8 EVs/mL
P for HF vs HF + spironolactone cMSC-EVs
p=0.238
p=0.208
HF cMSC-EVs (n=8)
HF + spironolactone cMSC-EVs (n=7)
Saline (n=8)
Number of cells
Time [h]
E. MC38 proliferation: 10^9 EVs/mL
P for HF vs HF + spironolactone cMSC-EVs
p=0.613
p=0.744
HF cMSC-EVs (n=8)
HF + spironolactone cMSC-EVs (n=7)
Saline (n=8)
Number of cells
Time [h]
F. MC38 migration: 10^8 EVs/mL
p=0.222
p=0.074
p=0.131
p=0.183
P for HF vs HF + spironolactone cMSC-EVs
HF cMSC-EVs (n=8)
HF + spironolactone cMSC-EVs (n=7)
Saline (n=8)
Scratch closure [%]
Time [h]
G. MC38 migration: 10^9 EVs/mL
P for HF vs HF + spironolactone cMSC-EVs > 0.5 for all comparisons
HF cMSC-EVs (n=8)
HF + spironolactone cMSC-EVs (n=7)
Saline (n=8)
Scratch closure [%]
Time [h]
H. C166 migration: 10^8 EVs/mL
P for HF vs HF + spironolactone cMSC-EVs > 0.5 for all comparisons
HF cMSC-EVs (n=8)
HF + spironolactone cMSC-EVs (n=7)
Saline (n=8)
Scratch closure [%]
Time [h]
I. C166 migration: 10^9 EVs/mL
P for HF vs HF + spironolactone cMSC-EVs > 0.4 for all comparisons
HF cMSC-EVs (n=8)
HF + spironolactone cMSC-EVs (n=7))
Saline (n=8)
Scratch closure [%]
Time [h]
J. C166 permeability: 10^9 EVs/mL
P for HF vs HF + spironolactone cMSC-EVs
p=0.579
p=0.235
HF cMSC-EVs (n=6)
HF + spironolactone cMSC-EVs (n=6)
Saline (n=6)
HRP concentration [pM]
Time [hours]

**Figure S1: Functional Properties of Crude Cardiac sEVs**

A) To isolate and analyze myocardial EVs, we subjected female C57BL/6 mice to MI or sham-MI. At day 10, the mice were euthanized, and the hearts were perfused with ice-cold PBS and then extracted and chopped on ice for EV isolation.

B) SEC successfully separated crude cEVs from free soluble proteins.

C) Typical EV morphology confirmed by transmission electron microscopy. Scale bars: 200 nm (left panel) and 100 nm (right panel).

D) The presence of typical EV markers: CD81 and TSG101 determined by a western blot.

E-F) To determine cardiac EV secretion, we used nanoparticle tracking analysis (NTA). The failing hearts generated 2.42-fold more sEVs compared to sham-operated hearts. Percentage change and p values were calculated using zero-inflated negative binomial regression.

G-H) A colorimetric proliferation assay showed that HF crude cEVs ($5 \cdot 10^8$ EVs/mL) facilitated LLC cancer cell proliferation compared to sham-MI crude cEVs. P values were calculated using two-way ANOVA with Holm-Šídák's post-test. P for post-MI HF crude cEVs $< 0.0001$, p for time $< 0.0001$, p for interaction $= 0.0015$. In addition, the effect followed a dose-response pattern. P values were calculated using two-way ANOVA with Holm-Šídák's post-test. P for post-MI HF crude cEVs $< 0.0001$, p for time $< 0.0001$, p for interaction $= 0.0018$. All samples were assayed in duplicates.

I-J) A migration (scratch) assay showed that HF crude cEVs ($5 \cdot 10^8$ EVs/mL) facilitated LLC cancer cell migration faster than sham-MI- HF crude cEVs. P values were calculated using two-way repeated measures ANOVA with Holm-Šídák's post-test. P for post-MI HF crude cEVs = 0.0015, p for time $< 0.0001$, p for interaction $< 0.0001$. In addition, the effect followed a dose-response pattern. P values were calculated using two-way repeated measures ANOVA with

Holm-Šídák’s post-test. P for post-MI HF crude cEVs = 0.0017, p for time < 0.0001, p for interaction = 0.0004. P values for HF crude cEVs vs. sham-MI- HF crude cEVs are indicated on the graphs. All samples were assayed in duplicates.

Illustration created with BioRender.com.

**Figure S2: Heart Failure Accelerated Tumor Growth.**

A) To confirm HF in mice 10 days after MI, we assessed lung congestion by hematoxylin and eosin staining of paraffin-embedded lung tissues. Compared with sham-MI, lungs after MI were characterized by distended, congested capillaries (marked by arrows), small alveoli filled with transudate (marked by *), and thickened alveolar septa, which did not occur in sham-MI-operated mice. Blood vessels marked with "v" and selected alveoli marked with "a". Scale bars: 1 mm (X2 magnification), 400 μm (X10 magnification), and 40 μm (X60 magnification).

B) Schematics for a model of heterotopic lung cancer and post-MI HF. We inoculated LLC cancer cells (750,000 in PBS) to the right hindlimb of Female C57BL/6 mice, and then at day 10, mice were randomized for either MI or sham-MI operation. We evaluated the cardiac function of operated mice by echocardiography.

C) Mice subjected to MI developed significant LV dysfunction. We measured LVEF by serial echocardiographic measurements. Two-way repeated measures ANOVA with Holm-Šídák's post-test determined P values. P for post-MI HF $< 0.0001$, p for time $< 0.0001$, p for interaction $< 0.0001$. P for post-MI HF vs. sham-MI indicated on the graph.

D-E) Mice after MI developed significantly larger tumors as measured by ultrasound (C) and confirmed by weight (D) at the end of the experiment. P values were determined by two-way repeated measures ANOVA with Holm-Šídák's post-test for ultrasound measurements (P for post-MI HF = 0.0046, p for time $< 0.0001$, p for interaction $< 0.0001$); and by two-tailed, unpaired Mann-Whitney U test.

Abbreviations: LLC - Lewis lung carcinoma. LV - left ventricle. LVEF - left ventricular ejection fraction.

Illustration created with BioRender.com.

**Figure S3: Characterization of Cardiac Mesenchymal Stromal Cells**

To characterize cMSCs, we used a viability assay and flow cytometry. We labeled isolated cMSCs for fibroblast markers: COL1α, CD90.2, mEF-SK4, PDGFRα, macrophage markers: F4/80 and endothelial cel marker: CD31.

Our gating strategy is shown for each experiment and included the exclusion of debris (FSC-A vs. SSC-A), inclusion of single cells (singlets) and not clusters of cells (FSC-A vs. FSC-H), and then gating for positive staining compared to the appropriate isotype control antibody.

A) COL1α was expressed on 94% of cMSCs.

B) CD90.2 expressed on 76% of cMSCs.

C) mEF-SK4 expressed on 98-99% of the cMSCs.

D) Expression of PDGFRα varied between MI (36%) and sham-MI (48.0%) groups, p=0.020. P values were determined by an unpaired two-tailed t-test. Normality was tested by the Shapiro-Wilk test.

E-F) The expression of F4/80 and CD31 was negligible.

G) To assess the viability of cMSCs after culture in serum-free conditions, we used trypan blue staining. The viability of the cells was 90.5% for MI-cMSCs and 86.7 for sham-MI-cMSCs. Scale bar: 200 μm.

Abbreviations: COL1α - collagen 1 alpha. FSC - forward scatter. SMA - smooth muscle actin. PDGFRα - platelets-derived growth factor receptor alpha.

**Figure S4: Images of the Untouched Membranes of Western Blots**

A) Images of the western blot membrane for crude cEVs western blots.

B) Images of the western blot membrane for cMSC-sEVs western blots.

**Figure S5: Biological Pathways and Proteomic Analysis**

A) Venn diagrams showing the overlap in protein expression between cMSC-sEVs and free proteins from either HF or sham-MI mice.

B) To further characterize the tumor-promoting cargo of cMSC-sEVs, we analyzed the differences in biological pathways enrichment between cMSC-sEVs from HF vs. sham-MI. We utilized the functional enrichment analysis tool (FunRich) of VesiclePedia using a gene ontology biological processes database. cMSC-sEVs from the failing heart were enriched in pathways related to the development and progression of cancer. For example, we identified pathways such as vascular endothelial cells proliferation (exclusively in HF-cMSC-sEVs), mitotic cell cycle phase transition (exclusively in HF-cMSC-sEVs ), and actin filament-based movement (+2.1-fold change), and negative regulation of extrinsic apoptotic pathway (+1.1-fold change). In addition, cMSC's-EVs from failing hearts downregulated expression of proteins related to negative regulation of notch signaling (-3.6-fold change), regulation of TGFβ activation (-2.2-fold change), and cellular response to tumor cells (-1.2-fold change).

**Figure S6: Soluble Cytokines from cMSCs and Additional cMSC-sEVs Cytokines**

A) To investigate additional cMSC-sEV-encapsulated cytokines, we used ELISA assays to evaluate cMSC-sEVs 10 days after MI or sham-MI. P values were determined by multiple two-tailed unpaired Mann-Whitney U tests with Holm Šídák's post-test to account for multiple comparisons between all investigated cytokines.

B). We used ELISA essays to determine the expression of soluble (not EV-encapsulated) cytokines from cMSC-conditioned medium 10 days after MI or sham-MI. P values were determined by multiple two-tailed unpaired Mann-Whitney U tests with Holm Šídák's post-test to account for multiple comparisons between all investigated cytokines.

**Figure S7: Proteomics of LLC Tumors and Potential microRNA's**

To identify potential cardiac miRs that could affect tumor growth, we analyzed the proteomic profiles of LLC tumors in search for miR binding sites.

A) We analyzed and compared the proteomics of LLC tumors from mice with failing or non-failing hearts. To analyze how HF affects protein expression in the tumor, we inoculated LLC cancer cells ($750 \cdot 10^3$) to the right hindlimb of Female C57BL/6 mice, and on day 10, they were subjected to either MI or sham-MI. On day 30, we removed the tumors and performed a proteomic analysis. The volcano plot showed a statistically significant change in 101 proteins, with marked downregulation of proteins in the HF group. P values were determined by multiple, two-tailed, unpaired t-tests with the FDR method to account for multiple comparisons.

B) To suggest potential miRs that could affect tumor proteomic profile, we used a series of filtrations demonstrated in the Venn diagram. First, we used the TargetScan tool and identified 48 of the 101 changed proteins as having a known miR binding site in their 3'-untranslated region (3'-UTR) and a total of 70 miRs that can bind to those sites. Next, to filter the miR list we generated, we used the miRBase tool to filter for miR with reported involvement in cancer. Then, we further filtered the list for miR involvement in cardiovascular diseases and found 17 potential neoplastic miRs.

Abbreviations: cMSC-EVs - cardiac mesenchymal stromal cells EVs. FDR - false discovery rate. LLC - Lewis lung carcinoma. miR - microRNA. UTR - untranslated region.

**Figure S8: Additional Functional Properties of cMSCs from Post-MI Failing Hearts**

A-B) Proliferation of LLC cancer cells increased with an increasing number of cMSC-sEVs from failing with the highest effect at the highest dose of $10^8$ EVs per mL. P values were calculated using two-way ANOVA with Holm-Šídák's post-test. P for post-MI HF cMSC-sEVs < 0.0001, p for time < 0.0001, p for interaction < 0.0001. In contrast, the proliferation of LLC cells did not increase with incremental doses of cMSC-sEVs from non-failing hearts, suggesting that the proliferative effect is associated with changes in the cargo of post-MI HF cMSC-sEVs. P values were calculated using two-way ANOVA with Holm-Šídák's post-test. P for post-sham-MI cMSC-sEVs = 0.0145, p for time < 0.0001, p for interaction = 0.0421. P for high dose ($10^8$ cMSC-EVs/mL) vs. medium dose ($10^7$ cMSC-sEVs/mL) indicated on the graphs. All samples were assayed in duplicates.

C-D) Increased concentrations of cMSC-sEVs (from 0 to $10^6$ and $10^7$ EVs per mL) from post-MI failing hearts accelerated LLC cell migration. With the highest concentrations, the migratory effect of cMSC-sEVs disappeared. P values were calculated using two-way repeated measures ANOVA with Holm-Šídák's post-test. P for post-MI HF cMSC-sEVs < 0.0001, p for time < 0.0001, p for interaction < 0.0001. We observed the same effect for cells treated with sham-MI cMSC-sEVs. However, in any concentration of EVs above 0, the migration of LLC cells was faster if they were treated with cMSC-sEVs from failing hearts, compared with cMSC-sEVs from sham-MI. P values were calculated using two-way repeated measures ANOVA with Holm-Šídák's post-test. P for sham-MI cMSC-sEVs = 0.000l7, p for time < 0.0001, p for interaction < 0.0001. P for medium dose ($10^7$ cMSC-EV/mL) vs. low dose ($10^6$ cMSC-sEVs/mL) indicated on the graphs. All samples were assayed in duplicates.

E) cMSC-sEVs from HF mice ($10^9$ EVs/mL) increased permeability of the C166 EC monolayer, compared to cMSC-sEVs from sham-MI operated mice, demonstrated by a horseradish peroxidase (HRP) transwell diffusion assay. P values were calculated using two-way repeated measures ANOVA with Holm-Šídák's post-test. P for post-MI HF cMSC-sEVs = 0.019, p for time < 0.0001, p for interaction = 0.012. P for post-MI HF vs. sham-MI is indicated on the graph. All samples were assayed in duplicates.

F) Scratch closure assay revealed that cMSC-sEVs from post-MI HF mice ($10^9$ EVs/mL) stimulated migration of the C166 EC line, compared to cMSC-sEVs from sham-MI-operated mice. P values were calculated using two-way repeated measures ANOVA with Holm-Šídák's post-test. P for HF-cMSC-sEVs = 0.024, p for time < 0.0001, p for interaction < 0.0001. P for post-MI HF vs. sham-MI is indicated on the graph. All samples were tested in duplicates.

**Figure S9: Cellular Proteome of Macrophages Educated by cMSC-sEVs**

To determine the effect of cMSC-sEVs on macrophage activation, we isolated peritoneal macrophages from female C57BL/6 mice using a resistance to trypsinization assay. Next, we incubated macrophages with cMSC-sEVs ($10^9$ EVs/mL) from the failing heart, sham-operated heart, or saline for 24 hours in serum-free conditions. Then, cells were washed and incubated for another 24 hours in a serum-free medium. After 24 hours, we lysed the macrophages. The proteome of macrophage lysate was analyzed.

A-B) Venn diagram and principal component analysis showing that macrophages activated by cMSC-sEVs from HF expressed unique proteomic profile.

C-F) Heat maps showing differential expression of cellular proteins involved in ECM remodeling (C), angiogenesis (D), immune modulation (E), and cytokines and growth factors (F).

G) Analysis of biological pathways using the STRING tool and annotation by gene ontology biological processes revealed that macrophages treated with cMSC-sEVs from post-MI HF mice are enriched in pathways that can promote cancer, such as endothelial cell migration, VEGF production, endothelial to mesenchymal transition, cell migration involved in angiogenesis, regulation of acute inflammatory response and regulation of T cell cytokine production. In addition, we found a relative depletion in several pathways related to RNA metabolism.

H) Analysis of molecular function using the STRING tool and annotation by gene ontology molecular function revealed that macrophages treated with cMSC-sEVs from post-MI HF mice are enriched in pathways that can promote cancer, such as cytokine and chemokine activity and ECM structural constituents.

I) To assess the expression of PD-L1 mRNA in cMSC-sEVs treated macrophages, we incubated macrophages with cMSC-sEVs ($10^9$ EVs/mL) from the failing heart, sham-operated heart, or saline for 24 hours in serum-free conditions. Then, cells were washed and incubated for another 24 hours in a serum-free medium. After 24 hours, we collected the macrophage-conditioned medium and isolated total macrophage RNA. We used RT-PCR to determine the expression of PD-L1 in isolated macrophages. P values were determined by the Mann-Whitney U test after the samples failed to pass the D'Agostino-Pearson normality test. P values for specific comparisons are indicated on the graph.

**Figure S10: Extracellular Proteome (Secretome) from cMSC-sEVs-Treated Macrophages**

To test the effect of cMSC-sEVs on macrophage polarization, we isolated peritoneal macrophages from female C57BL/6 mice using a resistance to trypsinization assay. Next, we incubated macrophages with cMSC-sEVs ($10^9$ EVs/mL) from the failing heart, sham-operated heart, or saline for 24 hours in serum-free conditions. Then, cells were washed and incubated for another 24 hours in a serum-free medium. After 24 hours, we collected the macrophage-conditioned medium and carried out a proteomic analysis.

A-B) Venn diagram and principal component analysis showing that cMSC-sEVs from HF activated macrophages express unique extracellular proteome.

C-E) Heat maps showing differential expression of extracellular proteins such as cytokines and chemokines (C), proteins related to ECM remodeling (D), and angiogenesis (E).

F) Analysis of biological pathways using the STRING tool and annotation by gene ontology biological processes revealed that macrophages treated with cMSC-sEVs from post-MI HF secrete proteins that are related to pathways that can promote cancer such as positive regulation of angiogenesis, chemokine production, macrophages chemotaxis, basement membrane disassembly and connective tissue remodeling. In addition, we found a relative depletion in several pathways related to regulation of cell communication, proliferation, and cell death; all are potential drivers of cancer.

G) Analysis of molecular function using the STRING tool and annotation by gene ontology molecular function revealed that macrophages treated with cMSC-sEVs from post-MI HF secrete proteins that are related to pathways that can promote cancer, such as fibronectin binding, extracellular matrix binding, and heparan sulfate proteoglycan binding.

**Figure S11: Effects of Educated Macrophage Secretome on Lung (LLC) and Colon (MC38) Cancer Cells**

A) LLC cell proliferation assay with conditioned medium from cMSC-sEVs treated macrophages (5 µg of total proteins). P values were calculated using two-way ANOVA with Holm-Šídák's post-test. P for conditioned medium from macrophages treated with post-MI HF cMSC=0.5136, p for time $< 0.0001$, p for interaction = 0.0796. P values for the conditioned medium of macrophages treated with post-MI HF cMSC-sEVs vs. sham-MI cMSC -sEVs are indicated on the graph. All samples assayed in duplicate.

B) LLC cell migration assay with conditioned medium from cMSC-sEVs treated macrophages (5 µg of total proteins). P values were calculated using two-way repeated measures ANOVA with Holm-Šídák's post-test. P for conditioned medium from macrophages treated with post-MI HF cMSC-sEVs = 0.0009, p for time $< 0.0001$, p for interaction $< 0.0001$. P values for the conditioned medium of macrophages treated with post-MI HF cMSC-sEVs vs. sham-MI cMSC -sEVs are indicated on the graph. All samples assayed in duplicates.

C) MC38 cell proliferation assay with conditioned medium from cMSC-sEVs treated macrophages (5 µg of total proteins). P values were calculated using two-way ANOVA with Holm-Šídák's post-test. P for conditioned medium from macrophages treated with post-MI HF cMSC = 0.0008, p for time $< 0.0001$, p for interaction = 0.0005. P values for the conditioned medium of macrophages treated with post-MI HF cMSC-sEVs vs. sham-MI cMSC-sEVs are indicated on the graph. All samples assayed in duplicates.

D) MC38 cell migration assay with conditioned medium from cMSC-sEVs treated macrophages (5 µg of total proteins). P values were calculated using two-way repeated measures ANOVA with Holm-Šídák's post-test. P for conditioned medium from macrophages treated with post-MI HF cMSC-sEVs = 0.5044, p for time $< 0.0001$, p for interaction = 0.5096. P values for the conditioned medium of macrophages treated with post-MI HF cMSC-sEVs vs. sham-MI cMSC-sEVs are indicated on the graph. All samples were assayed in duplicates.

**Figure S12: cMSC-sEVs from Post-MI Failing Hearts Accelerated Tumor Growth**

To dissect the effects of cMSC-sEVs from other soluble factors, we randomized mice to receive an equal amount of cMSC-sEVs from MI, sham-MI (2 µg of EVs protein), or saline every 48 hours. Each mouse received 3 subcutaneous injections to the inoculation site during the week before LLC inoculation (750,000 cells in 100 µL saline), and another 12 injections resuming 5 days after inoculation.

A) For EV dosing, we used 2 µg of EV protein and validated the amount using total EV count by NTA (E).

B) To assess if cMSC-sEVs educate the pre-cancerous environment, we measured the time from cancer cell inoculation to tumor development. We found that injection of post-MI HF cMSC-sEVs before tumor inoculation results in early tumor development compared with sham-MI EVs. P values were determined by Mantle-Cox (p log-rank = 0.0003, p for trend = 0.0027) test. We used the same test on each pair of groups to perform pairwise comparisons and adjusted for multiple comparisons by the Holm-Šídák's post-test. P value for post-MI HF vs. sham-MI cMSC-sEVs is indicated on the graph.

C) We analyzed tumors at early stages (until day 16), before differences in tumor volumes could be detected between groups. P values were calculated using two-way repeated measures ANOVA with Holm-Šídák's post-test. P for post-MI HF cMSC-sEVs = 0.6595, p for time < 0.0001, p for interaction = 0.8256.

D-E) Tumor weight at day 16 (D) and day 28 (E). P values were calculated using one-way ANOVA with Holm-Šídák's post-test for day 16 after the data passed the D'Agostino-Pearson normality test. P for day 16 = 0.6879. P for day 28 was calculated by Kruskal Wali's test with Dunn's post-test after the data failed the D'Agostino-Pearson normality test. p = 0.0010. P values for specific comparisons are indicated on the graph.

F-G) Correlation between tumor mitosis rate (Ki67%) and tumor volume. While we found a strong correlation between tumor mitosis rate and tumor volume at day 28 ($r=0.507$, $p=0.007$), this correlation was weak at day 16 ($r=0.317$, $p=0.088$), suggesting that cMSC-sEVs from post-MI HF accelerate tumor growth, at least in part, by induction of tumor cell mitosis. P values and r determined by Pearson correlation test.

**Figure S13: sEV Depletion and Transfer of cMSC-sEVs from Post-MI Hearts Affected Growth of Lung Tumors**

LLC cancer cells (750,000 cells in 100 µL saline) were inoculated to the hindlimb of mice. Ten days later, mice were randomized to MI or sham-MI operation. Starting three days after MI or sham-MI, mice from each group were further randomized to receive intraperitoneal (IP) injections of GW4869 (2.5 mg/kg) or DMSO (GW4869 vehicle as control) every 48 hours. Tumor growth and heart function were assessed by ultrasound and echocardiography.

A) Tumor weight at day 34 confirmed that EV depletion attenuated the tumor-trophic effect of HF. P values were calculated using two-way repeated measures ANOVA with Holm-Šídák's post-test. P for post-MI HF = 0.0.0547, p for GW4869 = 0.0010, p for interaction = 0.1163. P values for specific comparisons are indicated on the graph.

B) We stained the tumors for Ki67 and assessed cancer cell mitoses. We found a higher percentage of mitoses in tumors from mice with HF. EV depletion reduced the rate of mitoses. Scale bars: 100 µm (upper panels) and 30 µm (lower panels).

C) We found an inverse correlation between LV ejection fraction and tumor volume. EV depletion by GW4869 markedly reduced this correlation. β for interaction = 17.59, p= 0.0038, β for GW4869 = -1282, p<0.0001. β for LVEF = -21.38, p<0.0001. P values and adjusted $R^2$ were determined with multiple linear regression and a two-way interaction model.

D) cMSC-sEVs isolated from donor mice with MI were in the small-EVs range (<200nm).

E) We used NTA to confirm that GW4869 reduced the number of cardiac EVs in the GW4869-saline group and GW4869-EVs group (fold-changes are 0.45 and 0.57, p values are 0.009 and 0.031). Fold changes and p values were determined by zero-inflated negative binomial regression.

F) To confirm accelerated tumor growth following a transfer of HF cMSC-sEVs, we weighed the tumors at the end of the follow-up period. P-value determined by one-way ANOVA with Holm-Šídák's post-test. Normality was tested by the D'Agostino-Pearson omnibus test.

Illustration created with BioRender.com.

**Figure S14: Assessment of Tumor Burden in Orthotopic Lung Cancer Model.**

A) Schematics for a model of orthotopic lung cancer and post-MI HF. Female mice were randomized to either MI or sham-MI operation. Ten days after MI, we injected 1.5 x $10^6$ LLC cells, expressing stable luciferase, into the tail vein of mice. We evaluated LV function of operated mice using echocardiography and measured tumor growth using the IVIS Lumina LT system.

B) Post-MI HF facilitated the growth of lung cancer, measured by cancer cell luminescence. P values were determined by two-way repeated measures ANOVA with Holm-Šídák's post-test. P for post-MI HF = 0.0181, p for time = 0.0067, p for interaction = 0.0177.

**Figure S15: cMSC-sEVs from Post-MI Failing Hearts Promoted Retention of GF-GNPs in the Lungs after MI and Orthotopic Lung Cancer Growth**

To assess the role of cMSC-sEVs in the colonization and growth of lung tumors, we used an orthotopic lung cancer model, EV depletion by GW4869, and cMSC-sEVs transfer from either post-MI HF or sham-MI mice. We induced post-MI HF in female C57BL/6 mice, and 3 days after MI, we randomized the tumor-bearing mice into 4 treatment groups: GW4869 (intraperitoneal, 2.5 mg/kg), and then 10 µg of cMSC-sEVs from the failing heart, IP (n=9) every 48 hours; 2) GW4869 (intraperitoneal, 2.5 mg/kg), and then 10 µg of cMSC-sEVs from sham-MI, IP (n=9) every 48 hours; 3) GW4869 (intraperitoneal, 2.5 mg/kg) and saline (n=9) every 48 hours; or 4) DMSO (the vehicle of GW4869 as control) and saline (n=8) every 48 hours. Treatment with GW4869 started on day 3, and cMSC-sEV transfer started on day 4 to avoid mixing cMSC-sEVs with the GW4869 solution containing DMSO. Then, we injected luciferase-expressing LLC cells (750,000 in 100 µL PBS) into the tail vein on day 10 after MI. We scanned the lungs, with and without injection of GF-GNPs, at day 30 using micro-CT and determined the amount of GF-GNPs using inductively coupled plasma spectrometry (ICP).

A) While systemic EV depletion reduced the retention of GNPs in the lungs, exogenous administration of cMSC-sEVs from post-MI HF mice partially restored it.

B) GNPs are electron-dense. Thus, high retention in a specific area would increase the intensity of CT images. To determine if cMSC-sEVs from post-MI HF promoted the high GNPs retention in the lungs by either increasing its retention by tumor cells or by increased growth of the tumors, we analyzed the CT intensity of the tumor masses in the lungs. GW4869 reduced the retention of GNPs in the tumor tissue in the lungs, and cMSC-sEVs did not restore it. Thus, the high content of the gold signal in the lungs of mice treated with post-MI cMSC-sEVs is most likely the result of increased tumor burden and not increased retention by individual tumor cells.

**Figure S16: sEV Depletion and Transfer of cMSC-sEVs from Post-MI Hearts Modestly Affects Triple-Negative Breast Cancer Tumors**

A) We used an orthotopic model of triple negative breast cancer to assess the effects of post-MI HF and EV depletion in more malignant tumors. Female Balb/C mice were randomized for either MI of sham-MI. At day 6, the mice were further randomized to receive GW4869 (2.5 mg/kg) or its vehicle (DMSO) every 72 hours. Then, at day 10, 4T1 cancer cells (250,000 cells in 100 µL saline) were inoculated to the mammary pad of the mice. Tumor growth and heart function were assessed by ultrasound and echocardiography.

B) EV depletion and post-MI HF had a minor effect on growth of triple-negative breast cancer tumors. P values were calculated using two-way repeated measures ANOVA with Holm-Šídák's post-test. P for post-MI HF = 0.0209, p for time < 0.0001, p for interaction < 0.0001. P values for HF vs HF+GW4869 are indicated on the graph.

**Figure S17: Post-MI Spironolactone Reduced Left Ventricular Remodeling and Suppressed Tumor Growth**

We inoculated LLC cancer cells (750,000 in 100 µL PBS) in the hind limb of female C57BL/6 mice and randomized the mice for either MI or sham-MI 10 days later. Three days after MI, we started spironolactone treatment (50 mg/kg, daily) or its vehicle, injected subcutaneously. Finally, we performed serial echocardiographic and ultrasound measurements to assess cardiac remodeling, LV function, and tumor growth.

A-C) Mice subjected to MI developed significant LV dysfunction and adverse LV remodeling. Spironolactone ameliorated LV remodeling, attenuated the increase in LV diameter, and improved global longitudinal strain. P values were determined by two-way repeated measures ANOVA with Holm-Šídák's post-test. For LVEF: P for spironolactone $< 0.0001$, p for time $<$ 0.0001, p for interaction $< 0.0001$. P values for specific comparison are indicated on the graph. For global longitudinal strain: P for spironolactone = 0.0012, p for time $< 0.0001$, p for interaction $< 0.0001$. P values for HF vs. HF + spironolactone are indicated on the graph. For left ventricular internal diameter during diastole. P values for specific comparison are indicated on the graph.

D) To determine if the anti-tumor effects of spironolactone were correlated with the reduction in cardiac remodeling, we performed multiple linear regression with a two-way interaction model for spironolactone and left ventricular internal diameter during systole (LVIDs). Under spironolactone treatment, LVIDs were correlated with tumor volume (adjusted $R^2$=0.496, n=38, $p < 0.0001$, β for interaction = 177.6, p= 0.014, β for spironolactone = -458.1, p = 0.032. β for LVIDs = 65.3, p = 0.231).

E) To assess the effects of post-MI spironolactone on tumor cell mitoses, we stained the tumors for Ki67 and assessed tumor cell proliferation. We found that spironolactone reduced the number of proliferating cells in tumors from mice with HF but had no effect on the tumors from sham-operated mice. Scale bar: 100 μm (upper panels) and 30 μm (lower panels).

F) While MI accelerated the growth of heterotopic lung cancer, spironolactone significantly reduced the effect of MI and HF on tumor growth. To confirm the reduction in tumor volume, we weighed the tumors at the end of the follow-up period. Mice treated with spironolactone developed smaller tumors than mice treated with the vehicle. Notably, spironolactone did not affect the tumors of sham-operated mice. P values were determined by two-way ANOVA with Holm-Šídák's post-test. P for spironolactone = 0.0002, p for HF = 0.0761, p for interaction = 0.0754. P values for specific comparison are indicated on the graph.

G) To confirm that spironolactone successfully reduced adverse LV remodeling, we evaluated LV morphology on day 29. Representative images of the heart, stained with picrosirius red, show decreased remodeling and improved scar characteristics under spironolactone treatment. Scale bar: 1 mm.

H-K) LV morphometry indicated improved average scar thickness, relative scar thickness, and decreased scar length and expansion index. Average scar thickness was calculated from 3 measurements at the end and middle of the scar, average wall thickness from 3 measurements of septum thickness at the anterior, middle, and inferior areas, LV muscle area, LV cavity area, whole LV area, epicardial scar length, and endocardial scar length. Relative scar thickness was calculated as average scar thickness divided by average wall thickness. The expansion index was calculated as [LV cavity area/whole LV area]/relative scar thickness. P values determined by Mann Whitney U test.

Abbreviations: LV - left ventricle.

**Figure S18: Effect of Post-MI Spironolactone on cMSC-sEV-Mediated Proliferation and Migration of Cancer and Endothelial Cells**

To test the hypothesis that HF therapy will reduce the effects of cMSC-sEVs on tumor growth, we subjected female C57BL/6 mice to MI and started spironolactone treatment by SC injection (50 mg/kg, daily) three days later. On day 10, we harvested the hearts and isolated cMSCs. Then, we used cMSC conditioned medium, purified sEVs from the cMSC-conditioned medium or EV-depleted conditioned medium from cMSCs to test the effects of spironolactone on the pro-tumorigenic properties of cMSCs.

A) To test whether post-MI spironolactone treatment will reduce the ability of cMSC to promote cancer cell proliferation, we used the LLC lung cancer. Spironolactone did not affect the ability of cMSC conditioned medium (5 μg of total proteins) to promote lung cancer proliferation. P values were determined by two-way ANOVA with Holm-Šídák's post-test. P values for specific comparison are indicated on the graph. P for spironolactone $< 0.0001$, p for time $< 0.0001$, p for interaction $< 0.0001$.

B) We confirmed the depletion of sEVs from cMSC-conditioned medium by ultracentrifugation. We reduced the number of sEVs in the conditioned medium by 61-83 percent. P values and percentage change were determined by zero-inflated negative binomial regression.

C-G) To test whether post-MI spironolactone treatment will reduce the ability of cMSC-sEVs to promote cancer cell proliferation and migration, we used the LLC lung cancer and MC38 colon cancer cells. Spironolactone had minor effects on lung cancer proliferation (C), colon cancer proliferation (D-E), and colon cancer migration (F-G). P values for proliferation assays were determined by two-way ANOVA with Holm-Šídák's post-test. P values for migration assays were determined by two-way repeated measures ANOVA with Holm-Šídák's post-test. P values for specific comparison are indicated on the graph. For LLC proliferation with $10^9$ cMSC-sEVs/mL

(C): p for spironolactone = 0.0056, p for time < 0.0001, p for interaction = 0.0166. For MC38 proliferation with $10^8$ (medium dose) cMSC-sEVs/mL (D): p for spironolactone = 0.0003, p for time < 0.0001, p for interaction = 0.0009. For MC38 proliferation with $10^9$ (high dose) cMSC-sEVs/mL (E): p for spironolactone < 0.0001, p for time < 0.0001, p for interaction < 0.0001. For MC38 migration with $10^8$ (medium dose) cMSC-sEVs/mL (F): p for spironolactone = 0.0054, p for time < 0.0001, p for interaction = 0.1324. For MC38 migration with $10^9$ (high dose) cMSC-sEVs/mL (G): p for spironolactone = 0.0992, p for time < 0.0001, p for interaction = 0.1364.

H-I) Next, we tested if post-MI spironolactone treatment will reduce the ability of cMSC-sEVs to promote endothelial cell migration and permeability. While spironolactone did not affect EC C166 migration, it reduced EC C166 permeability. P values for migration and permeability assays were determined by two-way repeated measures ANOVA with Holm-Šídák's post-test. P values for specific comparison are indicated on the graph. For C166 migration with $10^8$ (medium dose) cMSC-sEVs/mL (H): p for spironolactone = 0.0025, p for time < 0.0001, p for interaction = 0.0002. For C166 migration with $10^9$ (high dose) cMSC-sEVs/mL (I): p for spironolactone = 0.0072, p for time < 0.0001, p for interaction < 0.0001. For C166 permeability (J) with $10^9$ (high dose) cMSC-sEVs/mL: p for spironolactone = 0.0352, p for time < 0.0001, p for interaction = 0.0750.
